# Integrated Pangenomic and Systems Biology Analyses Reveal the Genomic Basis of Virulence and Adaptation in *Bipolaris sorokiniana*

**DOI:** 10.64898/2026.07.31.741966

**Authors:** Anand Kumar Shukla, Narendra Kadoo

## Abstract

*Bipolaris sorokiniana* is a hemibiotrophic fungal pathogen responsible for foliar and root diseases of cereals, causing annual yield losses of 10–50%. The recurrent breakdown of host resistance and the emergence of fungicide-resistant pathogen populations underscore the urgent need to understand the genomic mechanisms underpinning pathogen adaptation and virulence. Here, we present the first comprehensive species-wide pangenomic and systems-level analyses of *B. sorokiniana* based on 19 genomes of globally distributed strains. Orthology-based analyses revealed an open pangenome comprising 16,981 orthogroups, partitioned into a conserved core genome (60.8%) and a highly dynamic accessory genome (39.2%), consisting of soft-core (8.5%), shell (19.7%), and cloud (11.0%) compartments. The core genes were predominantly associated with essential cellular and metabolic functions, while the accessory fractions were enriched in regulatory, stress-responsive, and adaptive processes. Secondary metabolite profiling identified 39–54 biosynthetic gene clusters per genome and revealed a largely conserved metabolic repertoire. Gene family evolution analyses revealed an excess of gene loss over expansion, indicating ongoing genome streamlining and lineage-specific adaptation. The core interactome comprised four densely connected functional communities governing genome maintenance, ribosome biogenesis, cellular bioenergetics, and protein translation. Collectively, this study elevates *B. sorokiniana* research from single-genome analyses to a population-scale analysis, providing vital insights into the evolutionary architecture of pathogenicity, adaptation, and genome diversification. These findings provide a valuable genomic resource for disease surveillance and functional characterization of virulence determinants, as well as the development of durable resistance strategies and next-generation antifungals for sustainable disease management in cereals.

## 1. Introduction

*Bipolaris sorokiniana* (teleomorph *Cochliobolus sativus*) is a highly adaptable, heterothallic, hemibiotrophic fungal pathogen responsible for devastating cereal diseases, including spot blotch, common root rot, crown rot, seedling blight, and black point of wheat (*Triticum* spp.) and barley (*Hordeum vulgare*) (Al-Sadi 2021; Farzana et al. 2025). Globally, spot blotch affects about 25 million hectares (mha) of wheat cultivation, with a high-intensity 10 mha epidemic hotspot in the warm and humid Eastern Gangetic Plains of South Asia, spanning Bangladesh, India, Nepal, and Pakistan (Roy et al. 2023). Driven by climate change, the disease incidence has recently expanded even into cooler, non-traditional environments such as the Kashmir Valley (Aggarwal et al. 2025). While the pathogen thrives optimally in temperatures between 18°C and 32°C, it exhibits exceptional stress tolerance. It has been reported to overwinter on crop residues or seeds for up to four years under sub-zero conditions in North America and Northern China, and can also tolerate high temperature and drought via thick-walled conidia (Viani et al. 2017). Consequently, *B. sorokiniana* inflicts severe economic damage, causing an average annual yield reduction of 17%, with localized field losses between 10% and 50%, and up to 100% during severe epidemics (Shukla et al. 2025).

*Bipolaris sorokiniana* exhibits remarkable ecological adaptability reflected in its broad host range, comprising more than 45 grass and cereal genera, including *Triticum*, *Hordeum*, *Avena*, *Secale*, *Sorghum*, and *Zea*. Although the species lacks recognized *formae speciales*, *B. sorokiniana* isolates frequently differ in aggressiveness and virulence across host genotypes (Al-Sadi 2021). This wide-ranging pathogenic lifestyle and intraspecific variation in aggressiveness are due to coordinated molecular responses comprising proteinaceous effectors, specialized carbohydrate-active enzymes (CAZymes), and highly evolved secondary metabolite biosynthetic gene clusters (BGCs) (Walton 1994; Ye et al. 2019). A critical determinant of *B. sorokiniana* virulence is the horizontally acquired necrotrophic effector *ToxA*, which triggers severe chlorotic and necrotic responses via an inverse gene-for-gene interaction with the host susceptibility locus *Tsn1* (McDonald et al. 2018). This effector operates synergistically with low-molecular-weight phytotoxins, such as helminthosporol, sorokinianin, bipolaroxin, and prehelminthosporol. These toxins are synthesized via a specialized terpene and non-ribosomal peptide synthetase (NRPS) biosynthetic pathway and disrupt host organelle oxidative phosphorylation prior to cellular colonization (Apoga et al. 2002; Jansson and Åkesson 2003).

Managing disease outbreaks requires an integrated approach that combines cultural practices, biological suppression, and targeted chemical interventions. The primary seedborne inoculum is usually managed using carboxin or thiram seed treatments, while foliar sprays of triazoles (propiconazole, tebuconazole, epoxiconazole) and strobilurins (azoxystrobin) are applied to control the disease in a standing crop (Roy et al. 2023; Basak et al. 2025). Indiscriminate use of these chemical fungicides can lead to the emergence of fungicide-resistant pathogen strains that are very difficult to control. However, chemical control remains the primary line of defense in the field. To identify the genetic factors underlying this adaptability, extensive phylogeographic and population analyses have been conducted. Transcriptomic profiling under sublethal propiconazole stress revealed rapid adaptive overexpression of sterol biosynthesis genes in *B. sorokiniana* (Somani et al. 2019). Similarly, multi-locus sequence typing (ITS, GAPDH, and TEF-1α) of 254 global isolates from 18 countries identified 40 distinct haplotypes, with H1, a globally dispersed haplotype, dominating. This haplotype accounted for ∼50% of the global pathogen population, indicating high gene flow and weak global geographic structure (Sharma et al. 2022). Likewise, ITS sequencing of 528 Indian isolates across five wheat-growing zones identified two major lineages and 40 haplotypes, with H1 remaining highly predominant at 71.4% and H2 accounting for 9.28% (Kashyap et al. 2022). This phenotypic heterogeneity is largely caused by heterokaryosis, hyphal anastomosis, and parasexual recombination, which continuously generate novel virulence combinations (Gupta et al. 2018).

Traditional population genetics and diversity studies, restricted to single-gene markers or isolated phenotypic assays, fail to capture genome-wide structural variation, dynamics of the accessory gene pool, or the spatial organization of pathogenicity loci. To fully decipher these genomic determinants of lineage-specific adaptation and host-pathogen interactions, a comprehensive pangenome framework is required. By capturing a species’ entire gene repertoire, a pangenome overcomes the limitations of a single reference genome and divides the genome into a conserved core shared by all isolates and a variable accessory genome present only in some strains (Perrier and Barber 2024). In fungal pathogens, these accessory genes exhibit frequent presence–absence variation and are heavily enriched for effectors, secondary metabolite clusters, and transposable-element-associated regions that drive phenotypic variation, virulence, host specialization, and antifungal resistance (Badet et al. 2020; Perrier and Barber 2024). Despite the power of this approach, current genomic studies of *B. sorokiniana* still rely heavily on single reference genomes, thereby limiting the detection of this vital intraspecific genetic diversity. Consequently, although specific virulence determinants such as effectors, phytotoxin biosynthetic genes, and *ToxA* have been identified in *B. sorokiniana*, their pangenome-wide distribution and evolution remain poorly understood. Furthermore, the lack of an integrated framework combining pangenomics, protein interaction networks, and adaptive selection analyses continues to hamper our understanding of how the core and accessory genomes cooperatively drive virulence, host adaptation, and genome evolution in this pathogen.

Therefore, we constructed the first comprehensive pangenome of 19 globally distributed *B. sorokiniana* strains to investigate species-wide genomic diversity, evolutionary dynamics, and pathogenicity **(Table S1)**. We characterized pangenome architecture, functionally annotated pathogenicity-associated genes, analyzed gene family evolution, secondary metabolite BGCs, and positively selected orthogroups, reconstructed protein–protein interaction networks, and integrated these datasets to reveal the molecular basis of virulence evolution, host adaptation, and genome diversification. These findings provide a valuable resource for functional genomics and support the development of durable host resistance and next-generation disease management strategies.

## 2. Materials and methods

### 2.1 Genome dataset

A total of 19 *B. sorokiniana* genomes were included in this study for comparative pangenome analysis. Eight publicly available genomes, namely BRIP10943a, BRIP27492a, BS112, LK93, ND90Pr, ND93-1, Shoemaker, and Yt-6, were retrieved from the NCBI Genome database (accessed August 1, 2025). In addition, three genomes (D2, SI, and BS52) reported by Yadav et al. (2022) and eight genomes (WAI2411, WAI2431, WAI2432, WAI3285, WAI3295, WAI3382, WAI3384, and WAI3398) reported by Bucknell et al. (2025) were incorporated into the dataset. Raw sequencing reads for these 19 strains were subjected to quality assessment using FastQC (v0.12.1) to evaluate overall sequencing quality, base quality distribution, GC content, and adapter contamination. High-quality reads were subsequently assembled *de novo* using the SPAdes Genome Assembler (v4.2.0) with the --isolate parameter enabled to optimize assembly performance for haploid fungal isolates (Prjibelski et al. 2020). Contigs shorter than 500 bp were removed from the final assemblies to improve assembly quality, reduce fragmentation, and facilitate downstream comparative genomic and pangenome analyses.

### 2.2 Genome quality assessment

The assembly quality of all 19 *B. sorokiniana* genomes was evaluated using QUAST (v5.3.0) to obtain standard assembly statistics, including total assembly size, GC content, N50, and L50 values (Gurevich et al. 2013). To further assess genome completeness, we employed BUSCO (Benchmarking Universal Single-Copy Orthologs, v6.0.0) with the dothideomycetes_odb10 lineage dataset run in euk_genome (creation date: 2024-01-08; 3,786 orthologs across 45 reference genomes) (Simão et al. 2015). This combined assessment provided both assembly-level and gene content-based quality metrics, ensuring reliability for downstream pangenome analysis.

### 2.3 Repeat identification and masking

Repeat annotation was conducted using a hybrid strategy that integrated homology-based and *de novo* approaches to comprehensively detect repetitive elements. In the first step, homology-based masking was performed with RepeatMasker (v4.1.9) (Chen 2004), utilizing the HMM-DFam database (v3.9) (Wheeler et al. 2012). This approach enabled the identification and masking of conserved repetitive elements by comparison with a curated library of known repeat families, ensuring consistent detection across all 19 genomes. Subsequently, a *de novo* repeat discovery was performed to capture novel or lineage-specific repetitive elements absent from existing databases. For this, RepeatModeler (v2.0.7) was used to construct genome-specific repeat libraries for each strain (Flynn et al. 2020). These custom libraries were then integrated into RepeatMasker for a second round of annotation, allowing identification and masking of previously uncharacterized repetitive elements.

### 2.4 Gene prediction and annotation

Gene prediction and annotation were performed using MAKER (v3.1.4) (Cantarel et al. 2008), which integrates both evidence-based and *ab initio* approaches to produce high-confidence gene models. For the evidence-based component, transcriptomic and protein homology evidence were incorporated. A total of 14 RNA-seq datasets were used: 12 from the *B. sorokiniana* D2 strain (Somani et al. 2019), and two RNA-seq datasets from the LK93 strain obtained from NCBI (Zhang et al. 2023). Raw RNA-seq reads were subjected to quality control and adapter trimming, followed by *de novo* transcriptome assembly using rnaSPAdes (v3.15.5) (Bushmanova et al. 2019). The resulting assemblies were merged and optimized using the EvidentialGene pipeline (http://arthropods.eugenes.org/EvidentialGene/), which reduces redundancy and selects the most biologically representative transcripts. In addition, protein homology evidence was compiled from NCBI RefSeq entries of five closely related species: *B. maydis*, *B. oryzae*, *B. zeicola*, *B. victoriae*, and *B. sorokiniana*. For the *ab initio* gene prediction, two widely used predictors, SNAP (v2006-07-28) (Korf 2004) and AUGUSTUS (v3.5.0) (Stanke and Morgenstern 2005), were integrated within MAKER. High-confidence gene models from the initial MAKER run were used to iteratively train both SNAP and AUGUSTUS in two successive rounds. After each training round, gene models were filtered to retain only those with an Annotation Edit Distance (AED) < 0.25, ensuring reliability of the training set. In the final annotation step, gene models with an AED < 0.50 were retained as the high-confidence set, balancing sensitivity and precision in gene discovery (Huang et al. 2025).

### 2.5 Orthology-based pangenome construction

The gene-based pangenome of *B. sorokiniana* was constructed using predicted protein sequences from all 19 genomes. Orthology inference was performed with OrthoFinder v2.5.5 (Emms and Kelly 2019), executed with DIAMOND (v2.1.13.167) for rapid all-versus-all protein similarity searches, MCL for orthogroup clustering, and MAFFT for multiple sequence alignment, followed by phylogenetic inference with FastTree (-S diamond -M msa -A mafft -T fasttree) (Katoh and Standley 2013; Buchfink et al. 2015). For pangenome partitioning, orthogroups were classified into four categories. Orthogroups shared by all 19 genomes were defined as the core genome, representing the most conserved genetic repertoire. Orthogroups present in 17–18 genomes were designated as soft-core genes, while those present in 2–16 genomes constituted the shell genome. Finally, orthogroups found in only one genome were considered a strain-specific cloud genome. To assess pangenome expansion and core genome conservation, 200 random genome permutations were performed, and cumulative pan- and core-orthogroup counts were used to estimate mean values and 95% confidence intervals. Pangenome and core genome dynamics were modeled using a Heaps’ law power function and an exponential decay function, respectively, to estimate pangenome openness and core genome reduction with increasing genome sampling (Guerra 2026).

### 2.6 Sequence-based functional annotation

For functional annotation, the longest representative protein sequence from each orthogroup was selected for downstream analyses. Sequence-based annotation was initially performed separately for each pangenome category (core, softcore, shell, and cloud orthogroups) using eggNOG-mapper (2.1.13) (Cantalapiedra et al. 2021; Hernández-Plaza et al. 2023) with DIAMOND-based searches in fast sensitivity mode. To further improve annotation coverage, representative protein sequences were searched against a clustered non-redundant (NR) protein database comprising 465,517,121 protein sequences (downloaded on September 26, 2025) using DIAMOND (v2.1.13.167) in BLASTP mode with an E-value cutoff of 1 × 10⁻⁵. Results were filtered to retain hits with ≥40% sequence identity, after which the top hit based on the highest bit score was selected for downstream annotation.

Protein domain and conserved motif annotation were performed using InterProScan (v5.75-106.0) (Jones et al. 2014). GO annotations, InterPro identifiers, and pathway information were retrieved using the -goterms, -iprlookup, and -pa options, respectively. Additionally, protein sequences were searched against the UniProt Swiss-Prot database (573661 sequences) using DIAMOND (v2.1.13.167) (Buchfink et al. 2015). GO terms were transferred only from high-confidence matches satisfying the thresholds of E-value ≤ 1 × 10⁻⁵, sequence identity ≥40%, and query coverage ≥50%. To minimize biologically misleading cross-kingdom annotations, taxonomy-based filtering was restricted to fungal-associated annotations (Boutet et al. 2007). To complement homology-based annotations, deep learning–based functional predictions were also generated using DeepFRI. Only GO annotations with confidence scores ≥0.5 were retained as high-confidence predictions for downstream analyses (Gligorijević et al. 2021).

### 2.7 GO enrichment

Gene Ontology (GO) terms obtained from eggNOG-mapper, InterProScan, Swiss-Prot fungal-filtered annotations, and high-confidence DeepFRI predictions were integrated to generate a comprehensive functional annotation dataset for all orthogroups. GO terms from multiple annotation sources were merged for each orthogroup, and duplicate GO identifiers were removed before downstream enrichment analysis. GO enrichment analysis was performed using the topGO (Alexa and Rahnenführer 2009) and GO.db packages (https://bioconductor.org/packages/GO.db). Fisher’s exact test was conducted using the “classic” algorithm, with all annotated orthogroups used as the background dataset (Upton 1992). Multiple testing correction was performed using the Benjamini–Hochberg false discovery rate (FDR) method, and GO terms with FDR ≤ 0.05 were considered significantly enriched. To reduce redundancy arising from GO term hierarchies, broad parent terms were removed when their child terms were significantly enriched.

### 2.8 Identification of virulence-associated gene categories

Carbohydrate-active enzymes (CAZymes) were identified using dbCAN3 (https://bcb.unl.edu/dbCAN2/index.php; accessed on November 4, 2025) (Zheng et al. 2023). Protein sequences were annotated using three complementary approaches with default parameters: HMMER against the dbCAN database (E-value < 1 × 10⁻¹⁵; coverage > 0.35), DIAMOND searches against the CAZy database (E-value < 1 × 10⁻¹⁰²), and HMMER searches against dbCAN-sub (E-value < 1 × 10⁻¹⁵; coverage > 0.35). Only CAZyme families supported by at least two of the three methods were retained as high-confidence predictions for downstream comparative analyses. Genes associated with pathogenicity and virulence were identified through sequence similarity searches against PHI-base (http://www.phi-base.org/, release 4.18) (Urban et al. 2020). Protein sequences from each strain were queried against the PHI-base protein dataset using DIAMOND in BLASTP mode with an E-value cutoff of 1 × 10⁻⁵ and a minimum sequence identity threshold of 40%. For each query, the top-scoring hit was extracted to generate final hits containing PHI-base annotations. Secreted proteins were predicted using a multi-step filtering strategy. Initially, all predicted proteins were screened for N-terminal signal peptides using SignalP (v6.0) in eukaryotic mode (Teufel et al. 2022). Proteins containing classical signal peptides were subsequently analyzed using TargetP (v2.0) in non-plant mode to distinguish secretory signal peptides from mitochondrial targeting peptides (Emanuelsson et al. 2007). Only proteins predicted as secreted were retained for downstream analyses. To further refine the secretome, subcellular localization was predicted using DeepLoc (v2.1) with the accurate prediction model (Armenteros et al., 2017). Proteins predicted as extracellular, containing a signal peptide, and classified as soluble non-membrane proteins were retained as high-confidence secreted proteins. These extracellular proteins were subsequently analyzed using EffectorP (v3.0) in fungal mode to identify candidate effector proteins using deep learning–based classification (Sperschneider and Dodds 2022).

### 2.9 BGC prediction and similarity network analysis

Biosynthetic gene clusters (BGCs) were predicted from the genome assemblies and corresponding gene annotation (GFF) files of 19 *B. sorokiniana* strains using the fungal version of antiSMASH v8.0.4 (Blin et al. 2025). Genomes were analyzed in fungal mode with the detection strictness set to “relaxed” to enable sensitive identification of both canonical and atypical secondary metabolite clusters. The resulting antiSMASH GenBank (.gbk) output files from all strains were subsequently analyzed using BiG-SCAPE (v2.0.0) to cluster related BGCs into Gene Cluster Families (GCFs) and construct BGC similarity networks (Zdouc et al. 2025; Draisma et al. 2026). Domain annotation during BiG-SCAPE analysis was performed using the Pfam-A HMM database (released May 28, 2025). Analyses were optimized for fungal BGCs using recursive input mode, global (“glocal”) alignment strategy, greedy extension, and inclusion of singleton clusters. Hybrid cluster duplication across multiple biosynthetic classes was prevented by using the -- hybrids-off option to avoid artificially inflating BGC counts. Clustering was performed using a GCF cutoff threshold of 0.4 with region-based records (--record-type region) (Kim and Dettman 2025). The resulting BGC similarity networks were visualized in Cytoscape (v3.10.4) (Shannon et al. 2003), with related BGCs grouped into GCFs based on sequence similarity and shared domain architecture. Representative clusters and syntenic relationships were further compared using clinker (v0.0.31) (Gilchrist and Chooi 2021), enabling visualization of gene organization conservation and pairwise cluster similarity across strains.

### 2.10 Gene family evolution analysis

Gene family evolution across 19 *B. sorokiniana* strains was performed using CAFE5 (Computational Analysis of gene Family Evolution; v5.0), which models gene family expansion and contraction under a stochastic birth–death evolutionary framework (Mendes et al. 2020). Orthogroup gene count matrices generated from the pangenome analysis were used as input for CAFE5. To improve statistical robustness, gene families absent at the inferred ancestral root node were excluded prior to analysis. A species phylogeny was obtained from the OrthoFinder species tree and subsequently midpoint-rooted. The tree was converted into an ultrametric form with the ape package v5.7-1 (Paradis et al. 2004), applying a penalized likelihood approach to ensure all terminal taxa were equidistant from the root. The resulting ultrametric phylogeny was used for downstream modeling of gene family evolution. CAFE5 analyses were performed using three gamma rate categories (-k 3) and a Poisson distribution for ancestral family size estimation (-p).

Lambda optimization was conducted with a maximum of 1000 iterations (-I 1000), and significantly expanded or contracted gene families were identified using a significance threshold of *P* ≤ 0.01. Gene families identified as significantly expanded or contracted were subsequently functionally annotated using eggNOG, CAZyme classifications, and PHI-base.

### 2.11 Selection pressure analysis of core genes

Orthology inference using OrthoFinder identified 8,396 single-copy orthogroups conserved across all *B. sorokiniana* strains. To increase the number of loci available for evolutionary analyses, the output was filtered to retain orthogroups present in all strains and occurring as single-copy in at least 70% of the 19 genomes (Santos and Kapheim 2024). For orthogroups containing duplicated genes in any strain, all paralogous copies from that strain were removed prior to downstream analyses to preserve orthology relationships. This additional filtering yielded 1,697 orthogroups, bringing the total to 10,093 orthogroups subjected to selection pressure analyses. Protein sequences for each orthogroup were aligned using MUSCLE (v3.8.1551) (Edgar 2004), and the corresponding coding sequences were converted into codon-aware nucleotide alignments using PAL2NAL (v14.1) to preserve reading frame integrity (Suyama et al. 2006). Codon alignments were generated with and without gap removal to evaluate codon retention. Only alignments that retained at least 50% of the original codon positions were used for downstream analyses. Orthogroups failing this criterion were subjected to iterative refinement, in which sequences contributing the highest proportion of gap-containing codons were sequentially removed until ≥50% codon retention was achieved, while maintaining a minimum of 13 sequences to preserve phylogenetic representation. All 9520 retained codon alignments were further screened for ambiguous nucleotides. If any codon position contained ambiguous bases (“N”) in any strain, the entire codon column was removed across all sequences to maintain codon frame consistency and alignment quality. The resulting high-confidence codon alignments were subsequently used for phylogenetic reconstruction with IQ-TREE (v3.0.1) under codon-substitution models selected with ModelFinder and AICc-based model selection (Kalyaanamoorthy et al. 2017; Wong et al. 2026).

Alignments exhibiting limited sequence variation were analyzed using conservative Goldman–Yang codon models with safe optimization parameters. Selection pressure analyses were conducted using HyPhy (v2.5.64) (Pond et al., 2020), which implements maximum-likelihood and Bayesian methods for estimating synonymous (dS) and nonsynonymous (dN) substitution rates. Multiple complementary methods were employed to detect pervasive, episodic, and lineage-specific adaptive evolution. Site-level pervasive selection was evaluated using FEL (Fixed Effects Likelihood) (v2.6) (Kosakovsky Pond and Frost 2005), which estimates codon-specific dN/dS ratios across phylogeny to identify sites under positive or purifying selection. Episodic diversifying selection affecting subsets of lineages was detected using MEME (Mixed Effects Model of Evolution) (v4.1) (Murrell et al. 2012), while gene-wide episodic selection was assessed using BUSTED (Branch-site Unrestricted Statistical Test for Episodic Diversification) (v4.6) (Murrell et al. 2015). Branch-specific adaptive evolution was further evaluated using aBSREL (adaptive Branch-Site Random Effects Likelihood) (v2.5) (Smith et al. 2015), which identifies individual phylogenetic branches experiencing episodic positive selection.

Statistical significance was determined using a threshold of *p* < 0.05. Evidence from FEL, MEME, aBSREL, and BUSTED was integrated to classify orthologous gene groups according to the strength of adaptive evolution signals. Genes were categorized as exhibiting Strong Positive Selection when all four methods supported adaptive evolution, including positively selected codon sites identified by FEL, episodically selected sites detected by MEME, positively selected branches identified by aBSREL, and significant gene-wide episodic selection detected by BUSTED (*p* < 0.05). Genes supported by at least two methods were classified as Positive Selection, whereas genes lacking sufficient evidence were categorized as Not Positive Selection. This integrative multi-model framework enabled robust identification of pervasive, episodic, and lineage-specific selection patterns across the *B. sorokiniana* core genome.

### 2.12 Genome Compartmentalization Analysis

Genome compartmentalization was assessed using the chromosome-level assembly of strain LK93. Genes were ordered by chromosomal position, and local gene density was quantified using 5′ and 3′ flanking intergenic region (FIR) lengths, calculated as the distances to the nearest upstream and downstream genes, respectively; Genes located at chromosome termini and loci lacking valid upstream or downstream neighbors were excluded from the analysis. The distributions of effectors, CAZymes, PHI genes, positively selected genes, and CAFE-expanded/contracted gene families were compared with all other genes using the Mann– Whitney U test with FDR correction (*p* < 0.05) and visualized as log₁₀-transformed hexbin density plots to identify enrichment in gene-dense or gene-sparse genomic compartments (Parker et al. 2025).

### 2.13 Protein–protein interaction (PPI) network analysis

Protein–protein interaction (PPI) network analysis was performed to investigate the interaction architecture, functional organization, and central regulatory proteins associated with core orthogroups in *B. sorokiniana*. Initially, the complete proteome of the D2 strain was uploaded to the STRING database v12.0 (https://string-db.org/) (organism identifier STRG0A63ZQP) (Mering et al. 2003). Predicted interaction data comprising 2,313,410 protein interaction pairs were retrieved from the STRING database (accessed April 30, 2026). To focus specifically on conserved components of the species pangenome, only interactions in which either the source or target protein belonged to a core orthogroup were retained, resulting in 1,113,261 unique interaction pairs for downstream analyses. To construct a robust, biologically meaningful interaction network, STRING interactions were filtered using a medium-confidence threshold (combined interaction score ≥ 400). Following filtering, 224,198 high-confidence interactions were retained, whereas 889,063 lower-confidence interaction pairs were excluded. The final network consisted of 6,539 protein nodes connected by 224,198 interaction edges.

Network decomposition identified a giant, connected component containing 6,450 nodes and 224,128 edges, while isolated proteins and minor disconnected components were excluded from subsequent topological analyses. Global network properties, including mean degree, network density, average path length, graph diameter, and global clustering coefficient, were calculated using the igraph package (v2.3.2) in R (Csardi and Nepusz 2006). The PPI network was analyzed as an undirected weighted graph using six centrality metrics (Degree centrality, Betweenness centrality, PageRank, Eigenvector centrality, Closeness centrality, and local Clustering coefficient) to identify hub proteins, which were integrated into a composite centrality index by averaging the rank-normalized values of all six metrics. The top 10% of proteins were used for Louvain community detection and modularity analysis, while the top 1% were designated as global hubs. Functional annotations (CAFE, CAZymes, effectors, PHI-base homologs, and positively selected orthogroups) were overlaid to examine their distribution within network modules.

## 3. Results

### 3.1 Assembly statistics and completeness of *B. sorokiniana* genomes

A total of 19 *B. sorokiniana* genomes representing diverse geographical and evolutionary origins were analyzed. Genome sizes ranged from 33.91–39.07 Mb with GC content of 49.24–51.44%. Assembly contiguity varied substantially, with most recently sequenced genomes assembled into 16–43 contigs and high N50 values (>2.2 Mb), whereas D2, SI, and BS52 were more fragmented. Despite these differences in contiguity, total assembly lengths remained broadly comparable across strains, indicating that fragmentation primarily affected scaffold continuity rather than overall genome completeness **(Table S2)**. BUSCO analysis indicated consistently high gene-space completeness across all 19 genomes, with 3,750–3,776 (99.0–99.7%) complete BUSCOs and very few fragmented (0–2) or missing (10–36) orthologs **(Figures 1A and 1B)**. Most genomes contained predominantly single-copy BUSCOs, although BS112 showed the highest duplication (25 BUSCOs). At the same time, BRIP27492a and LK93 exhibited entirely single-copy complete profiles, confirming that all genomes possessed highly complete gene repertoires and were therefore suitable for robust comparative pangenomic, evolutionary, and network-based analyses **(Table S3)**.

**Figure 1:**
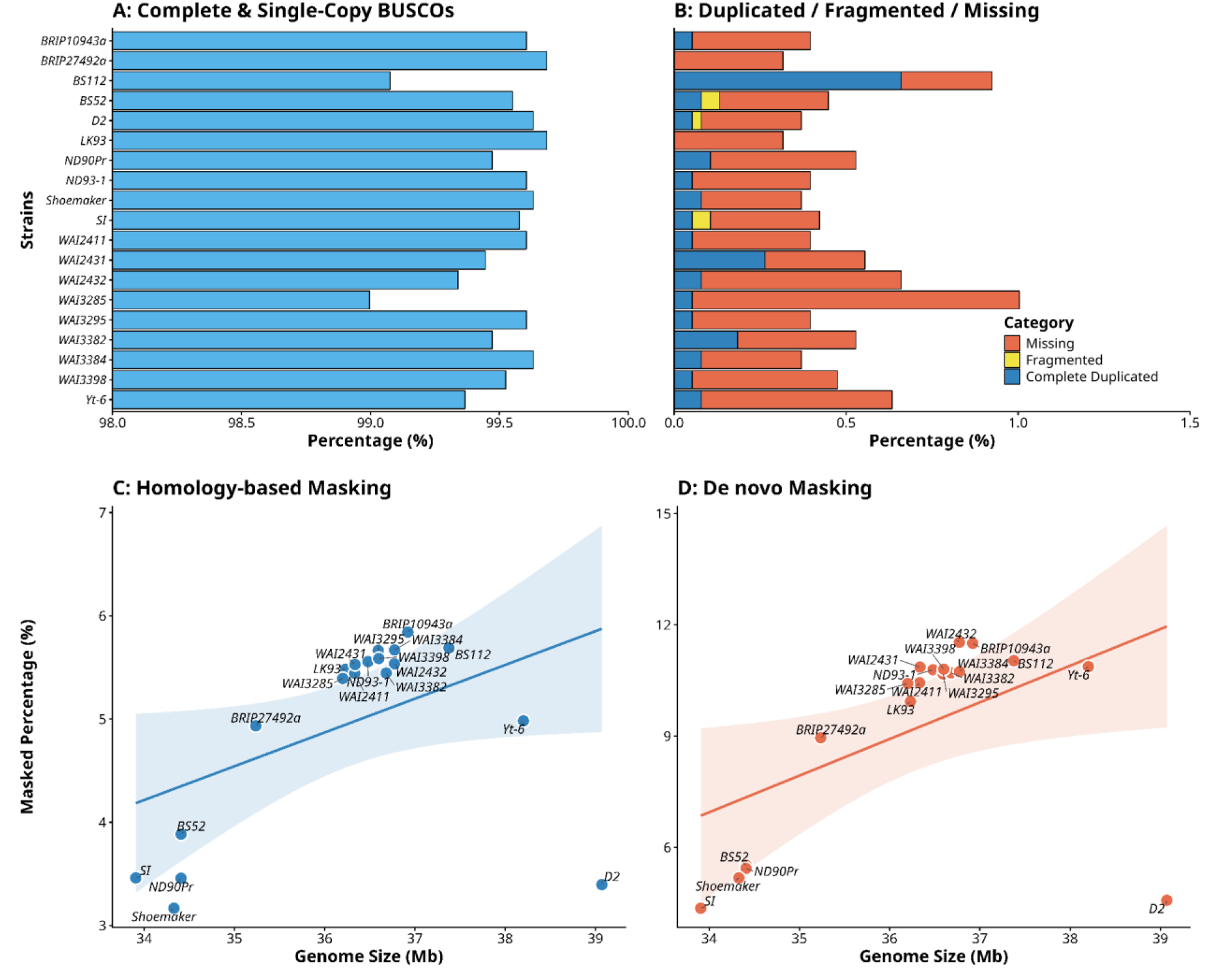
Genome completeness and repeat landscape across *Bipolaris sorokiniana* strains. (A) Percentage of complete and single-copy BUSCOs for each strain, showing consistently high genome completeness (∼99%), indicating high-quality assemblies across all isolates. (B) Distribution of the remaining BUSCO categories, including completely duplicated, fragmented, and missing genes, highlighting minor variation among strains with very low levels of duplication and missing content. (C) Relationship between genome size (Mb) and percentage of repeats identified using homology-based masking (HMM-DFam v3.9). A positive trend suggests that larger genomes tend to harbor a higher proportion of known repetitive elements. (D) Relationship between genome size (Mb) and percentage of repeats identified using de novo prediction (RepeatModeler v2.0.7), capturing novel and lineage-specific repeats. Similar to panel C, an increase in genome size is associated with higher repeat content.

### 3.2 Repeat landscape across the *B. sorokiniana* pangenome

Repeat masking using both *de novo* and homology-based approaches showed that *de novo* prediction consistently identified a larger repetitive fraction (4.36–11.53%) than homology-based masking (3.17–5.84%), indicating the presence of lineage-specific or previously uncharacterized repeats. WAI2432 (11.53%), BRIP10943a (11.50%), and BS112 (11.03%) contained the highest repeat contents, whereas D2, SI, Shoemaker, ND90Pr, and BS52 exhibited comparatively lower repeat proportions (4.36–5.52%). Retroelements, particularly LTR retrotransposons, were the predominant repeat class, with Gypsy/DIRS1 elements accounting for ∼4.95% of the genome, while Ty1/Copia elements occurred at lower abundance (<1%). DNA transposons represented the second largest repeat class (0.7–3.26%), predominantly Tc1/IS630/Pogo elements, and unclassified repeats accounted for 0.7–2.0% of the genomes, suggesting species-specific repeat families **(Tables S4A and S4B)**. *De novo* masking recovered nearly twice as much repeat content as homology-based methods, highlighting the importance of *de novo* repeat identification in *B. sorokiniana* **(Table S5)**. Genome size showed a positive correlation with repeat content identified by both homology-based and *de novo* approaches. Larger genomes consistently harbored higher proportions of repetitive DNA, including both known and lineage-specific repeat elements **(Figures 1C and 1D)**.

### 3.3 High-confidence gene prediction and annotation across 19 *B. sorokiniana* genomes

Gene prediction using protein homology, *ab initio* prediction, and transcript evidence produced 13,124–14,578 protein-coding genes per genome **(Table S6)**. Annotation quality was high, with most gene models exhibiting low Annotation Edit Distance (AED) scores **(Figure 2A)**, indicating strong agreement with biological evidence. RNA-seq/EST-supported annotations showed the lowest AED values, followed by protein homology. In contrast, SNAP and AUGUSTUS predictions displayed slightly broader AED distributions but remained largely below the filtering threshold. AED score distributions were consistent across genomes **(Figure 2B)**, with median values close to zero. Although the WAI strains exhibited slightly broader AED distributions, the vast majority of gene models across all genomes had AED ≤ 0.5, supporting their reliability. Filtering retained 12,867–14,241 high-confidence genes, representing >97% of predicted genes in every genome **(Figure 2C)**. Evidence composition varied among strains. Protein homology contributed 12.46– 64.67% of filtered annotations and predominated in ND90Pr, Shoemaker, BS112, BRIP10943a, and BRIP27492a. AUGUSTUS-supported predictions were more abundant in D2, SI, and WAI2431, whereas SNAP predictions were enriched in BS52, ND93-1, and Yt-6. RNA-seq/EST evidence supported 11.81–30.55% of annotations, with the highest proportions in LK93, BRIP10943a, BRIP27492a, and D2 **(Figure 2D and Table S7)**. The low AED scores, high retention of filtered genes, and complementary evidence support the robustness and suitability of the annotations for downstream comparative genomics, pangenome, and functional analyses.

**Figure 2:**
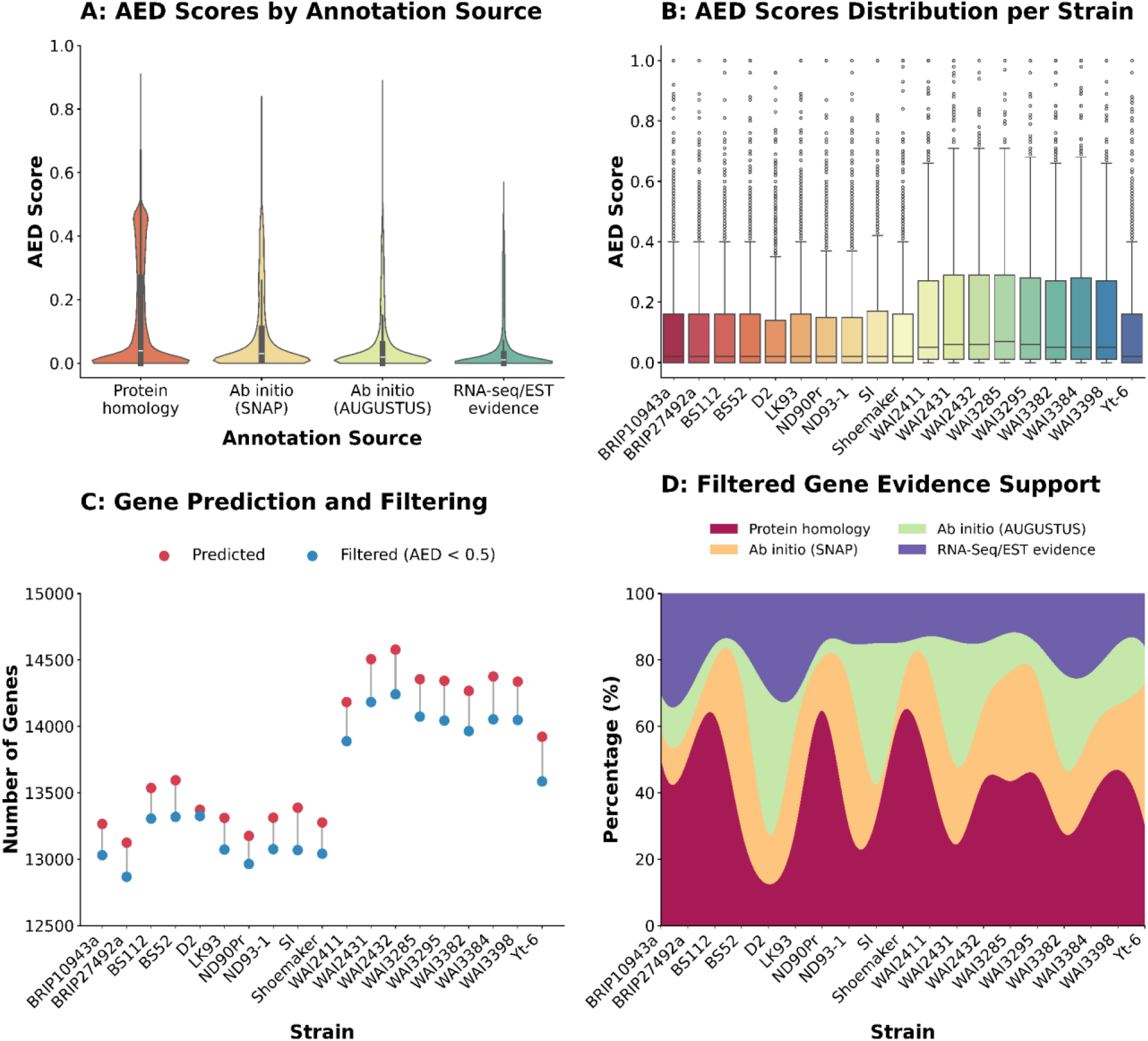
Evaluation of gene prediction quality and evidence support across *Bipolaris sorokiniana* strains. (A) Distribution of Annotation Edit Distance (AED) scores across four gene prediction evidence sources: protein homology, ab initio prediction (SNAP and AUGUSTUS), and RNA-seq/EST evidence. Lower AED values indicate stronger agreement between predicted gene models and supporting evidence. (B) AED score distributions across 19 strains, illustrating overall annotation quality and consistency. A threshold of AED ≤ 0.5 was applied to retain high-confidence gene models for downstream analyses. (C) Comparison of total predicted gene models and filtered high-confidence genes (AED < 0.5) for each strain. (D) Proportional contribution of different evidence types supporting the filtered gene models, shown as a stacked distribution across strains.

### 3.4 Construction and partitioning of the *B. sorokiniana* pangenome

Orthology analysis of 19 *B. sorokiniana* genomes identified 257,142 predicted genes, of which 255,325 (99.3%) were assigned to 15,164 orthogroups, while only 1,817 (0.7%) remained unassigned, indicating a highly conserved gene repertoire with minimal orphan genes. After incorporating unassigned genes and strain-specific orthogroups, the complete pangenome comprised 16,981 orthogroups with a mean of 16.8 genes per orthogroup (median = 19) **(Table S8)**. The pangenome consisted of 10,323 (60.8%) core, 1,444 (8.5%) softcore, 3,350 (19.7%) shell, and 1,864 (11.0%) cloud orthogroups **(Figure 3A)**. Among the core orthogroups, 8,396 were single-copy genes conserved across all genomes. In contrast, the accessory genome (softcore, shell, and cloud) accounted for 39.2% of the pangenome, indicating substantial intraspecific genomic diversity despite a large conserved core. Softcore orthogroups showed relatively limited variation (1,201–1,433 per genome), with ND93-1, BS52, BRIP10943a, BS112, and LK93 containing the highest numbers, while WAI2432 and WAI3285 contained the fewest. In contrast, shell orthogroups varied markedly, with WAI strains exhibiting consistently larger shell genomes (1,446–1,598 orthogroups) than non-WAI strains, including BRIP27492a (1,003), ND90Pr (1,013), and LK93 (1,056). A similar pattern was observed for cloud orthogroups, which ranged from 3 (BRIP10943a) to 294 (WAI2432), with most WAI strains possessing substantially higher numbers than non-WAI isolates, indicating greater lineage-specific accessory genome diversification **(Table S9)**. Pangenome growth analysis showed a continuous increase in orthogroup number from 13,884 (two genomes) to 16,981 (19 genomes). In contrast, the core genome declined from 12,222 to 10,323 orthogroups before stabilizing **(Figure 3B)**, consistent with an open pangenome. Hierarchical clustering of orthogroup presence–absence profiles further revealed extensive variation in accessory gene content among strains while confirming the conservation of the core genome **(Figure 3C)**.

**Figure 3:**
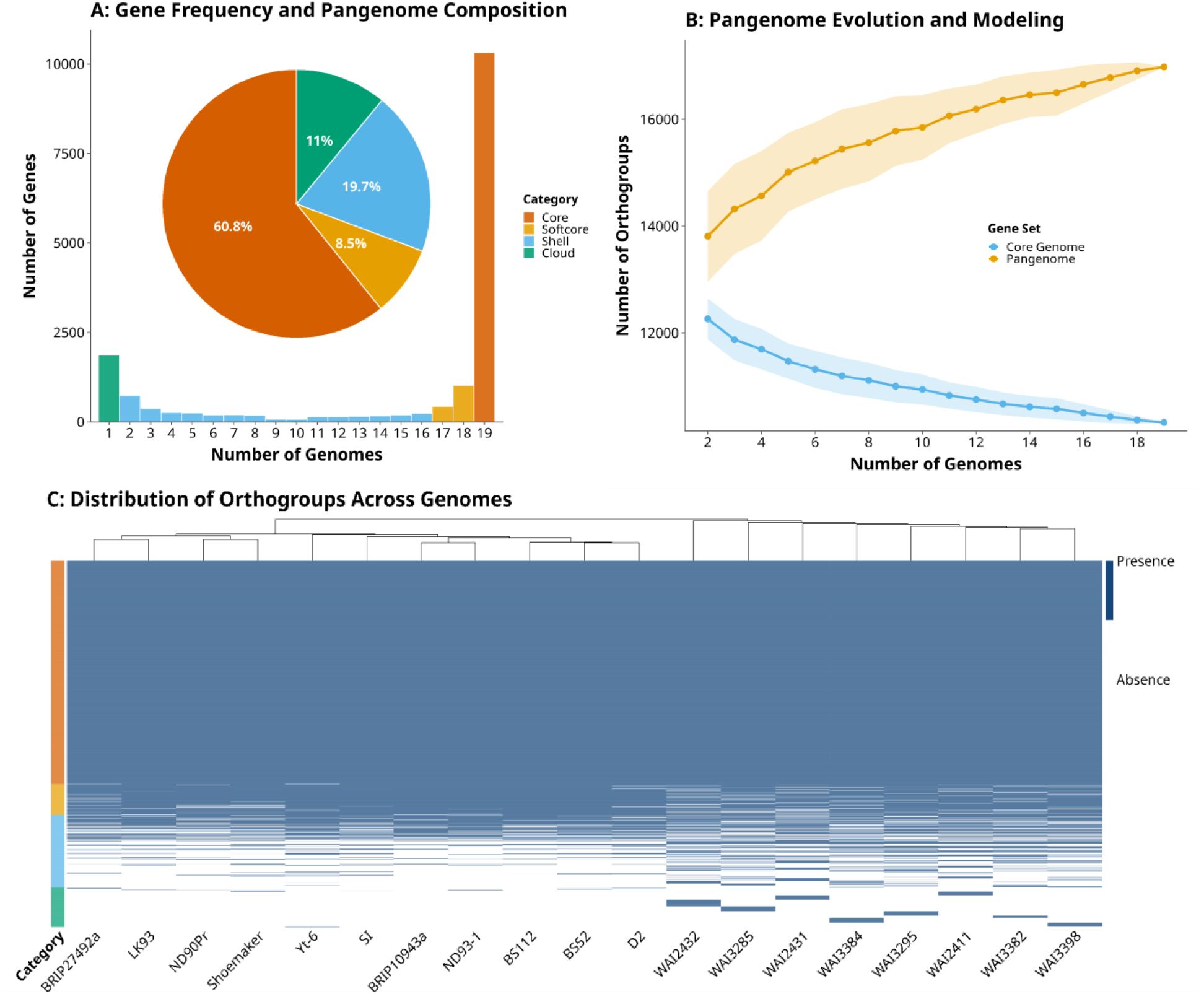
Pangenome structure, evolution, and orthogroup distribution across *Bipolaris sorokiniana* strains. (A) Distribution of orthogroup frequency across genomes, showing the number of genes shared among different numbers of strains. Genes are classified into four pangenome categories: core, softcore, shell, and cloud. The inset pie chart summarizes the proportional contribution of each category, highlighting the overall pangenome composition. (B) Pangenome and core genome dynamics estimated through iterative genome addition simulations. The pangenome curve shows a continuous increase in total orthogroups with the inclusion of additional genomes, indicating an open pangenome, while the core genome curve decreases and stabilizes, reflecting conserved gene content. Shaded regions represent variation across simulation replicates. (C) Presence–absence heatmap of orthogroups across all strains, with hierarchical clustering applied to genomes. Rows represent orthogroups and columns represent strains, with blue indicating presence and white indicating absence.

### 3.5 Functional landscape of pangenome

Functional annotation coverage declined progressively from the conserved to the accessory genome. Among the 10,323 core orthogroups, 99.0%, 86.7%, and 76.8% were annotated by NR, eggNOG, and InterProScan, respectively, whereas Swiss-Prot and DeepFRI assigned functions to 28.0% and 33.5% of orthogroups **(Tables S10, S11, S12, S13, and S14)**. Annotation coverage was lower in softcore orthogroups (NR: 94.7%; eggNOG: 59.1%; InterProScan: 48.0%; Swiss-Prot: 13.4%; DeepFRI: 29.7%) and shell orthogroups (NR: 91.2%; eggNOG: 63.7%; InterProScan: 44.6%; Swiss-Prot: 14.9%; DeepFRI: 29.1%). Cloud orthogroups showed the lowest overall coverage (NR: 84.6%; eggNOG: 70.2%; InterProScan: 42.9%; Swiss-Prot: 20.0%; DeepFRI: 29.7%), indicating that conserved orthogroups are enriched in well-characterized functions, whereas accessory and strain-specific orthogroups comprise a greater proportion of poorly characterized or rapidly evolving proteins (**Table S15)**.

### 3.6 Functional specialization among pangenome compartments

Integrated annotations from EggNOG, InterProScan, Swiss-Prot, and DeepFRI assigned GO terms to 11,431 (67.3%) of the 16,981 orthogroups. InterProScan provided the broadest coverage (8,267; 48.7%), followed by DeepFRI (5,413; 31.9%), EggNOG (4,905; 28.9%), and Swiss-Prot (3,565; 21.0%) **(Figure S1)**. Although EggNOG annotated fewer orthogroups, it assigned the highest average number of GO terms per orthogroup (88.5), whereas InterProScan produced more conservative annotations (3.9 GO terms per orthogroup). The integrated dataset averaged 47.9 GO terms per orthogroup (maximum 593), with 43.8% of orthogroups supported by a single annotation source and 56.2% supported by two or more sources, demonstrating the complementary value of combining annotation methods. GO annotation coverage was highest in the core genome (7,572/10,323; 73.4%), followed by cloud (61.5%), shell (57.2%), and softcore (55.1%) orthogroups. Core orthogroups showed extensive functional characterization, whereas cloud orthogroups had the highest average GO term density (63.4 GO terms per orthogroup), followed by shell (49.2), core (46.4), and softcore (37.7), suggesting that many strain-specific genes are associated with multifunctional regulatory and adaptive biological processes despite their limited conservation across genomes.

Gene Ontology enrichment revealed clear functional partitioning among pangenome compartments, whereas no significantly enriched biological process terms were detected in the softcore genome (FDR ≤ 0.05). Core orthogroups were enriched for conserved housekeeping and primary metabolic functions, including small-molecule, carbohydrate, carboxylic acid, organophosphate, nucleotide, sulfur, and glucan metabolism, as well as transmembrane transport and mycotoxin biosynthesis, reflecting conserved cellular maintenance, nutrient acquisition, and secondary metabolism. Shell orthogroups were enriched for regulatory and adaptive processes, including regulation of metabolism and gene expression, RNA metabolism, development, cell communication, aromatic and heterocyclic compound biosynthesis, amino acid activation for non-ribosomal peptide biosynthesis, and intron homing, indicating roles in phenotypic plasticity, specialized metabolism, and genome plasticity. Cloud orthogroups showed the strongest enrichment for strain-specific regulatory functions, including transcriptional regulation, chromatin and chromosome organization, DNA conformation and replication, developmental and stress responses, interspecies interactions, meiotic nuclear division, and mitochondria–nucleus signaling, highlighting their contribution to genome remodeling, host adaptation, and ecological specialization **(Figure S2)**.

### 3.7 Virulence-associated genes across the pangenome

Comprehensive CAZyme profiling identified 158 CAZyme subfamilies distributed among six classes: GH (68), GT (30), CBM (28), AA (17), CE (10), and PL (5) **(Figure 4A)**. The CAZyme repertoire was highly conserved, with 99 core, 35 softcore, 17 shell, and 7 cloud subfamilies. Similarly, 533 of 598 CAZyme-associated orthogroups belonged to the core genome. In contrast, only 34, 5, and 26 orthogroups were assigned to the softcore, shell, and cloud compartments, respectively, indicating that carbohydrate metabolism and plant cell wall degradation machinery are largely conserved across *B. sorokiniana*. Individual genomes contained 396–585 CAZyme genes, with Yt-6 (585) harboring the largest repertoire and WAI3384 (396) and WAI2432 (403) the smallest. Several conserved CAZyme families predominated across genomes for instance AA7 (12–36 copies), AA3 (21–28), and AA9 (14–24) were the major AA families; CBM91_GH43 (2–5) dominated the CBMs; CE5 (9–14) and CE4 (6–12) were the principal carbohydrate esterases; and GH18 (11–20), GH16 (12–16), GH5 (12–16), GH3 (11–15), and GH47 (7–10) were the most abundant glycoside hydrolases. Within glycosyltransferases, GT2 (11–19) was the predominant family, followed by GT1 (5–10) and GT34 (3–8), while PL1 (3–6) and PL3 (4–5) were conserved across most strains. Reduced copy numbers across several major CAZyme families were consistently observed in WAI2432 and WAI3384, whereas the remaining strains maintained highly conserved CAZyme repertoires (**Tables S16 and S17)**.

**Figure 4:**
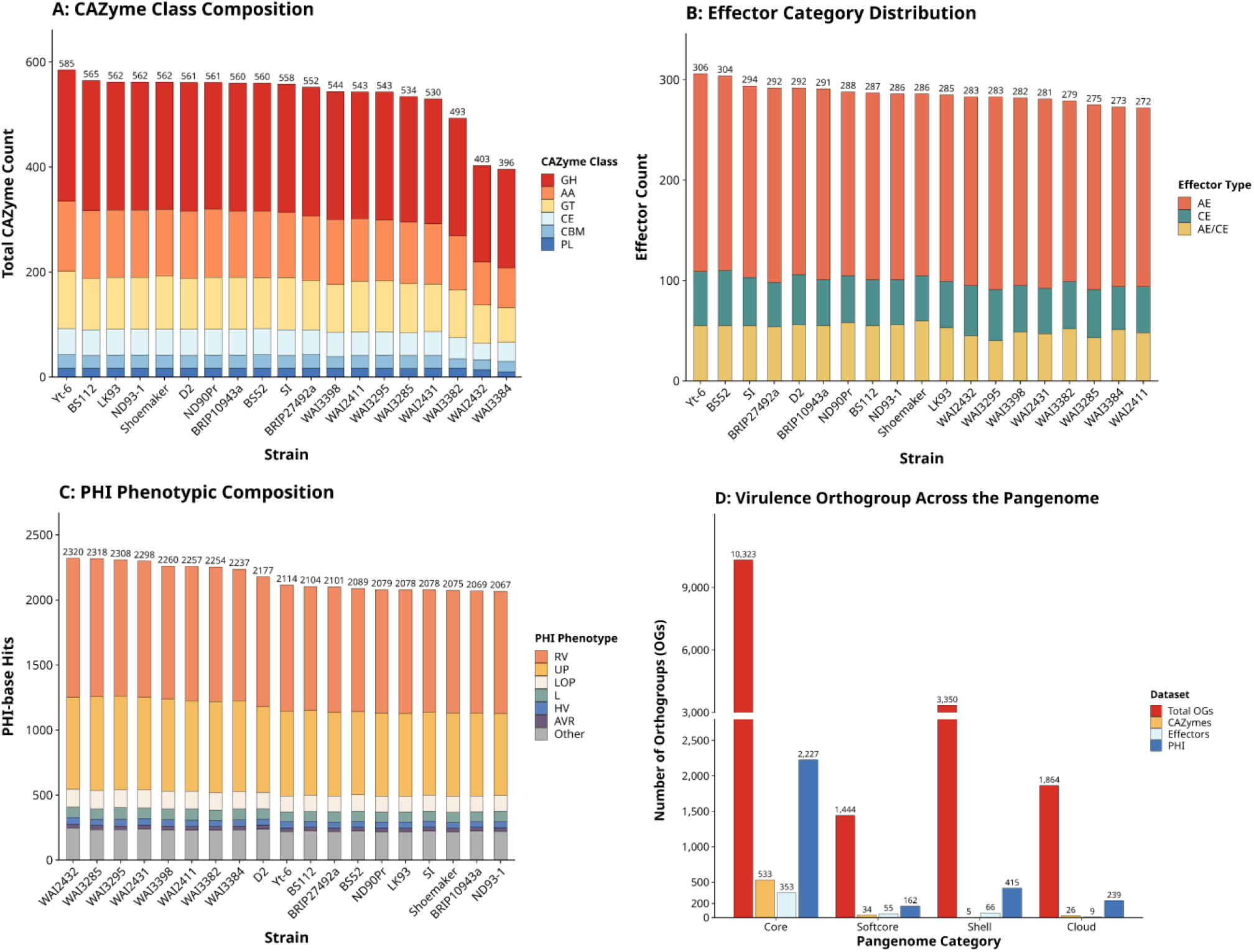
Comparative distribution of CAZymes, effectors, PHI-base genes, and virulence-associated orthogroups across the *Bipolaris sorokiniana* pangenome. (A) Stacked bar plot showing the distribution of carbohydrate-active enzyme (CAZyme) classes across all *Bipolaris sorokiniana* strains. CAZyme categories include glycoside hydrolases (GH), auxiliary activities (AA), glycosyl-transferases (GT), carbohydrate esterases (CE), carbohydrate-binding modules (CBM), and polysaccharide lyases (PL). Total CAZyme counts are shown above each bar, illustrating variation in carbohydrate metabolism and plant cell wall-degrading potential among strains. (B) Distribution of predicted effector proteins across strains, categorized as apoplastic effectors (AE), cytoplasmic effectors (CE), and dual-localized apoplastic/cytoplasmic effectors (AE/CE). (C) Composition of pathogen–host interaction (PHI-base) phenotypic categories identified across strains. PHI annotations were grouped into reduced virulence (RV), unaffected pathogenicity (UP), loss of pathogenicity (LOP), lethal (L), hypervirulence (HV), avirulence effectors (AVR), and other categories. (D) Distribution of total orthogroups (OGs), CAZyme-associated orthogroups, effector orthogroups, and PHI-associated orthogroups across pangenome compartments (core, softcore, shell, and cloud).

Comprehensive secretome prediction using SignalP, TargetP, and DeepLoc identified a high-confidence secretome of 14,865 extracellular proteins across the 19 genomes. SignalP initially detected 20,172 proteins with N-terminal secretion signals (973–1,151 per strain), which were reduced to 19,556 after TargetP filtering and, finally, to 727–850 extracellular proteins per genome following DeepLoc prediction. Yt-6 consistently possessed the largest secretome, whereas WAI2432 contained the fewest predicted secreted proteins, indicating an overall conserved secretory machinery across the *B. sorokiniana* pangenome. EffectorP identified 5,439 candidate effectors, comprising 3,213 apoplastic, 902 cytoplasmic, and 1,324 dual-localized effectors. Individual genomes encoded 272–306 effectors, with Yt-6 (306), BS52 (304), and SI (294) containing the largest repertoires, whereas WAI2411 (272) contained the fewest. Apoplastic effectors predominated in all strains (160–179 proteins), followed by dual-localized (59–76) and cytoplasmic (43–55) effectors **(Figure 4B)**. At the pangenome level, effector-associated orthogroups were predominantly conserved, with 353 of 483 orthogroups assigned to the core genome, compared with 55 softcore, 66 shell, and 9 cloud orthogroups, indicating that most host-interaction and virulence mechanisms are evolutionarily conserved across *B. sorokiniana* **(Tables S18 and S19)**.

To investigate the virulence potential of the *Bipolaris* pangenome, predicted proteins were also annotated against the PHI-base database, enabling the identification of proteins associated with experimentally validated pathogenicity and host–pathogen interaction phenotypes. PHI-base annotation identified 41,283 proteins associated with 44 experimentally validated pathogenicity phenotypes, revealing that virulence-associated functions are predominantly conserved across the *B. sorokiniana* pangenome. The core genome contained the vast majority of PHI-base hits (36,419 hits distributed across 2,227 orthogroups representing 42 phenotypes), whereas the softcore (2,266 hits; 162 orthogroups; 13 phenotypes), shell (2,352 hits; 415 orthogroups; 20 phenotypes), and cloud (246 hits; 239 orthogroups; 18 phenotypes) compartments contributed comparatively fewer virulence-associated genes. Notably, cloud-associated PHI genes were detected in only nine strains, indicating lineage-specific acquisition of virulence functions. Among all phenotype categories, reduced virulence was by far the most abundant (16,665 hits), followed by unaffected pathogenicity (11,119), loss of pathogenicity (2,167), and lethal (1,357) phenotypes, all of which were largely confined to the core genome. Conserved hypervirulence-associated genes (767 hits) and plant avirulence effector (452 hits) further demonstrate that the principal determinants of host colonization, pathogenic fitness, and host manipulation are evolutionarily stable within the species. In contrast, the accessory genome primarily contributed additional reduced-virulence and unaffected-pathogenicity genes. At the same time, the cloud compartment was enriched for strain-specific genes associated with reduced virulence (109 hits), unaffected pathogenicity (76), loss of pathogenicity (18), and lethal phenotypes (11), suggesting roles in niche adaptation and pathogenic specialization. Despite modest quantitative variation among genomes (2,067–2,320 PHI-associated genes per strain), all isolates maintained highly similar pathogenicity profiles. WAI2432 possessed the largest PHI-base repertoire (2,320 hits), whereas ND93-1 contained the fewest (2,067 hits), reinforcing the presence of a highly conserved virulence backbone across the *B. sorokiniana* population **(Figures 4C, 4D, and Table S20)**.

### 3.8 BGC diversity and similarity network architecture

Comprehensive BGC mining across the 19 *B. sorokiniana* genomes identified diverse secondary metabolite biosynthetic loci, including NRPS, NRPS-like, T1PKS, terpene, indole, hybrid NRPS– PKS, and metallophore-associated clusters. Individual genomes encoded 39–54 BGCs, with BS52 (54), SI (53), ND90Pr, and Yt-6 (51) harboring the largest repertoires, whereas WAI3398 (39) contained the fewest. Most strains possessed 42–50 BGCs, indicating a largely conserved but moderately dynamic secondary metabolite repertoire. Terpene clusters were the most abundant (15–18 per genome), followed by T1PKS (7–12) and NRPS (3–11) clusters. Hybrid classes, including NRPS-like, NRPS–indole, NRPS–terpene, and NRP–metallophore clusters, were broadly distributed at lower frequencies, whereas rare cluster types such as NRPS-like–terpene, NRPS– NRPS-like, and terpene–terpene precursor were restricted to individual strains, suggesting recent lineage-specific diversification **(Figure 5A and Table S21)**.

**Figure 5:**
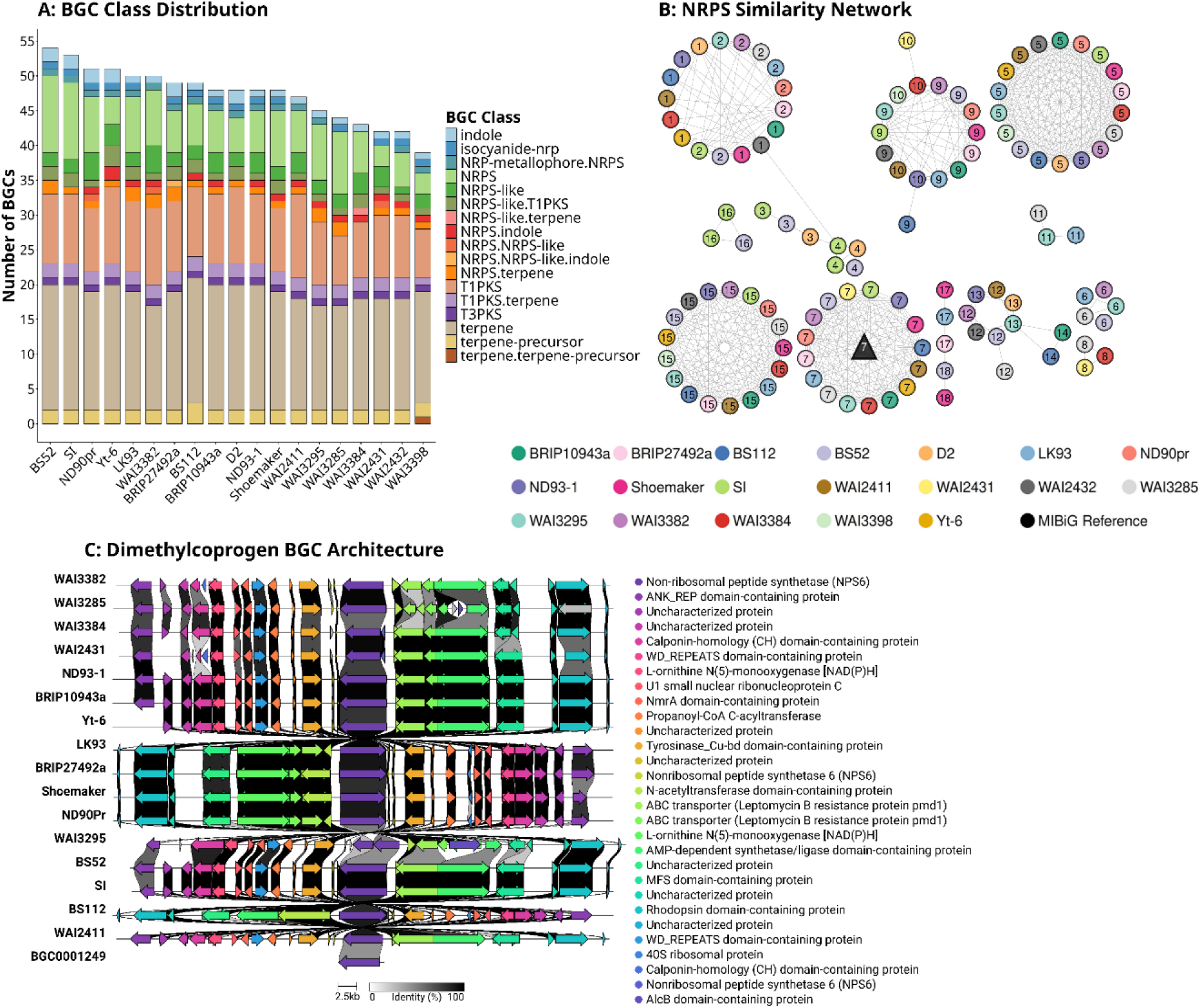
Biosynthetic gene cluster (BGC) diversity and NRPS similarity network analysis across the pangenome strains. (A) Stacked bar plot showing the distribution of biosynthetic gene cluster (BGC) classes identified across all analyzed strains using antiSMASH. Each bar represents an individual strain, while colored segments indicate different BGC classes. The y-axis represents the total number of predicted BGCs per strain. (B) Sequence similarity network of non-ribosomal peptide synthetase (NRPS) biosynthetic gene clusters generated from BiG-SCAPE clustering analysis. Circular nodes represent strain-derived NRPS clusters and are colored according to strain identity, whereas triangular nodes represent reference clusters from the MIBiG database. Edges indicate sequence similarity relationships among BGCs. Node labels within circles correspond to BGC family identifiers. (C) Genome architecture and genomic organization of the Dimethylcoprogen-associated biosynthetic gene cluster identified in the analyzed strains.

Sequence similarity network analysis generated using BiG-SCAPE revealed extensive connectivity among NRPS-associated BGCs and demonstrated strong relationships with previously characterized MIBiG reference clusters **(Figure 5B)**. Multiple NRPS clusters showed high similarity to the dimethylcoprogen BGC (MIBiG accession: BGC0001249) originally characterized in *Alternaria alternata*. Homologous dimethylcoprogen-associated clusters were identified in 16 *B. sorokiniana* genomes, suggesting broad conservation of siderophore-mediated iron acquisition pathways and metallophore-associated virulence mechanisms across the species **(Figure 5C)**. Similarly, NRPS-like similarity networks identified clusters closely related to the choline BGC (BGC0002276) from *Aspergillus nidulans* in 18 strains, indicating strong conservation of metabolite biosynthetic pathways potentially associated with membrane metabolism, stress adaptation, and fungal fitness (**Figure S3)**. T1PKS-associated similarity analysis further revealed widespread conservation of the 4-chloropinselin biosynthetic cluster (BGC0002726), previously described in *B. sorokiniana*, in 18 strains (**Figure S4)**. This cluster is linked to the biosynthesis of several polyketide-derived metabolites, including 4-chloropinselin, pinselin, chloromonilinic acid B, chloromonilinic acid D, and 4-hydroxyvertixanthone, highlighting conservation of phytotoxic and pigment-associated secondary metabolism within the species. Terpene-associated similarity networks identified homologous clusters related to the terpestacin biosynthetic pathway (BGC0002245) from *Bipolaris maydis* in all 19 *B. sorokiniana* strains, indicating complete conservation of this terpene-associated biosynthetic capability across the pangenome (**Figure S5)**. In addition, homologs of the prehelminthosporol/sativene biosynthetic cluster (BGC0003041), previously characterized in *B. sorokiniana* ND90Pr, were identified in 11 strains **(Figure S6)**. These terpene-associated clusters encode pathways involved in sesquiterpene biosynthesis and are probably associated with pathogenicity, host colonization, and ecological adaptation (**Tables S22 and S23)**.

### 3.9 Gene family expansion and contraction dynamics

CAFE analysis identified 434 orthogroups exhibiting significant gene family evolution (*P* < 0.01) across the 19 *B. sorokiniana* genomes (**Table S24)**. Of these, 192 orthogroups exhibited both expansion and contraction events across different lineages, 241 showed contraction only, and only a single orthogroup exhibited expansion only, indicating a strong overall bias toward gene loss during species diversification. Dynamic orthogroups were overwhelmingly associated with the accessory genome, with 389 shell, 34 softcore, and only 11 core orthogroups, demonstrating that gene family evolution primarily affects accessory genomic regions. Gene family dynamics varied among strains. Yt-6 exhibited the greatest turnover (84 expansions; 83 contractions), whereas ND93-1 (259 contractions), BRIP10943a (254), WAI2432 (241), WAI2411 (233), and WAI3384 (229) showed pronounced contraction, consistent with lineage-specific gene loss. Moderate expansion was observed in WAI3384 (46), WAI2432 (42), LK93 and Shoemaker (39 each), while BRIP27492a showed the fewest expansions (12). Overall, contraction events exceeded expansions in nearly all genomes, indicating progressive reductive evolution of accessory gene families **(Figures 6A, 6B, and Table S25)**. Functional annotation identified 84 rapidly evolving orthogroups associated with virulence and metabolism, including 65 PHI-base, 5 CAZyme, and 10 effector-associated orthogroups. Two orthogroups overlapped between the PHI-base and effector, and two others were shared between the PHI-base and CAZyme, highlighting functional overlap among pathogenicity- and metabolism-related gene families (**Table S26)**.

**Figure 6:**
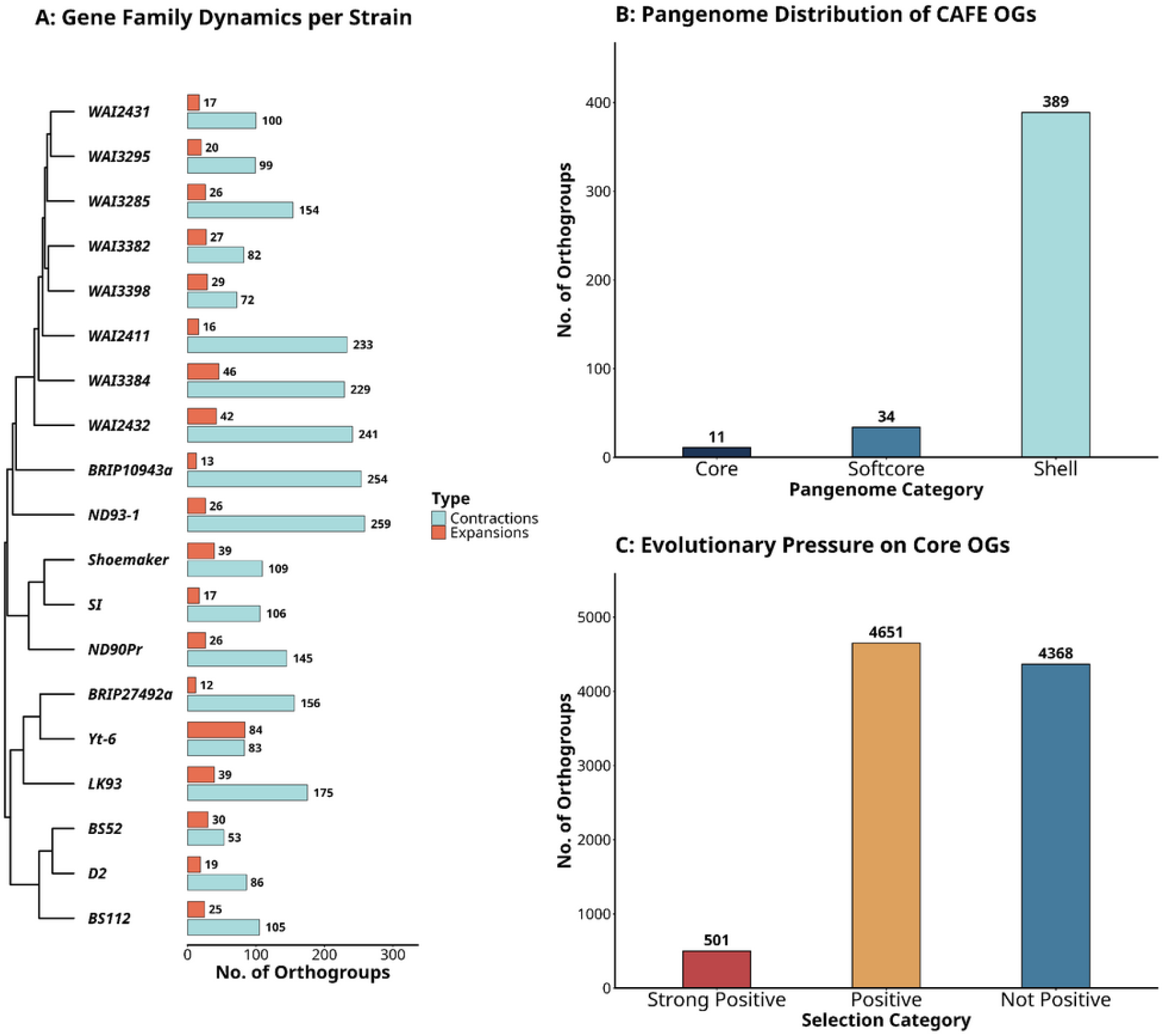
Gene family evolution and selection pressure across the *Bipolaris sorokiniana* pangenome. (A) Phylogenetic distribution of gene family expansions and contractions inferred using CAFE analysis across *Bipolaris sorokiniana* strains. The left panel shows the ultrametric species phylogeny, while the right panel displays the number of expanded and contracted orthogroups for each strain. (B) Distribution of CAFE-associated orthogroups across pangenome compartments. (C) Classification of core orthogroups based on selection pressure analysis. Orthogroups were categorized as strong positive selection, positive selection, or not under positive selection.

GO enrichment analysis revealed that contraction-only orthogroups were significantly associated with folate metabolism, RNA-directed DNA polymerase/transposable element-related functions, and host cell membrane, cytoplasm, and nucleus categories, suggesting lineage-specific loss of genes involved in host interaction and genome dynamics. In contrast, orthogroups exhibiting both expansion and contraction were enriched for secondary metabolite and nonribosomal peptide biosynthesis, toxin production, ferric-chelate reductase activity, copper ion import, glycerol, polyol, and amino acid transport, and histone H3-K9 deacetylation, indicating that rapidly evolving gene families contribute to metabolic adaptation, nutrient acquisition, epigenetic regulation, and virulence diversification within the *B. sorokiniana* pangenome (**Figure S7)**.

### 3.10 Evolutionary selection signatures across conserved core orthogroups

Selection analyses integrating FEL, MEME, BUSTED, and aBSREL identified widespread adaptive evolution across 9,520 conserved core orthogroups. Of these, 501 (5.3%) were classified as Strong Positive Selection orthogroups, supported by all four methods, 4,651 (48.9%) showed evidence of positive selection supported by at least two methods, and 4,368 (45.9%) were classified as Not Under Positive Selection. Method-specific analyses further identified 619 orthogroups with pervasive positive selection and 6,554 under purifying selection (FEL), 4,542 with episodic diversifying selection (MEME), 4,408 with gene-wide episodic positive selection (BUSTED), and 6,927 with branch-specific adaptive evolution (aBSREL). Collectively, these results demonstrate pervasive, episodic, and lineage-specific adaptive evolution, together with strong purifying selection, across the conserved core genome of *B. sorokiniana* **(Figure 6C and Table S27)**.

Functional annotation of the 501 orthogroups under strong positive selection revealed extensive enrichment of virulence- and adaptation-associated genes. These included 101 PHI-base, 19 CAZyme, and 4 effector-associated orthogroups, with 13 overlapping between the PHI-base and CAZyme datasets. The predicted effectors comprised UPF0016/GDT1-like, α/β-hydrolase, and two hypothetical proteins (COCSADRAFT_200222 and GGP41_008921). OG0004947, encoding a Class II fungal hydrophobin, was jointly annotated as a PHI-base-associated dual apoplastic/cytoplasmic effector. Two orthogroups (OG0000938 and OG0004798) overlapped among the CAZyme, effector, and PHI-base datasets, encoding an AA5 copper radical oxidase/glyoxal oxidase associated with unaffected pathogenicity in *Magnaporthe oryzae* and a GH132 SUN-domain β-glucosidase associated with reduced virulence in *Ustilaginoidea virens*, respectively. Additionally, OG0000151, encoding an aldehyde/histidinol dehydrogenase, overlapped between the PHI-base and CAFE expansion datasets and was associated with reduced virulence in *Parastagonospora nodorum*. COG classification showed that ‘function unknown’ (S) was the largest category (129 orthogroups). Among annotated functions, the most abundant categories were carbohydrate transport and metabolism (G; 32), post-translational modification/chaperones (O; 31), amino acid transport and metabolism (E; 28), lipid metabolism (I; 23), and intracellular trafficking and secretion (U; 23), followed by signal transduction (T; 18), secondary metabolite biosynthesis (Q; 15), RNA processing and transcription (A, J, and K; 13–14 each), inorganic ion transport (P; 13), energy production (C; 10), cytoskeleton organization (Z; 10), and cell wall/membrane biogenesis (M; 8), whereas defense, DNA repair, motility, extracellular structures, and nuclear structure were comparatively underrepresented (**Table S28)**.

### 3.11 *S*tructural compartmentalization of the *B. sorokiniana* genome

Genome compartmentalization analysis of the chromosome-level reference genome LK93 retained 10,662 of 13,311 protein-coding genes after excluding boundary-edge and overlapping loci. The analysis included 242 effectors, 459 CAZymes, 1,660 PHI-associated genes, 388 positively selected genes, and 296 CAFE-associated genes. Genome-wide distributions of flanking intergenic regions (FIRs) indicated a continuous genomic landscape rather than distinct gene-dense and gene-sparse compartments, suggesting that *B. sorokiniana* lacks a classical two-speed genome architecture. Nevertheless, candidate effectors were significantly enriched in relatively gene-sparse regions, exhibiting 1.50-fold and 1.60-fold larger median 5′ and 3′ FIRs than background genes (BH-adjusted *P* = 5.53 × 10⁻¹⁶). CAFE-associated genes (1.44- and 1.29-fold increases; BH-adjusted *P* = 5.39 × 10⁻¹¹) and CAZymes (1.24- and 1.19-fold increases; BH-adjusted *P* = 1.09 × 10⁻⁵) also showed significant enrichment in gene-sparse regions, whereas PHI-associated and positively selected genes did not differ significantly from the genomic background **(Figure S8 and Table S29)**.

### 3.12 Identification of hub proteins and functional communities within the core interactome

The protein–protein interaction (PPI) network of conserved core orthogroups comprised 6,539 nodes and 224,188 edges, with 6,450 nodes forming a giant connected component containing 224,128 interactions **(Figure S9 and Table S30)**. The network exhibited a mean degree of 69.5, an average path length of 2.94, a density of 0.0108, and a global clustering coefficient of 0.2612, indicating extensive functional connectivity. Annotation mapping revealed that the majority of nodes corresponded to uncharacterized or general proteins (62.1%), followed by PHI-associated (25.8%), positively selected (3.1%), CAZyme (2.8%), effector (0.8%), and CAFE-associated (0.4%) proteins, with several proteins carrying multiple functional annotations. Integration of six centrality metrics identified a densely connected top 10% subnetwork comprising 645 nodes and 50,458 edges, representing the core interactome **(Figure 7)**. Louvain community detection identified four major functional modules (modularity Q = 0.3170) with distinct biological characteristics. Community 1 (235 nodes; 8,687 edges; density = 0.3159) and Community 3 (87 nodes; 1,693 edges; density = 0.4526) were enriched in PHI-associated proteins, suggesting roles in pathogenicity, cellular metabolism, and survival. Community 2 (181 nodes; 12,179 edges; density = 0.7476) showed a high proportion of PHI-associated and positively selected proteins, indicating strong integration of adaptive and virulence-related pathways, whereas Community 4 (142 nodes; 8,052 edges; density = 0.8043) formed the most densely connected module and was enriched for conserved housekeeping and translation-associated functions (**Table S31)**.

**Figure 7:**
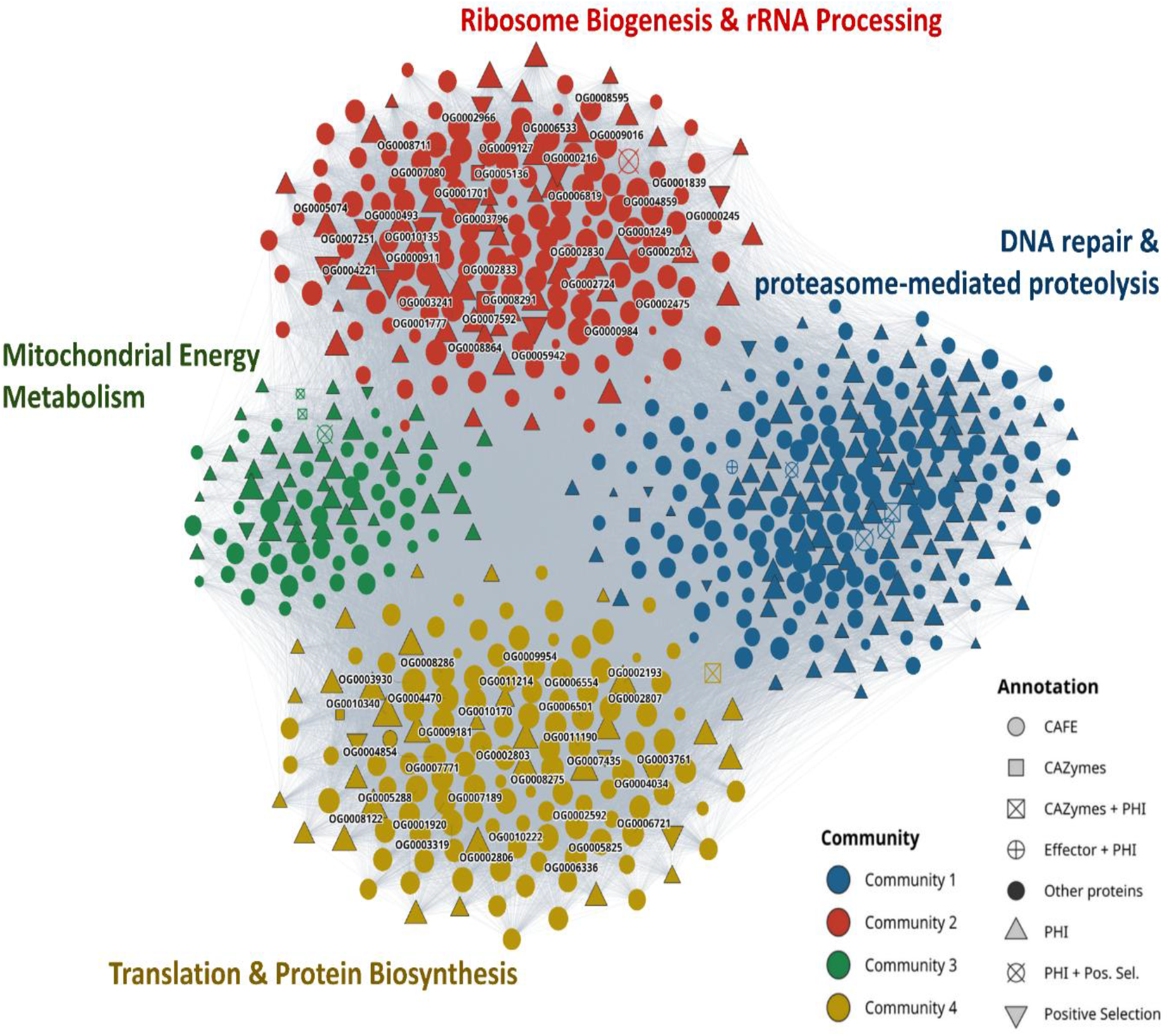
Protein–protein interaction (PPI) network architecture and functional community analysis of core orthogroups in *Bipolaris sorokiniana*. STRING-derived protein–protein interaction (PPI) network constructed from the top 10% highly connected core orthogroups, comprising 645 nodes and 50,458 edges. Nodes represent core orthogroups, while edges indicate predicted or known functional protein interactions retrieved from the STRING database. The top 1% hub orthogroups within the global network were highlighted to associate with modules. Community detection analysis identified four major functional communities (C1–C4). Community C1 was enriched in genome maintenance, proteostasis, DNA repair, ubiquitin-mediated proteolysis, and stress adaptation pathways, suggesting an important role in pathogenic fitness and host colonization. Community C3 was associated with mitochondrial energy metabolism and respiration-related functions linked to invasive growth, oxidative stress tolerance, and infection-associated energy demands. Communities C2 and C4 were primarily enriched in ribosome biogenesis, rRNA processing, translation, and protein biosynthesis pathways.

Functional enrichment analysis demonstrated that Communities 2 and 4 were strongly associated with ribosome biogenesis, translation, and ribonucleoprotein assembly pathways. Community 2 showed highly significant enrichment for rRNA processing, ribosome biogenesis, and ribonucleoprotein complex assembly functions, including GO terms associated with rRNA metabolic processes and preribosomal complex formation, indicating that this module represents a major ribosome biogenesis hub within the fungal interactome **(Figure S10)**. Similarly, Community 4 was predominantly enriched for translation-associated pathways, including cytosolic ribosome organization, translational initiation, and ribosomal structural constituents, consistent with its extremely high interaction density and connectivity **(Figure S11)**. In contrast, Community 1 displayed enrichment for DNA repair, DNA metabolic processes, and proteasome-associated functions, suggesting roles in genome maintenance and stress-response adaptation **(Figure S12)**, whereas Community 3 was associated with mitochondrial protein complexes, ATP synthesis, oxidative phosphorylation, and tricarboxylic acid cycle pathways, reflecting conserved energy metabolism and respiratory functions **(Figure S13)**. These results indicate that the *B. sorokiniana* core interactome is organized into highly structured functional modules integrating translation, ribosome biogenesis, metabolism, stress adaptation, and pathogenicity-associated processes.

## 4. Discussion

Comparative pangenome analysis has emerged as a powerful approach for understanding genome evolution, adaptation, and pathogenicity in fungal plant pathogens. Through the present integration of comparative genomics, evolutionary selection analyses, virulence profiling, and protein interaction networks, the present study provides a comprehensive view of genome plasticity and adaptive diversification in *B. sorokiniana*. The analysis of 19 strains revealed an open, highly dynamic genome architecture in *B. sorokiniana*, shaped by extensive diversification of the accessory genome, adaptive evolution, and virulence-associated functional innovation. While the conserved core genome retained essential metabolic and pathogenicity-related functions, the accessory genome comprised rapidly evolving genes associated with host interaction, secondary metabolism, and environmental adaptation.

### 4.1 Genome contiguity, masking strategies, and transposable element dynamics

Despite substantial variation in assembly contiguity among the 19 *B. sorokiniana* genomes, BUSCO analysis demonstrated consistently high gene-space completeness (∼99%), indicating that even fragmented draft assemblies can reliably support gene-based pangenome analyses when combined with rigorous gene prediction and filtering. This agrees with recent chromosome-scale assemblies generated using long-read sequencing, which have substantially improved genome continuity compared with earlier short-read assemblies (Bucknell et al. 2025). The combined *de novo* and homology-based repeat annotation strategy revealed that *de novo* prediction recovered nearly twice as many repeats as identified by RepeatMasker/DFam, indicating that a substantial proportion of the *B. sorokiniana* repeatome consists of lineage-specific or highly divergent transposable elements absent from current reference repeat libraries. Improved repeat recovery in chromosome-scale assemblies further reflects the ability of long-read sequencing to resolve repetitive regions that are often collapsed in short-read assemblies (Du and Liang 2019). The total repeat content observed in our genomes (13.89–17.34%) is comparable to that reported for other phytopathogenic Dothideomycetes, including *Setosphaeria turcica* (12.96%) and *Pyrenophora tritici-repentis* (∼16%) (Manning et al. 2013; Condon et al. 2013). The predominance of Gypsy/DIRS1 LTR retrotransposons and Tc1/IS630/Pogo DNA transposons indicates that these elements are the principal drivers of genome structural variation in *B. sorokiniana*. Similar repeat landscapes have been reported in its close relative *Cochliobolus heterostrophus* (*B. maydis*) and other Dothideomycetes, where Gypsy retrotransposons and Pogo-family DNA transposons also dominate the transposable element repertoire (Santana et al. 2014; Wang et al. 2020). These conserved patterns suggest that although repeat abundance varies among species and assemblies, the overall composition of mobile genetic elements has remained evolutionarily conserved across host-adapted filamentous fungi.

### 4.2 Functional stratification and compartmentalization of the pangenome

The analyses of the 19 *B. sorokiniana* genomes revealed a stratified pangenome. The identification of a large core genome (60.8%) and a substantial accessory fraction (39.2%) distributed across the softcore, shell, and cloud compartments aligns with the typical genome organization observed in filamentous ascomycetes. This structural distribution closely mirrors the 19-isolate reference-quality global pangenome described for the fungal wheat pathogen *Zymoseptoria tritici* (Badet et al. 2020), which similarly comprises a conserved core genome spanning only 60% of all genes, with the remaining 40% forming a highly dynamic accessory fraction that closely resembles the genomic partitioning observed in *B. sorokiniana*. A similar core to accessory genome ratio has also been reported in another major wheat pathogen, *P. tritici-repentis* (the causal agent of tan spot disease), where a 26-strain pangenome analysis identified a conserved core-gene content of 57%, supporting extensive accessory genome diversification among necrotrophic wheat pathogens (Moolhuijzen et al. 2022). Conversely, in the barley-specific net blotch pathogen *Pyrenophora teres* f. *teres*, a pangenome analysis of five strains revealed a significantly higher core genome proportion of 86.5% (Wyatt et al. 2020). Comparative genomics of 20 *Fusarium graminearum* strains, the causal agent of Fusarium Head Blight (FHB) with contrasting aggressiveness, the core fraction dropped to 44%, revealing an expanded, wide-open pangenome heavily governed by its accessory components (Alouane et al. 2021).

The progressive expansion of the *B. sorokiniana* pangenome without evidence of saturation further demonstrated that the species possesses an open pangenome architecture, consistent with Heaps’ law models reported for multiple Dothideomycete plant pathogens. Open pangenomes are considered a hallmark of highly adaptable cereal pathogens such as *Zymoseptoria tritici, P. tritici-repentis,* and *F. graminearum*, where continual acquisition and diversification of accessory genes facilitate adaptation to changing host environments, ecological pressures, and fungicide exposure (Guimarães et al. 2015). A notable feature of our dataset is the marked expansion of the accessory genome in the recently sequenced WAI lineage, which contains substantially larger shell and cloud compartments than previously sequenced reference strains. This expansion, accompanied by a reduction in the softcore fraction, suggests accelerated lineage-specific gene gain, loss, and duplication, indicating that WAI isolates represent an evolutionary hotspot while retaining a stable conserved core genome.

Functional annotation further reflected this genomic partitioning. Core orthogroups exhibited nearly complete homology-based annotation, whereas annotation rates declined markedly across accessory compartments, consistent with the rapid evolution of accessory genes. Nevertheless, integration of the deep learning-based predictor DeepFRI substantially improved annotation of lineage-specific genes, assigning functions to 2,153 orthogroups that lacked homology-based annotations and demonstrating that many apparently orphan genes likely perform important metabolic or regulatory functions (Gligorijević et al. 2021). GO enrichment analysis revealed clear functional specialization among pangenome compartments. The core genome was enriched for conserved housekeeping, primary metabolism, carbohydrate utilization, and mycotoxin biosynthesis, consistent with observations in other fungal pathogens and reflecting the conservation of essential metabolic and pathogenicity functions (McCarthy and Fitzpatrick 2019; Alouane et al. 2021; Sweany et al. 2022). In contrast, the shell genome was enriched for regulatory processes, stress responses, intron homing, and nonribosomal peptide biosynthesis, supporting its role in transcriptional plasticity, genome diversification, and secondary metabolism, similar to the accessory genome of *F. graminearum* (Alouane et al. 2021). The cloud genome showed enrichment for chromatin organization, DNA conformational change, stress response, meiotic processes, interspecies interactions, and mitochondria–nucleus signaling, suggesting that this highly dynamic compartment contributes to genome remodeling, environmental adaptation, and virulence, consistent with chromatin-mediated regulation in filamentous fungi and retrograde signaling described in *Candida albicans* (Hans et al. 2019; Aslam et al. 2024).

### 4.3 Virulence repertoire and secondary metabolite biosynthetic networks

The predominance of conserved CAZyme families in the *B. sorokiniana* pangenome supports their central role in virulence and host colonization. Several highly abundant families, including AA9, GH16, GH43, CE4, CE5, PL1, GH47, and GH28, are also key virulence-associated CAZymes in *Fusarium oxysporum*, where they mediate plant cell wall degradation and host invasion (A. Roy et al., 2020). Similar roles have been reported for GH28 during infection by *Calonectria eucalypti* (Liu et al. 2025). The pangenome also revealed extensive functional overlap among CAZymes, effectors, and PHI-associated genes, with 176 CAZyme–PHI, 45 CAZyme–effector, and 51 PHI– effector shared orthogroups. Notably, 65 orthogroups were common to all three categories, and the broader virulence repertoire accounted for 3,722 orthogroups (22% of the pangenome), highlighting an integrated network linking host cell wall degradation, immune modulation, and pathogenic fitness **(Figure S14)**.

Functional analysis of the 65 orthogroups shared by CAZyme, effector, and PHI-base datasets identified a highly conserved virulence backbone in *B. sorokiniana* **(Table S32)**. Sixty-one orthogroups belonged to the core genome and only four to the shell genome, indicating strong evolutionary conservation of this integrated pathogenicity network. These orthogroups were predominantly associated with reduced virulence, hypervirulence, effector (plant avirulence determinant) phenotypes, and loss-of-pathogenicity phenotypes, underscoring their importance for fungal fitness and successful host infection. Consistent with the extracellular infection strategy of *B. sorokiniana*, 61 of the 65 orthogroups encoded apoplastic effectors. Notably, seven orthogroups were simultaneously annotated as CAZymes, apoplastic effectors, and PHI-base effectors (plant avirulence determinants), including AA9 lytic polysaccharide monooxygenases (OG0003022 and OG0007865), GH16 glycoside hydrolases (OG0003654 and OG0009687), a GH12 glycoside hydrolase (OG0011182), a CBM18_GH16-containing protein (OG0005172), and an additional glycoside hydrolase (OG0004592). Except for the softcore GH12 orthogroup (OG0011182), all were core conserved, underscoring their likely roles in host cell wall degradation, oxidative lignocellulose breakdown, immune modulation, and apoplastic colonization during infection.

PHI-based homology and phylogenetic analyses identified conserved host-selective toxins (HSTs) that contribute to virulence diversification in the *B. sorokiniana* accessory genome. OG0012030 comprised a conserved *ToxA*-like effector family (14 copies in 13 strains) sharing 94– 99.3% identity with the *P. tritici-repentis ToxA* reference and moderate similarity (51.5%) to *B. maydis ToxA*, supporting conservation of this hypervirulence-associated effector within Pleosporales. In contrast, OG0014085 represented a *ToxB*-like lineage showing 51.9% identity to *P. tritici-repentis ToxB* in three isolates, indicating diversification of necrotrophic effectors among cereal pathogens (Ciuffetti et al. 2010). Phylogenetic analysis further revealed lineage-specific diversification, including a duplicated and divergent *ToxA* copy in WAI3285, multiple *ToxA* haplotypes in SI, and the coexistence of both *ToxA* and *ToxB* homologs in Yt-6, a toxin combination associated with increased virulence and broader host range in *P. tritici-repentis* (Ciuffetti et al. 2010; Gourlie et al. 2022). The contrasting evolutionary histories of these toxins support distinct origins. The limited sequence variation of *ToxA* is consistent with its recent acquisition through horizontal gene transfer (HGT) among *P. tritici-repentis*, *Parastagonospora nodorum*, and *B. sorokiniana*, followed by localized duplications in strains such as WAI3285 and SI. In contrast, *ToxB* appears to represent an ancient vertically inherited lineage that has diversified through transposon activity, copy number variation, and Starship-like mobile elements, including Icarus, resulting in continued structural evolution across Ascomycota (Gourlie et al. 2022; Hafez et al. 2024) **(Figures S15 and S16)**.

The quantitative variation in BGCs (39–54 clusters per genome) indicates that *B. sorokiniana* possesses a conserved yet flexible secondary metabolome, enabling adaptation to different ecological niches and host genotypes. BiG-SCAPE analysis showed that NRPS clusters from 16 strains were highly similar to the dimethylcoprogen siderophore cluster (BGC0001249) of *Alternaria alternata*. Dimethylcoprogen-mediated iron acquisition is essential for fungal growth, oxidative stress tolerance, and virulence, and disruption of the core *AaNPS6* gene abolishes siderophore production, reducing pathogenicity, conidiation, and melanin biosynthesis (Li et al. 2023; Byers et al. 2025; González et al. 2025). Its widespread conservation across *B. sorokiniana* and diverse Pleosporales, including *B. maydis*, *P. tritici-repentis*, *Alternaria*, *Curvularia*, and *Lasiodiplodia* spp., suggests that iron scavenging is a core adaptation to host nutritional immunity and an evolutionarily conserved virulence strategy (Kim and Dettman 2025). Similarly, an NRPS-like choline-associated BGC (BGC0002276) was conserved in 18 strains, indicating that phospholipid metabolism is another core adaptive trait. Choline metabolism maintains membrane integrity, osmotic balance, and cellular homeostasis (Markham et al. 1993; Hai et al. 2019), while disruption of choline-associated genes impairs growth and virulence in *Botrytis cinerea*, *M. oryzae*, *F. graminearum*, and *Rhizopus microsporus* (Chand Arya et al. 2022; Xu et al. 2023; Carrillo-Marín et al. 2026). Its conservation in *B. sorokiniana* and related pathogens, including *B. oryzae*, *Alternaria alstroemeriae*, and *Elsinoë batatas*, further supports a fundamental role in fungal fitness and host colonization (Xu et al. 2024; Zhang et al. 2025; Hue et al. 2025).

The T1PKS 4-chloropinselin cluster (BGC0002726) was also conserved in 18 strains, encoding chlorinated chromones and xanthones such as 4-chloropinselin, pinselin, chloromonilinic acids, and 4-hydroxyvertixanthone, metabolites associated with antimicrobial activity, oxidative stress tolerance, melanization, ecological competition, and pathogenic fitness (Han et al. 2019). Similarly, metabolomic investigations in *Alternaria sonchi* link T1PKS with antimicrobial, insecticidal, and phytotoxic activities, supporting their ecological role in microbial competition and environmental fitness (Berestetskiy et al. 2019; Dalinova et al. 2020; Yu et al. 2020). The terpestacin BGC (BGC0002245) was conserved in all 19 strains, establishing it as a core metabolic feature of the species. The tpcA–D pathway synthesizes terpestacin through a coordinated enzymatic cascade (Narita et al. 2018; Sun et al. 2025), and homologous clusters occur in *B. maydis*, *Fusarium proliferatum*, *A. alstroemeriae*, and *Arthrinium* spp. (Ćeranić et al. 2021; Sun et al. 2025; Zhang et al. 2025). In addition to its role in phytotoxicity, terpestacin promotes host colonization by inducing chloroplast damage, thylakoid disorganization, growth inhibition, and plant senescence, while also exhibiting antiangiogenic activity and broad-spectrum antifungal properties that enhance ecological competitiveness (Masi et al. 2018, 2022; Ćeranić et al. 2021). Its universal conservation, therefore, suggests that this cluster is a core adaptive trait supporting both pathogenicity and environmental persistence.

In contrast, the prehelminthosporol/sativene cluster (BGC0003041) was detected in only 11 of the 19 strains, indicating that it represents an accessory pathogenicity-associated BGC. This cluster synthesizes sativene-derived sesquiterpenoids, including prehelminthosporol, dihydroprehelminthosporol, and prehelminthosporolactone, which function as host-selective phytotoxins affecting wheat, rice, and weed species (Xu et al. 2021; Zhang et al. 2024; He et al. 2024). Sesquiterpenoid production correlates with increased virulence in *B. sorokiniana* and is conserved in related necrotrophs such as *B. setariae*, *B. zeicola*, *B. victoriae*, *Drechslera*, and *Cochliobolus* (Apoga et al. 2002). Its restricted distribution suggests lineage-specific specialization, with retaining strains potentially gaining enhanced phytotoxicity and host adaptation, whereas strains lacking the cluster may rely on alternative virulence strategies. Thus, unlike the universally conserved terpestacin locus, BGC0003041 represents an accessory metabolic innovation driving pathotypic diversification and is an attractive target for future functional, transcriptomic, and metabolomic studies.

### 4.4 Evolutionary pressures and gene family dynamics

CAFE analysis showed that gene family evolution in *B. sorokiniana* was dominated by contraction rather than expansion. This bias suggests ongoing genome streamlining accompanying host specialization, a pattern previously reported in *Calonectria* spp. and other Hypocrealean fungi (Zhang et al. 2018; Rogers et al. 2022). Accordingly, strains ND93-1, BRIP10943a, WAI2432, WAI3384, and WAI2411 exhibited extensive lineage-specific gene loss. In contrast, only one orthogroup, OG0000151, encoding an aldehyde dehydrogenase (ALDH), showed expansion exclusively in strain D2 (11 copies versus one copy in all other strains) along with signatures of positive selection. ALDHs detoxify reactive aldehydes generated during host-induced oxidative stress and are required for fungal growth and virulence in *F. graminearum*, *M. oryzae*, and *F. verticillioides* (Abdul et al. 2018; Tang et al. 2023; Ren et al. 2024). Similar infection-associated induction has been reported in *Bipolaris cookei*, suggesting that expansion of OG0000151 may enhance oxidative stress tolerance and contribute to the increased virulence previously reported for strain D2 (Yadav et al. 2022).

Rapidly evolving gene families were enriched for proteins involved in self/non-self-recognition, secondary metabolism, and environmental adaptation. Extensive copy number variation was observed among fungal NLR proteins, including NACHT-WD40 (OG0000000, OG0000003, OG0012129, OG0012876), NACHT-Ankyrin (OG0000002, OG0000021, OG0013420), and HET/HET-E-like proteins (OG0000010, OG0000040, OG0013028, OG0013405, OG0013772) **(Figure S17)**. These multidomain receptors regulate vegetative incompatibility and immune-like signaling, and their rapid diversification through domain reshuffling is a characteristic feature of the evolution of the fungal Nucleotide-binding domain and Leucine-rich Repeat containing (NLR) genes (Marino and Brodsky 2023; Bonometti et al. 2025). Similar copy number variation was also observed in NRPS- and PKS-associated orthogroups (OG0000004, OG0000005, OG0000013, OG0000950, OG0001000, OG0013042, OG0013565, OG0000078, OG0012952, OG0012980), the siderophore synthetase OG0013251, and multiple cytochrome P450 families (OG0012948, OG0013003, OG0000095, OG0013054, OG0011911), reflecting continual remodeling of secondary metabolite production, iron acquisition, xenobiotic detoxification, and host adaptation (Sav et al. 2018; Carroll and Moore 2018).

Positive selection analysis further identified 501 core orthogroups evolving under strong adaptive pressure, indicating continuous optimization of genes associated with pathogenicity and environmental fitness. Adaptive evolution preferentially targeted regulatory proteins, including protein kinases, Zn(II)₂Cys₆ and MYB transcription factors, and Rho-family GTPases that coordinate infection-related signaling pathways such as MAPK and cAMP-PKA cascades (An et al. 2015; Fu et al. 2018; John et al. 2024). Selection also acted on nutrient acquisition and stress-response systems, including ammonium and sugar transporters, copper and calcium transporters, aquaporins, and ABC/MFS efflux pumps, reflecting adaptation to host nutrient limitation, oxidative stress, and chemical defenses (O’Mara et al. 2023; Palos-Fernández et al. 2024).

Several positively selected orthogroups encoded proteins central to oxidative stress tolerance and cellular detoxification, including ALDHs, Cu/Zn superoxide dismutases, GMC (glucose-methanol-choline) oxidoreductases, amine oxidases, pyridine nucleotide-disulfide oxidoreductases, and homogentisate dioxygenases, all of which contribute to detoxification, redox homeostasis, melanin biosynthesis, and protection against host-derived reactive oxygen species (Yuste-Lisbona et al. 2020; Binder et al. 2020; Singh et al. 2021). Likewise, positive selection also targets multiple secreted virulence factors, including GH, GT, and CBM-family CAZymes, reflecting continual optimization of plant cell wall degradation and evasion of host pattern-recognition receptors (Hage and Rosso 2021). Several effector-associated proteins, including a Class II hydrophobin, a GDT1-like (UPF0016) transporter, a copper radical oxidase, and a SUN-family β-glucosidase, also exhibited strong adaptive signatures. These proteins regulate host adhesion, ion homeostasis, stress tolerance, and cell wall remodeling and are essential for virulence in several fungal pathogens (Aguiar et al. 2021; Wu et al. 2022; Meng et al. 2025). These results indicate that adaptive evolution in *B. sorokiniana* primarily targets regulatory networks, nutrient acquisition, oxidative stress responses, secondary metabolism, and secreted pathogenicity factors, enabling rapid adaptation while maintaining a conserved pathogenic core.

### 4.5 One-compartment genome architecture with localized adaptive gene enrichment

The genome architecture of *B. sorokiniana* resembles the one-compartment organization reported for the wheat pathogen *P. tritici-repentis*, where intergenic distance distributions form a continuous landscape rather than distinct gene-dense and gene-sparse compartments (Gourlie et al. 2022). Nevertheless, effector genes, CAZymes, and rapidly evolving CAFE-associated gene families were significantly enriched in relatively gene-sparse genomic neighborhoods, suggesting localized structural environments that may facilitate adaptation and host interaction. In contrast, PHI-associated and positively selected genes showed distributions similar to the genomic background. These findings indicate that adaptive and pathogenicity-related functions in *B. sorokiniana* are associated with regions of reduced gene density despite the absence of discrete genome compartments.

### 4.6 Topological architecture and modular organization of the core interactome

The highly connected core PPI network indicates that virulence functions are embedded within the conserved molecular framework of *B. sorokiniana*, with CAZymes, effectors, and PHI-associated proteins frequently occupying multifunctional hub nodes. Louvain clustering further resolved the interactome into housekeeping modules that are tightly integrated with stress-response and pathogenicity pathways, ensuring cellular robustness during host infection.

#### 4.6.1 Community 1: The proteostatic and genome maintenance core

Community 1 represents the regulatory backbone of the core interactome, integrating transcription, ubiquitin-mediated proteolysis, RNA processing, and genome maintenance. Its central hub, OG0002112 (Rrn7), is a conserved RNA polymerase I initiation factor (composite centrality = 0.87) essential for rRNA synthesis and transcriptional regulation (Knutson and Hahn 2011). A defining feature of this module is the enrichment of 20S and 19S proteasome components, including OG0003506 (SCL1), OG0006019 (PUP2), OG0005314 (PRE8), OG0003005 (PRE6), OG0006460 (RPN10), OG0006021 (RPN8), and OG0002010 (RPN2), highlighting proteostasis as a central network function. Since the disruption of proteasome integrity compromises fungal development, stress tolerance, and infection, PHI-base evidence linking RPN11 (OG0005284) and RPN2 (OG0002010) to reduced virulence in *Alternaria alternata* and *Verticillium dahliae*, respectively, underscores the importance of this machinery for pathogenicity (Wang et al. 2018; Ren et al. 2022).

Proteasome components are closely connected with AAA+ ATPases OG0009610 (RPT5), OG0002715 (RPT2), OG0006920 (RPT3), OG0001619 (RPT1), and OG0009305 (AFG2), whose disruption causes lethal phenotypes in *F. graminearum*. Notably, the 26S regulatory ATPase OG0001924 also exhibits positive selection, suggesting adaptive optimization of proteostasis under host-induced proteotoxic stress. This proteolytic machinery interfaces with RNA polymerase subunits OG0010227 (RPO26), OG0001335 (RPB3), OG0001532 (RPB2), OG0001538 (RPO21), and OG0003562 (RPB8) and ubiquitin–ribosomal fusion proteins (OG0013035 and OG0002111) to coordinate transcription, translation, and stress-responsive protein turnover, consistent with reduced-virulence phenotypes reported in *Cryptococcus neoformans* (Zhao et al. 2020). Community 1 also contains a conserved genome maintenance network comprising OG0000501 (MEC1), OG0007982 (RFA1), OG0009078 (SSB2), OG0010004 (CDC6), OG0008087 (CDC7), OG0003911 (RVB1), and OG0007108 (HTA1). Together, these checkpoint, replication, and chromatin-remodeling factors preserve genome stability while facilitating adaptive genetic variation under oxidative stress and fungicide pressure, providing the genomic plasticity required for pathotypic diversification (Shor et al. 2020; Huang and Cook 2022).

#### 4.6.2 Community 2: The ribosome biogenesis and cellular proliferation hub

Community 2 forms a highly interconnected nucleolar ribosome biogenesis module, enriched for ribonucleoprotein (rRNP) assembly, small-subunit (SSU) processome, and 60S ribosomal maturation pathways that sustain the high translational demands of fungal growth and host colonization. Defects in preribosomal components, including FgEXOSC1, the Lsm8 exosome, and MoRRP8, consistently impair conidiation, appressorium formation, mycotoxin production, and virulence, emphasizing the essential role of ribosome biogenesis in pathogenicity (Yuan et al. 2023; Kim et al. 2023).

The central hub of this community is OG0003241 (PWP2; composite centrality = 0.920), a conserved WD40-repeat protein required for early 90S pre-ribosome assembly and cell-cycle progression (Shafaatian et al. 1996). PHI-base links its *Alternaria panax* homolog (ApWD40a) to reduced virulence, sporulation, and toxin production (Lan et al. 2025). PWP2 functions with UTP4 (OG0002112), UTP10 (OG0000984), and UTP15 (OG0005074) of the UtpA complex to organize the 5′ ETS during early SSU assembly . The U3 snoRNP component MPP10 (OG0001701), a key regulator of 18S rRNA cleavage, showed positive selection, suggesting adaptive fine-tuning of early translational machinery under host-derived stress (Dunbar et al. 1997; Sá-Moura et al. 2017). Additional SSU factors, including NOP14 (OG0006533), KRR1 (OG0008864), Utp11 (OG0005942), and RRP5 (OG0001839), coordinate 18S rRNA maturation and 40S assembly (Kirsch et al. 2020; Yan et al. 2024), while KRR1 is classified as a lethal PHI-base determinant in *Aspergillus fumigatus* (Hu et al. 2007).

The module also coordinates 60S ribosome biogenesis through the conserved NOP7 subcomplex comprising ERB1 (OG0009127), YTM1 (OG0000216), and NOP7/PES1 (OG0010135), which mediate 5.8S and 25S rRNA maturation (Tang et al. 2008). PHI-base links YTM1 and NOP7/PES1 to severe virulence defects, whereas MRT4 (OG0008595) and Nop53 (OG0003796) are essential for appressorium formation, lesion development, and successful host colonization (Yang et al. 2023; Li et al. 2025). Translation fidelity is further supported by conserved rRNA-modifying enzymes (SPB1, DIM1, NOP1/Fibrillarin, NAT10, CBF5, NOP2), Brix-domain proteins RPF1 (OG0002966) and RPF2 (OG0006819), and RNA helicases SPB4 (OG0002475), PRP43 (OG0000493), and MTR4 (OG0007251), which drive preribosomal remodeling and RNA quality control (Maekawa et al. 2018; Vanden Broeck and Klinge 2022). Their association with reduced virulence in *M. oryzae* (PHI:124978) highlights the importance of post-transcriptional RNA surveillance for pathogenic fitness (Shi et al. 2019). Together, these findings identify Community 2 as a highly conserved ribosome biogenesis hub that supports fungal growth, translation, toxin production, and virulence in *B. sorokiniana*. Because ribosome assembly is indispensable for fungal viability, this module represents an attractive antifungal target, as demonstrated by the antiproliferative activity of benzisothiazolinone (BIT) and the ability of thiosemicarbazone NSC319726 to inhibit fungal growth by blocking rRNA maturation and small-subunit assembly (Liu et al. 2026).

#### 4.6.3 Community 3: The bioenergetic engine and metabolic integration axis

Community 3 represents the bioenergetic and metabolic core of the *B. sorokiniana* interactome, integrating mitochondrial respiration, ATP synthesis, organellar translation, and central carbon metabolism. Its central hub, OG0010829 (RPS3), is a conserved ribosomal protein that not only supports translation but also ribosome quality control through stalled-ribosome surveillance and no-go mRNA decay (Simms et al. 2018). Together with mitochondrial ribosomal proteins RML2 (OG0008970), MRPL8 (OG0007313), MRPL10 (OG0004711), MRPS9 (OG0008230), and MRPS16 (OG0007687), it maintains mitochondrial translation and respiratory competence, with RML2 playing an essential role in ribosome assembly and peptidyl-transferase activity (Diedrich et al. 2000; Korovesi et al. 2018).

A prominent feature of this community is the enrichment of the F₁F₀-ATP synthase (ATP1/OG0001612, ATP2/OG0000888, ATP3/OG0006682, ATP5/OG0004085, ATP7/OG0001494, ATP16/OG0010463, ATP9/OG0002210) and electron transport chain complexes (RIP1, CYC1, QCR7, COX4, PET9), which sustain oxidative phosphorylation and ATP production. PHI-base links ATP1 to loss of pathogenicity in *C. albicans* and ATP16 to reduced virulence, demonstrating that mitochondrial energy metabolism is indispensable for fungal infection. Likewise, respiratory complexes associated with reduced-virulence phenotypes emphasize that maintaining proton motive force is essential for supporting the energy-intensive processes required during host colonization. This respiratory network is tightly coupled with glycolytic and nucleotide metabolic enzymes, including PYK1 (OG0010147), GAPDH (OG0001577), PGK1 (OG0001876), ENO1 (OG0002002), URA7 (OG001367), and NDK1 (OG0003136), whose disruption consistently reduces virulence or restricts *in planta* growth. Notably, the alpha-enolase, ENO1, is under strong positive selection despite its conserved metabolic role. Beyond glycolysis, it functions as a moonlighting surface protein that mediates host adhesion through interactions with fibronectin, vitronectin, and plasminogen, promotes immune evasion, and represents a promising antifungal target (Li et al. 2022; Langenhorst et al. 2023), suggesting adaptive optimization of both metabolism and host interaction. Community 3 also integrates mitochondrial quality-control components, including the Hsp70 chaperone SSC1 (OG0001834) and prohibitin PHB1 (OG0008718), which preserve membrane integrity, mitigate oxidative stress, and facilitate respiratory complex assembly. Collectively, these findings establish Community 3 as a conserved bioenergetic hub that links mitochondrial homeostasis, metabolic flexibility, and adaptive evolution, thereby ensuring an energy supply for successful pathogenicity.

#### 4.6.4 Community 4: The cytosolic translational engine and elongation scaffold

Community 4 was driven by cytosolic translation pathways, ribosome organization, and translational initiation. This module was dominated by highly central core orthogroups encoding structural constituents of both the small (40S) and large (60S) ribosomal subunits, explicitly including *RPS4*, *RPS5*, *RPS6*, *RPS9*, *RPS13*, *RPS15*, *RPS19*, *RPS20*, *RPL2*, *RPL3*, *RPL5*, *RPL11*, *RPL17*, *RPL20*, *RPL27*, *RPL35*, and *RPP0*. As constituents of the translational apparatus, these proteins are under rigid evolutionary constraints to ensure flawless mRNA decoding, tRNA positioning, and peptide elongation, making this module a non-redundant determinant of vegetative viability and pathotypic fitness. PHI-base links this translational core to virulence. RPS4 (OG0010340; PHI:6076) is a validated virulence determinant in *C. albicans* (Lu et al. 2016), while the translation initiation factor Eif2α (OG0002193; PHI:1582) regulates selective protein synthesis during host-induced stress. Although RPL2 (PHI:1584) and RPS23 (PHI:1604) show no virulence phenotype individually in *F. graminearum*, their central network positions indicate that maintenance of global translational capacity is critical for cellular homeostasis (Son et al. 2011).

The module was further anchored by the ribosomal stalk protein RPP0 (P0), which recruits translation elongation factors and governs elongation efficiency. Alterations in RPP0 impair translation while conferring resistance to the antifungal sordarin, highlighting ribosomal stalk proteins as key determinants of fungal fitness (Gómez-Lorenzo and García-Bustos 1998). Likewise, disruption of the PlCYP5–PlRPS15 assembly complex abolishes ribosome biogenesis and vegetative growth (Mo et al. 2021), whereas D-limonene suppresses fungal proliferation by coordinately repressing ribosomal genes, including RPS4, RPS9, and RPS3 in *F. proliferatum* (Zhou et al. 2025). As proper ribosomal stalk assembly and translation elongation kinetics dictate the rapid production of effectors, CAZymes, and stress-tolerance enzymes, this consolidated protein synthesis engine represents a prime target for small-molecule antifungal interventions aimed at breaking down the pathogen’s adaptive capacity during wheat infection. We also developed an interactive R Shiny dashboard (*Bipolaris PanExplorer:* https://pangenomebipolaris.shinyapps.io/bipolaris_panexplorer/) to provide user-friendly access to the pangenome datasets and analytical results, facilitating exploration and reuse by the research community.

## 5. Conclusions

This study presents the first comprehensive pangenome-scale analysis of *B. sorokiniana*, integrating comparative genomics, evolutionary analyses, and protein interaction networks to elucidate the genomic basis of adaptation and pathogenicity. Although the reference genome (ND90Pr) contains 12,210 predicted protein-coding genes, the species-wide pangenome comprises 16,981 orthogroups, revealing more extensive genetic diversity than any single isolate.

The open pangenome reflects continuous genome evolution driven by transposable element activity, structural variation, and gene gain and loss events. While the conserved core genome primarily encodes essential cellular and metabolic functions, the accessory genome serves as a reservoir of genes involved in virulence, host adaptation, and environmental responsiveness. This genomic plasticity is particularly evident in the recently sequenced WAI lineage, which exhibits accelerated gene turnover and expansion of shell and cloud gene families. Integration of positive selection, gene family evolution, PHI-base, CAZyme, effector, and network analyses identified numerous candidate genes and pathways underlying virulence, stress adaptation, and evolutionary fitness. The discovery of *ToxA*- and *ToxB*-like effector homologs, together with evidence of lineage-specific diversification and duplication, further highlights the dynamic evolution of necrotrophic effectors within *B. sorokiniana* populations. The pangenome resource generated in this study provides a comprehensive framework for future functional genomics investigations, facilitating the identification of key virulence determinants and supporting the development of durable disease-management strategies and resistant wheat cultivars through targeted breeding and molecular surveillance.

## Acknowledgements

This research was funded by Agrihub, Indian Institute of Technology Indore, and the Council of Scientific and Industrial Research (CSIR), India. AKS acknowledges AcSIR, India, for the Integrated Dual-Degree Program (IDDP) and the research fellowship for IDDP under the CSIR-GATE-JRF program (Award No.: 31/ GATE/11(55)/2025-EMR-I).

## Author contributions statement

**Anand Kumar Shukla**: Conceptualization, Resources, Methodology, Investigation, Software, Formal analysis, Data curation, Visualization, Writing-Original draft preparation, Editing; **Narendra Kadoo**: Resources, Supervision, Writing-Reviewing and Editing, Funding acquisition, Project administration. All authors have contributed to the writing and substantial revisions of the manuscript and have read and approved the final version.

## Declaration of Generative AI and AI-assisted technologies in the writing process

During the preparation of this work, the authors used ChatGPT to improve the readability of the text. After using this tool/service, the authors reviewed and edited the content as needed and take full responsibility for the content of the publication.

## Data availability

The interactive *Bipolaris PanExplorer* R Shiny dashboard developed in this study is freely available at https://pangenomebipolaris.shinyapps.io/bipolaris_panexplorer/ and enables exploration of the pangenome. The dashboard enables interactive exploration of the *B. sorokiniana* pangenome, including genome assembly statistics, orthogroup composition, functional annotations, virulence-associated genes, gene family evolution, positive selection analyses, protein–protein interaction networks, and downloadable datasets. The complete source code and associated datasets are publicly available at GitHub (https://github.com/Anand41352/Bipolaris-PanExplorer/tree/main). The genome assemblies, gene annotation (GFF3) files, predicted protein sequences, and coding DNA sequences (CDS) for all 19 *B. sorokiniana* strains generated in this study have been uploaded to Zenodo and are publicly available (https://doi.org/10.5281/zenodo.21263238).

## Supplementary Figure Legends

**Figure S1:** Overlap of functional annotation across multiple annotation sources. UpSet plot showing the overlap of Gene Ontology (GO) annotations for predicted orthogroups across four annotation sources: InterProScan, EggNOG, SwissProt, and DeepFRI. Each set represents proteins annotated by a given source, while the intersection bars indicate the number of proteins sharing annotations between one or more sources. The matrix layout below the bars specifies the combination of sources contributing to each intersection. The left-hand bar plot shows the total number of proteins annotated by each source.

**Figure S2:** Gene Ontology (GO) enrichment analysis of pangenome orthogroup categories (Core, Shell, and Cloud) in *Bipolaris sorokiniana*.

**Figure S3:** Comparative clinker visualization of the choline biosynthetic gene cluster (BGC0002276) identified across *Bipolaris sorokiniana* genomes. Conserved gene organization and synteny with the corresponding MIBiG reference cluster illustrate the high conservation of this NRPS-like biosynthetic locus, which was detected in 18 strains.

**Figure S4:** Comparative clinker visualization of the 4-chloropinselin biosynthetic gene cluster (BGC0002726) across *B. sorokiniana* genomes. The conserved arrangement of biosynthetic genes shows strong synteny with the MIBiG reference cluster, supporting the widespread conservation of this T1PKS-associated secondary metabolite pathway across 18 strains.

**Figure S5:** Comparative clinker visualization of the terpestacin biosynthetic gene cluster (BGC0002245) identified in *B. sorokiniana*. Conserved gene content and organization relative to the MIBiG reference cluster indicate that this terpene biosynthetic pathway is completely conserved across all 19 analyzed genomes.

**Figure S6:** Comparative clinker visualization of the prehelminthosporol/sativene biosynthetic gene cluster (BGC0003041) across *B. sorokiniana* genomes. Conserved synteny with the reference cluster demonstrates the presence of this terpene-associated biosynthetic pathway in 11 strains, highlighting lineage-specific conservation of genes implicated in sesquiterpene biosynthesis and pathogenic adaptation.

**Figure S7:** Gene Ontology (GO) enrichment analysis of significantly expanded and contracted orthogroups identified by CAFE.

**Figure S8:** Genome compartmentalization analysis of the chromosome-level *Bipolaris sorokiniana* LK93 genome. Distribution of genes based on 5′ and 3′ flanking intergenic regions (FIRs) demonstrates a one-compartment genome architecture, lacking distinct gene-dense and gene-sparse compartments.

**Figure S9:** Global STRING-derived protein–protein interaction (PPI) network of the core proteome in *Bipolaris sorokiniana*. The figure represents the large-scale protein–protein interaction (PPI) network constructed from conserved core orthogroups identified across multiple *B. sorokiniana* genomes using the STRING database. The complete raw network consisted of 6,539 nodes and 224,198 edges, representing predicted and experimentally supported functional protein associations. Extraction of the giant connected component retained 6,450 nodes and 224,128 edges, indicating extensive connectivity among conserved proteins within the fungal core proteome. Nodes represent core orthogroups/proteins, while edges denote functional interactions inferred from STRING evidence channels.

**Figure S10:** Gene Ontology (GO) enrichment analysis of Biological Processes for Community 2 (C2) identified from the STRING-derived protein–protein interaction (PPI) network of the top 10% core proteome in *Bipolaris sorokiniana*.

**Figure S11:** Gene Ontology (GO) enrichment analysis of Biological Processes for Community 4 (C4) identified from the STRING-derived protein–protein interaction (PPI) network of the top 10% core proteome in *Bipolaris sorokiniana*.

**Figure S12:** Gene Ontology (GO) enrichment analysis of Biological Processes for Community 1 (C1) identified from the STRING-derived protein–protein interaction (PPI) network of the top 10% core proteome in *Bipolaris sorokiniana*.

**Figure S13:** Gene Ontology (GO) enrichment analysis of Biological Processes for Community 3 (C3) identified from the STRING-derived protein–protein interaction (PPI) network of the top 10% core proteome in *Bipolaris sorokiniana*.

**Figure S14:** Venn diagram illustrating the overlap among orthogroups associated with carbohydrate-active enzymes (CAZymes), pathogen–host interaction (PHI-base) homologs, predicted effector proteins, genes under positive selection, and gene families identified as significantly expanded or contracted by CAFE analysis. The diagram highlights shared and unique orthogroups across these functional and evolutionary categories, providing an overview of candidate genes potentially involved in pathogenicity, host adaptation, and genome evolution in the pangenome.

**Figure S15:** Multiple sequence alignment of host-selective effector homologs identified in the *Bipolaris sorokiniana* pangenome. (A) Alignment of OG0012030-encoded ToxA homologs against the ToxA reference sequence from Pyrenophora tritici-repentis, revealing highly conserved homologs with sequence identities reaching ∼99%, consistent with strong conservation across multiple *B. sorokiniana isolates.* (B) Alignment of OG0012030 homologs against the ToxA sequence from Bipolaris maydis, showing substantially lower sequence similarity (∼51%), indicative of a more divergent ToxA-like lineage. (C) Alignment of OG0014085-encoded ToxB homologs against the ToxB reference sequence from Pyrenophora tritici-repentis. ToxB homologs were detected in only three *B. sorokiniana isolates a*nd exhibited moderate sequence similarity (∼52%), suggesting a restricted distribution and greater evolutionary divergence compared with Ptr-like ToxA homologs. Conserved amino acid residues are highlighted according to sequence similarity, and gaps introduced during alignment are represented by dashes.

**Figure S16:** Phylogenetic distribution and copy number variation of necrotrophic effectors ToxA and ToxB across pangenomic lineages of *Bipolaris sorokiniana*. Maximum likelihood phylogenetic tree constructed using IQ-TREE (version 2.0.7) from aligned amino acid sequences of ToxA and ToxB homologs across fungal pathogen strains. Clades are color-coded based on pangenomic orthogroup annotations: OG0012030 (ToxA Clade I) is shown in vermillion, OG0012030 (ToxA Clade II) in orange/yellow, and OG0014085 (ToxB Clade) in ocean blue. Core reference strains (*Pyrenophora tritici-repentis* and *Bipolaris maydis*) are highlighted in bold. Duplicated copy of ToxA in strain WAI3285 is highlighted in bold-italic and annotated with a red asterisk (∗), indicating a structural duplication event containing two distinct paralogous copies of the *ToxA* locus (WAI3285_05990 and WAI3285_06714)

**Figure S17:** Gene family expansion and contraction dynamics of pathogenicity- and defense-associated gene repertoires across 19 *Bipolaris sorokiniana* genomes. Heatmap showing copy-number variation for 30 significantly evolving orthogroups identified by CAFE analysis. Orthogroups were grouped into four biologically relevant functional classes: NACHT/NLR-associated defense proteins (13 orthogroups), non-ribosomal peptide synthetases (NRPS; 8 orthogroups), polyketide synthases (PKS; 3 orthogroups), and cytochrome P450 monooxygenases (CYP450; 6 orthogroups). Cell values represent actual gene copy numbers in each genome, while color intensity indicates relative abundance categories derived from within-orthogroup quantile normalization (Rare, Low, Moderate, and Expanded), enabling comparison of expansion patterns among strains despite differences in absolute copy numbers. Genomes were hierarchically clustered using Ward’s method based on orthogroup copy number profiles. Left-side annotation tracks indicate pangenome classification (Core, Softcore, and Shell), CAFE evolutionary dynamics (Both Expansion and Contraction or Contraction Only), and eggNOG COG functional categories. COG category abbreviations: A, RNA processing and modification; D, cell cycle control, cell division and chromosome partitioning; I, lipid transport and metabolism; K, transcription; M, cell wall/membrane/envelope biogenesis; Q, secondary metabolite biosynthesis, transport and catabolism; S, function unknown; T, signal transduction mechanisms; B, Chromatin structure and dynamics.

## Supplementary Table Legends

**Table S1:** Summary of the *Bipolaris sorokiniana* genomes used for pangenome construction

**Table S2:** QUAST-based genome assembly quality metrics, contiguity statistics, and sequence characteristics of the 19 *B. sorokiniana* genomes analyzed in this study

**Table S3:** BUSCO completeness statistics, including complete, single-copy, duplicated, fragmented, and missing orthologs, for the 19 *B. sorokiniana* genome assemblies

**Table S4A:** *De novo* repeat annotation and genome-wide repeat composition of the 19 *B. sorokiniana* genomes identified using RepeatModeler and RepeatMasker

**Table S4B:** Homology-based repeat landscape and transposable element composition of the 19 *B. sorokiniana* genomes

**Table S5:** Summary of repeat content identified by de novo and homology-based repeat masking across the 19 *B. sorokiniana* genomes, including genome size, masked bases, and total repeat percentages.

**Table S6:** Summary of predicted protein-coding genes and high-confidence gene models (AED < 0.5) across the 19 *B. sorokiniana* genomes.

**Table S7:** Summary of gene prediction and evidence support across the 19 *B. sorokiniana* genomes

**Table S8:** Orthogroup composition across the 19 *B. sorokiniana* genomes. The table lists all 16981 orthogroups identified by pangenome analysis, their pangenome category (Core, SoftCore, Shell, or Cloud), and the number of strains in which each orthogroup is present (presence count). For each orthogroup, the corresponding gene IDs from all 19 genomes are provided.

**Table S9:** Distribution of orthogroups across pangenome categories in the 19 *B. sorokiniana* genomes

**Table S10:** NR database annotation of representative orthogroups identified in the *B. sorokiniana* pangenome. For each orthogroup, the longest protein sequence was selected as the representative sequence and annotated against the NCBI non-redundant (NR) protein database. The table includes the pangenome category (Core, SoftCore, Shell, or Cloud), representative orthogroup gene ID, best-matching NR protein accession, percent sequence identity, alignment length, E-value, bit score, and the corresponding functional annotation. This approach assigns a representative functional role to each orthogroup based on its longest protein sequence.

**Table S11:** eggNOG functional annotation of representative orthogroups from the pangenome. For each orthogroup, the longest protein sequence was selected as the representative sequence and queried against the eggNOG database.

**Table S12:** InterPro domain annotation of representative pangenome orthogroups. For each orthogroup, the longest protein sequence was selected as the representative for InterProScan analysis.

**Table S13:** Functional annotation of representative orthogroups using the UniProtKB/Swiss-Prot database. For each orthogroup, the longest representative protein sequence was queried against the curated Swiss-Prot database using sequence similarity searches.

**Table S14:** DeepFRI-predicted Gene Ontology (GO) annotations of pangenome genes. The table lists representative genes from the pangenome along with their predicted Gene Ontology (GO) terms generated using DeepFRI.

**Table S15:** Distribution of functionally annotated orthogroups across pangenome categories based on NRDB, eggNOG, InterProScan, Swiss-Prot, and DeepFRI annotations

**Table S16:** Distribution of carbohydrate-active enzymes (CAZymes) across pangenome compartments in individual strains

**Table S17:** Distribution of CAZyme subclasses across the genomes of 19 *B. sorokiniana strains*

**Table S18:** Strain-wise distribution of secreted proteins and predicted effector classes identified using SignalP, TargetP, DeepLoc, and EffectorP

**Table S19:** Predicted effector repertoire showing effector type, orthogroup, and pangenome compartment

**Table S20:** Comparative distribution of pathogen–host interaction (PHI)-base phenotype classes among *B. sorokiniana strains*

**Table S21:** Distribution of biosynthetic gene cluster (BGC) classes across *B. sorokiniana strains*

**Table S22:** BiG-SCAPE-based comparison of biosynthetic gene clusters showing similarity metrics and network relationships

**Table S23:** Clustering of biosynthetic gene clusters into gene cluster families (GCFs) based on BiG-SCAPE similarity analysis

**Table S24:** Orthogroups showing significant gene family expansion or contraction identified by CAFE analysis

**Table S25:** Gene Family Expansion and Contraction Dynamics Inferred by CAFE Analysis

**Table S26:** Functional annotation of orthogroups exhibiting significant expansion and contraction identified by CAFE analysis

**Table S27:** Selection signatures detected in orthogroups using HyPhy evolutionary analyses

**Table S28:** Functional and pathogenicity-related annotations of orthogroups under strong positive selection

**Table S29:** Genome compartmentalization analysis of functional and evolutionary gene categories in *B. sorokiniana* strain LK93 based on flanking intergenic region (FIR) distributions

**Table S30:** Network topology metrics and functional annotations of orthogroups in the core protein–protein interaction (PPI) network

**Table S31:** Top 10% most central orthogroups and hub genes (top 1% of nodes) in the core protein–protein interaction network

**Table S32:** Overlapping orthogroups shared among predicted effectors, CAZymes, and PHI-base homologs in the *B. sorokiniana* pangenome

## References

Abdul W, Aliyu SR, Lin L, et al (2018) Family-Four Aldehyde Dehydrogenases Play an Indispensable Role in the Pathogenesis of Magnaporthe oryzae. Front Plant Sci 9:. 10.3389/fpls.2018.00980

Aggarwal N, Rathore M, Shakeel M, et al (2025) First Report of Bipolaris sorokiniana Causing Spot Blotch on Wheat (Triticum aestivum) from the Kashmir Valley of Western Himalayas, India. Plant Disease 109:2206. 10.1094/PDIS-03-25-0697-PDN

Aguiar M, Orasch T, Misslinger M, et al (2021) The Siderophore Transporters Sit1 and Sit2 Are Essential for Utilization of Ferrichrome-, Ferrioxamine- and Coprogen-Type Siderophores in Aspergillus fumigatus. Journal of Fungi 7:768. 10.3390/jof7090768

Alexa A, Rahnenführer J (2009) Gene set enrichment analysis with topGO. Bioconductor improv 27:776

Almagro Armenteros JJ, Sønderby CK, Sønderby SK, et al (2017) DeepLoc: prediction of protein subcellular localization using deep learning. Bioinformatics 33:3387–3395

Alouane T, Rimbert H, Bormann J, et al (2021) Comparative Genomics of Eight Fusarium graminearum Strains with Contrasting Aggressiveness Reveals an Expanded Open Pangenome and Extended Effector Content Signatures. Int J Mol Sci 22:6257. 10.3390/ijms22126257

Al-Sadi AM (2021) Bipolaris sorokiniana-Induced Black Point, Common Root Rot, and Spot Blotch Diseases of Wheat: A Review. Front Cell Infect Microbiol 11:. 10.3389/fcimb.2021.584899

An B, Li B, Qin G, Tian S (2015) Function of small GTPase Rho3 in regulating growth, conidiation and virulence of *Botrytis cinerea*. Fungal Genetics and Biology 75:46–55. 10.1016/j.fgb.2015.01.007

Apoga D, Åkesson H, Jansson H-B, Odham G (2002) Relationship Between Production of the Phytotoxin Prehelminthosporol and Virulence in Isolates of the Plant Pathogenic Fungus Bipolaris sorokiniana. European Journal of Plant Pathology 108:519–526. 10.1023/A:1019976403391

Aslam HMU, Chikh-Ali M, Zhou X-G, et al (2024) Epigenetic modulation of fungal pathogens: a focus on Magnaporthe oryzae. Front Microbiol 15:1463987. 10.3389/fmicb.2024.1463987

Badet T, Oggenfuss U, Abraham L, et al (2020) A 19-isolate reference-quality global pangenome for the fungal wheat pathogen Zymoseptoria tritici. BMC Biol 18:12. 10.1186/s12915-020-0744-3

Basak P, Kashyap N, Gurjar MS, et al (2025) Recent Advances in Spot Blotch of Barley and Its Future Perspectives. Plant Pathology 74:2579–2597. 10.1111/ppa.70057

Berestetskiy AO, Dalinova AA, Volosatova NS (2019) Metabolite Profiles and Biological Activity of Extracts from Alternaria sonchi S-102 Culture Grown by Different Fermentation Methods. Appl Biochem Microbiol 55:284–293. 10.1134/S0003683819030049

Binder J, Shadkchan Y, Osherov N, Krappmann S (2020) The Essential Thioredoxin Reductase of the Human Pathogenic Mold Aspergillus fumigatus Is a Promising Antifungal Target. Front Microbiol 11:. 10.3389/fmicb.2020.01383

Blin K, Shaw S, Vader L, et al (2025) antiSMASH 8.0: extended gene cluster detection capabilities and analyses of chemistry, enzymology, and regulation<? mode pagerangestyle?>. Nucleic acids research 53:W32–W38

Bonometti L, Charriat F, Hensen N, et al (2025) Genomic organization, domain assortments, and nucleotide-binding domain diversity of NLR proteins in Sordariales fungi. PLOS Genetics 21:e1011739. 10.1371/journal.pgen.1011739

Boutet E, Lieberherr D, Tognolli M, et al (2007) UniProtKB/Swiss-Prot. In: Edwards D (ed) Plant Bioinformatics. Humana Press, Totowa, NJ, pp 89–112

Buchfink B, Xie C, Huson DH (2015) Fast and sensitive protein alignment using DIAMOND. Nature Methods 12:59–60

Bucknell A, Wilson HM, Gonçalves dos Santos KC, et al (2025) Sanctuary: a Starship transposon facilitating the movement of the virulence factor ToxA in fungal wheat pathogens. mBio 16:e01371-25. 10.1128/mbio.01371-25

Bushmanova E, Antipov D, Lapidus A, Prjibelski AD (2019) rnaSPAdes: a de novo transcriptome assembler and its application to RNA-Seq data. GigaScience 8:giz100

Byers AK, Condron L, O’Callaghan M, et al (2025) Whole genome sequencing of Penicillium and Burkholderia strains antagonistic to the causal agent of kauri dieback disease (Phytophthora agathidicida) reveals biosynthetic gene clusters related to antimicrobial secondary metabolites. Molecular Ecology Resources 25:e13810. 10.1111/1755-0998.13810

Cantalapiedra CP, Hernández-Plaza A, Letunic I, et al (2021) eggNOG-mapper v2: functional annotation, orthology assignments, and domain prediction at the metagenomic scale. Molecular biology and evolution 38:5825–5829

Cantarel BL, Korf I, Robb SM, et al (2008) MAKER: an easy-to-use annotation pipeline designed for emerging model organism genomes. Genome research 18:188

Carrillo-Marín P, Tahiri G, Camuña-Pardo L, et al (2026) Phosphatidylcholine biosynthesis via ChoC is crucial for cellular integrity and virulence in Rhizopus microsporus. Virulence 17:2654119. 10.1080/21505594.2026.2654119

Carroll CS, Moore MM (2018) Ironing out siderophore biosynthesis: a review of non-ribosomal peptide synthetase (NRPS)-independent siderophore synthetases. Critical Reviews in Biochemistry and Molecular Biology 53:356–381. 10.1080/10409238.2018.1476449

Ćeranić A, Svoboda T, Berthiller F, et al (2021) Identification and Functional Characterization of the Gene Cluster Responsible for Fusaproliferin Biosynthesis in Fusarium proliferatum. Toxins 13:468. 10.3390/toxins13070468

Chand Arya G, Aditya Srivastava D, Manasherova E, et al (2022) BcHnm1, a predicted choline transporter, modulates conidial germination and virulence in *Botrytis cinerea*. Fungal Genetics and Biology 158:103653. 10.1016/j.fgb.2021.103653

Chen N (2004) Using REPEAT MASKER to Identify Repetitive Elements in Genomic Sequences. CP in Bioinformatics 5:. 10.1002/0471250953.bi0410s05

Ciuffetti LM, Manning VA, Pandelova I, et al (2010) Host-selective toxins, Ptr ToxA and Ptr ToxB, as necrotrophic effectors in the Pyrenophora tritici-repentis–wheat interaction. New Phytologist 187:911–919. 10.1111/j.1469-8137.2010.03362.x

Condon BJ, Leng Y, Wu D, et al (2013) Comparative Genome Structure, Secondary Metabolite, and Effector Coding Capacity across Cochliobolus Pathogens. PLoS Genet 9:e1003233. 10.1371/journal.pgen.1003233

Csardi G, Nepusz T (2006) The igraph software. Complex syst 1695:1–9

Dalinova A, Chisty L, Kochura D, et al (2020) Isolation and Bioactivity of Secondary Metabolites from Solid Culture of the Fungus, Alternaria sonchi. Biomolecules 10:81. 10.3390/biom10010081

Diedrich G, Spahn CMT, Stelzl U, et al (2000) Ribosomal protein L2 is involved in the association of the ribosomal subunits, tRNA binding to A and P sites and peptidyl transfer. EMBO J 19:5241–5250. 10.1093/emboj/19.19.5241

Draisma A, Loureiro C, Louwen NLL, et al (2026) BiG-SCAPE 2.0 and BiG-SLiCE 2.0: scalable, accurate and interactive sequence clustering of metabolic gene clusters. Nat Commun 17:2000. 10.1038/s41467-026-68733-5

Du H, Liang C (2019) Assembly of chromosome-scale contigs by efficiently resolving repetitive sequences with long reads. Nat Commun 10:5360. 10.1038/s41467-019-13355-3

Dunbar DA, Wormsley S, Agentis TM, Baserga SJ (1997) Mpp10p, a U3 small nucleolar ribonucleoprotein component required for pre-18S rRNA processing in yeast. Mol Cell Biol 17:5803–5812. 10.1128/MCB.17.10.5803

Edgar RC (2004) MUSCLE: multiple sequence alignment with high accuracy and high throughput. Nucleic acids research 32:1792–1797

Emanuelsson O, Brunak S, Von Heijne G, Nielsen H (2007) Locating proteins in the cell using TargetP, SignalP and related tools. Nature protocols 2:953–971

Emms DM, Kelly S (2019) OrthoFinder: phylogenetic orthology inference for comparative genomics. Genome Biol 20:238. 10.1186/s13059-019-1832-y

Farzana L, Islam SS, Rahman M, et al (2025) Addressing Bipolaris sorokiniana in wheat: challenges, management strategies, and future directions for sustainable food security. Discov Agric 3:211. 10.1007/s44279-025-00389-z

Flynn JM, Hubley R, Goubert C, et al (2020) RepeatModeler2 for automated genomic discovery of transposable element families. Proc Natl Acad Sci USA 117:9451–9457. 10.1073/pnas.1921046117

Fu T, Kim J-O, Han J-H, et al (2018) A Small GTPase RHO2 Plays an Important Role in Pre-infection Development in the Rice Blast Pathogen *Magnaporthe oryzae*. Plant Pathol J 34:470–479. 10.5423/PPJ.OA.04.2018.0069

Gilchrist CL, Chooi Y-H (2021) Clinker & clustermap. js: automatic generation of gene cluster comparison figures. Bioinformatics 37:2473–2475

Gligorijević V, Renfrew PD, Kosciolek T, et al (2021) Structure-based protein function prediction using graph convolutional networks. Nat Commun 12:3168. 10.1038/s41467-021-23303-9

Gómez-Lorenzo MG, García-Bustos JF (1998) Ribosomal P-protein Stalk Function Is Targeted by Sordarin Antifungals*. Journal of Biological Chemistry 273:25041–25044. 10.1074/jbc.273.39.25041

González K, Montanares M, Gallardo M, et al (2025) Molecular basis for the biosynthesis of the siderophore coprogen in the cheese-ripening fungus Penicillium roqueforti. Biol Res 58:51. 10.1186/s40659-025-00633-2

Gourlie R, McDonald M, Hafez M, et al (2022) The pangenome of the wheat pathogen Pyrenophora tritici-repentis reveals novel transposons associated with necrotrophic effectors ToxA and ToxB. BMC Biol 20:239. 10.1186/s12915-022-01433-w

Guerra A (2026) The pangenome: a statistical model, not a fixed biological property. Bioinformatics Advances 6:vbag069. 10.1093/bioadv/vbag069

Guimarães LC, Florczak-Wyspianska J, de Jesus LB, et al (2015) Inside the Pan-genome - Methods and Software Overview. Curr Genomics 16:245–252. 10.2174/1389202916666150423002311

Gupta PK, Vasistha NK, Aggarwal R, Joshi AK (2018) Biology of B. sorokiniana (syn. Cochliobolus sativus) in genomics era. J Plant Biochem Biotechnol 27:123–138. 10.1007/s13562-017-0426-6

Gurevich A, Saveliev V, Vyahhi N, Tesler G (2013) QUAST: quality assessment tool for genome assemblies. Bioinformatics 29:1072–1075

Hafez M, Gourlie R, McDonald M, et al (2024) Evolution of the ToxB Gene in Pyrenophora tritici-repentis and Related Species. MPMI 37:327–337. 10.1094/MPMI-08-23-0114-FI

Hage H, Rosso M-N (2021) Evolution of Fungal Carbohydrate-Active Enzyme Portfolios and Adaptation to Plant Cell-Wall Polymers. J Fungi (Basel) 7:185. 10.3390/jof7030185

Hai Y, Huang AM, Tang Y (2019) Structure-guided function discovery of an NRPS-like glycine betaine reductase for choline biosynthesis in fungi. Proceedings of the National Academy of Sciences 116:10348–10353. 10.1073/pnas.1903282116

Han J, Zhang J, Song Z, et al (2019) Genome- and MS-based mining of antibacterial chlorinated chromones and xanthones from the phytopathogenic fungus Bipolaris sorokiniana strain 11134. Appl Microbiol Biotechnol 103:5167–5181. 10.1007/s00253-019-09821-z

Hans S, Fatima Z, Hameed S (2019) Retrograde signaling disruption influences ABC superfamily transporter, ergosterol and chitin levels along with biofilm formation in *Candida albicans*. Journal de Mycologie Médicale 29:210–218. 10.1016/j.mycmed.2019.07.003

He X, Wang D, Liu J, et al (2024) Engineering the Methylerythritol Phosphate Pathway and Using a Temporal Promoter for Enhanced Lycopene Production in Rhodobacter sphaeroides HY01. J Agric Food Chem 72:28040–28047. 10.1021/acs.jafc.4c07848

Hernández-Plaza A, Szklarczyk D, Botas J, et al (2023) eggNOG 6.0: enabling comparative genomics across 12 535 organisms. Nucleic Acids Research 51:D389–D394

Hu W, Sillaots S, Lemieux S, et al (2007) Essential Gene Identification and Drug Target Prioritization in Aspergillus fumigatus. PLOS Pathogens 3:e24. 10.1371/journal.ppat.0030024

Huang J, Cook DE (2022) The contribution of DNA repair pathways to genome editing and evolution in filamentous pathogens. FEMS Microbiol Rev 46:fuac035. 10.1093/femsre/fuac035

Huang R, Zhang J, Lu L, et al (2025) High-quality genome assembly and annotation of the crested gecko (Correlophus ciliatus). G3 Genes|Genomes|Genetics 15:jkae265. 10.1093/g3journal/jkae265

Hue Y, Nam Y, Choi B, et al (2025) Comparative Genomics Reveals Conserved Ophiobolin Biosynthetic Gene Cluster and Necrotrophic Adaptation in *Bipolaris oryzae*. Plant Pathol J 41:682–698. 10.5423/PPJ.FT.08.2025.0107

Jansson H-B, Åkesson H (2003) Extracellular Matrix, Esterase and the Phytotoxin Prehelminthosporol in Infection of Barley Leaves by Bipolaris sorokiniana. European Journal of Plant Pathology 109:599–605. 10.1023/A:1024773531256

John E, Verdonk C, Singh KB, et al (2024) Regulatory insight for a Zn2Cys6 transcription factor controlling effector-mediated virulence in a fungal pathogen of wheat. PLOS Pathogens 20:e1012536. 10.1371/journal.ppat.1012536

Jones P, Binns D, Chang H-Y, et al (2014) InterProScan 5: genome-scale protein function classification. Bioinformatics 30:1236–1240

Kalyaanamoorthy S, Minh BQ, Wong TK, et al (2017) ModelFinder: fast model selection for accurate phylogenetic estimates. Nature methods 14:587–589

Kashyap PL, Kumar S, Sharma A, et al (2022) Molecular diversity, haplotype distribution and genetic variation flow of Bipolaris sorokiniana fungus causing spot blotch disease in different wheat-growing zones. J Appl Genetics 63:793–803. 10.1007/s13353-022-00716-w

Katoh K, Standley DM (2013) MAFFT multiple sequence alignment software version 7: improvements in performance and usability. Molecular biology and evolution 30:772–780

Kim M, Lee SH, Jeon J (2023) A Nucleolar Protein, MoRRP8 Is Required for Development and Pathogenicity in the Rice Blast Fungus. Mycobiology 51:273–280. 10.1080/12298093.2023.2257996

Kim NE, Dettman JR (2025) Genome mining reveals the distribution of biosynthetic gene clusters in Alternaria and related fungal taxa within the family Pleosporaceae. BMC Genomics 26:678. 10.1186/s12864-025-11754-z

Kirsch VC, Orgler C, Braig S, et al (2020) The Cytotoxic Natural Product Vioprolide A Targets Nucleolar Protein 14, Which Is Essential for Ribosome Biogenesis. Angew Chem Int Ed Engl 59:1595–1600. 10.1002/anie.201911158

Knutson BA, Hahn S (2011) Yeast Rrn7 and Human TAF1B Are TFIIB-Related RNA Polymerase I General Transcription Factors. Science 333:1637–1640. 10.1126/science.1207699

Korf I (2004) Gene finding in novel genomes. BMC Bioinformatics 5:59. 10.1186/1471-2105-5-59

Korovesi AG, Ntertilis M, Kouvelis VN (2018) Mt-*rps*3 is an ancient gene which provides insight into the evolution of fungal mitochondrial genomes. Molecular Phylogenetics and Evolution 127:74–86. 10.1016/j.ympev.2018.04.037

Kosakovsky Pond SL, Frost SDW (2005) Not So Different After All: A Comparison of Methods for Detecting Amino Acid Sites Under Selection. Mol Biol Evol 22:1208–1222. 10.1093/molbev/msi105

Kosakovsky Pond SL, Poon AF, Velazquez R, et al (2020) HyPhy 2.5—a customizable platform for evolutionary hypothesis testing using phylogenies. Molecular biology and evolution 37:295–299

Lan J, Mei S, Du Y, et al (2025) ApWD40a, a Member of the WD40-Repeat Protein Family, Is Crucial for Fungal Development, Toxin Synthesis, and Pathogenicity in the Ginseng Alternaria Leaf Blight Fungus Alternaria panax. Journal of Fungi 11:59. 10.3390/jof11010059

Langenhorst D, Fürst A-L, Alberter K, et al (2023) Soluble Enolase 1 of Candida albicans and Aspergillus fumigatus Stimulates Human and Mouse B Cells and Monocytes. J Immunol 211:804–815. 10.4049/jimmunol.2200318

Li L, Lu H, Zhang X, et al (2022) Baicalein Acts against Candida albicans by Targeting Eno1 and Inhibiting Glycolysis. Microbiology Spectrum 10:e02085–22. 10.1128/spectrum.02085-22

Li R, Yao J, Xiao J, et al (2025) Transcriptomic and functional analyses reveal a complex unexplored landscape of Botrytis cinerea colonization in rose. J Exp Bot 76:4654–4668. 10.1093/jxb/eraf219

Li R, Zheng P, Sun X, et al (2023) Genome Sequencing and Analysis Reveal Potential High-Valued Metabolites Synthesized by Lasiodiplodia iranensis DWH-2. Journal of Fungi 9:522. 10.3390/jof9050522

Liu C, Chen X, Huang Y, et al (2026) Antifungal effect of benziothiazolinone (BIT) against *Fusarium fujikuroi*: Disruption of ribosome biogenesis and inhibition of cellular proliferation. Pesticide Biochemistry and Physiology 219:107007. 10.1016/j.pestbp.2026.107007

Liu Q, Li G, Liang Y, et al (2025) Integrated genome and transcriptome analysis reveals pathogenic mechanisms of Calonectria eucalypti in Eucalyptus leaf blight. BMC Genomics 26:695. 10.1186/s12864-025-11884-4

Lu H, Xiong J, Shang Q, et al (2016) Roles of RPS41 in Biofilm Formation, Virulence, and Hydrogen Peroxide Sensitivity in Candida albicans. Curr Microbiol 72:783–787. 10.1007/s00284-016-1019-7

Maekawa S, Ueda Y, Yanagisawa S (2018) Overexpression of a Brix Domain-Containing Ribosome Biogenesis Factor ARPF2 and its Interactor ARRS1 Causes Morphological Changes and Lifespan Extension in Arabidopsis thaliana. Front Plant Sci 9:. 10.3389/fpls.2018.01177

Manning VA, Pandelova I, Dhillon B, et al (2013) Comparative Genomics of a Plant-Pathogenic Fungus, Pyrenophora tritici-repentis, Reveals Transduplication and the Impact of Repeat Elements on Pathogenicity and Population Divergence. G3 Genes|Genomes|Genetics 3:41–63. 10.1534/g3.112.004044

Marino ND, Brodsky IE (2023) Immunology: NACHT domain proteins get a prokaryotic origin story. Current Biology 33:R875–R878. 10.1016/j.cub.2023.06.084

Markham P, Robson GD, Bainbridge BW, Trinci AP (1993) Choline: its role in the growth of filamentous fungi and the regulation of mycelial morphology. FEMS Microbiol Rev 10:287–300. 10.1111/j.1574-6968.1993.tb05872.x

Masi M, Meyer S, Górecki M, et al (2018) Phytotoxic Activity of Metabolites Isolated from Rutstroemia sp.n., the Causal Agent of Bleach Blonde Syndrome on Cheatgrass (Bromus tectorum). Molecules 23:1734. 10.3390/molecules23071734

Masi M, Zonno MC, Boari A, et al (2022) Terpestacin, a toxin produced by Phoma exigua var. heteromorpha, the causal agent of a severe foliar disease of oleander (Nerium oleander L.). Natural Product Research 36:1253–1259. 10.1080/14786419.2021.1872570

McCarthy CGP, Fitzpatrick DA (2019) Pan-genome analyses of model fungal species. Microb Genom 5:e000243. 10.1099/mgen.0.000243

McDonald MC, Ahren D, Simpfendorfer S, et al (2018) The discovery of the virulence gene ToxA in the wheat and barley pathogen Bipolaris sorokiniana. Molecular Plant Pathology 19:432–439. 10.1111/mpp.12535

Mendes FK, Vanderpool D, Fulton B, Hahn MW (2020) CAFE 5 models variation in evolutionary rates among gene families. Bioinformatics 36:5516–5518

Meng S, Chao S, Xiong M, et al (2025) CaSun1, a SUN family protein, governs the pathogenicity of Colletotrichum camelliae by recruiting CaAtg8 to promote mitophagy. Hortic Res 12:uhaf121. 10.1093/hr/uhaf121

Mering C von, Huynen M, Jaeggi D, et al (2003) STRING: a database of predicted functional associations between proteins. Nucleic acids research 31:258–261

Mo C, Xie C, Wang G, et al (2021) Cyclophilin acts as a ribosome biogenesis factor by chaperoning the ribosomal protein (PlRPS15) in filamentous fungi. Nucleic Acids Res 49:12358–12376. 10.1093/nar/gkab1102

Moolhuijzen PM, See PT, Shi G, et al (2022) A global pangenome for the wheat fungal pathogen Pyrenophora tritici-repentis and prediction of effector protein structural homology. Microb Genom 8:mgen000872. 10.1099/mgen.0.000872

Murrell B, Weaver S, Smith MD, et al (2015) Gene-Wide Identification of Episodic Selection. Mol Biol Evol 32:1365–1371. 10.1093/molbev/msv035

Murrell B, Wertheim JO, Moola S, et al (2012) Detecting individual sites subject to episodic diversifying selection. PLoS Genet 8:e1002764. 10.1371/journal.pgen.1002764

Narita K, Minami A, Ozaki T, et al (2018) Total Biosynthesis of Antiangiogenic Agent (−)- Terpestacin by Artificial Reconstitution of the Biosynthetic Machinery in *Aspergillus oryzae*. J Org Chem 83:7042–7048. 10.1021/acs.joc.7b03220

O’Mara SP, Broz K, Schwister EM, et al (2023) The Fusarium graminearum Transporters Abc1 and Abc6 Are Important for Xenobiotic Resistance, Trichothecene Accumulation, and Virulence to Wheat. Phytopathology® 113:1916–1923. 10.1094/PHYTO-09-22-0345-R

Palos-Fernández R, Aguilar-Pontes MV, Puebla-Planas G, et al (2024) Copper acquisition is essential for plant colonization and virulence in a root-infecting vascular wilt fungus. PLOS Pathogens 20:e1012671. 10.1371/journal.ppat.1012671

Paradis E, Claude J, Strimmer K (2004) APE: Analyses of Phylogenetics and Evolution in R language. Bioinformatics 20:289–290. 10.1093/bioinformatics/btg412

Parker DM, Wilson AM, Nogueira C, et al (2025) Genome compartmentalization in a host-specific fungal insect pathogen reveals a putative mating type locus on an accessory chromosome. 2025.08.27.672719

Perrier M, Barber AE (2024) Unraveling the genomic diversity and virulence of human fungal pathogens through pangenomics. PLoS Pathog 20:e1012313. 10.1371/journal.ppat.1012313

Prjibelski A, Antipov D, Meleshko D, et al (2020) Using SPAdes De Novo Assembler. CP in Bioinformatics 70:e102. 10.1002/cpbi.102

Ren H, Li X, Li Y, et al (2022) Loss of function of VdDrs2, a P4-ATPase, impairs the toxin secretion and microsclerotia formation, and decreases the pathogenicity of Verticillium dahliae. Front Plant Sci 13:. 10.3389/fpls.2022.944364

Ren Z, Liu N, Jia H, et al (2024) Discovery of Aldehyde Dehydrogenase as a Potential Fungicide Target and Screening of its Natural Inhibitors against Fusarium verticillioides. J Agric Food Chem 72:19424–19435. 10.1021/acs.jafc.4c05553

Rogers LW, Koehler AM, Crouch JA, et al (2022) Comparative genomic analysis reveals contraction of gene families with putative roles in pathogenesis in the fungal boxwood pathogens Calonectria henricotiae and C. pseudonaviculata. BMC Ecol Evo 22:79. 10.1186/s12862-022-02035-4

Roy C, He X, Gahtyari NC, et al (2023) Managing spot blotch disease in wheat: Conventional to molecular aspects. Front Plant Sci 14:. 10.3389/fpls.2023.1098648

Sá-Moura B, Kornprobst M, Kharde S, et al (2017) Mpp10 represents a platform for the interaction of multiple factors within the 90S pre-ribosome. PLOS ONE 12:e0183272. 10.1371/journal.pone.0183272

Santana MF, Silva JC, Mizubuti ES, et al (2014) Characterization and potential evolutionary impact of transposable elements in the genome of Cochliobolus heterostrophus. BMC Genomics 15:536. 10.1186/1471-2164-15-536

Santos PKF, Kapheim KM (2024) Convergent Evolution Associated with the Loss of Developmental Diapause May Promote Extended Lifespan in Bees. Genome Biol Evol 16:evae255. 10.1093/gbe/evae255

Sav H, Rafati H, Öz Y, et al (2018) Biofilm Formation and Resistance to Fungicides in Clinically Relevant Members of the Fungal Genus Fusarium. Journal of Fungi 4:16. 10.3390/jof4010016

Shafaatian R, Payton MA, Reid JD (1996) PWP2, a member of the WD-repeat family of proteins, is an essential Saccharomyces cerevisiae gene involved in cell separation. Mol Gen Genet 252:101–114. 10.1007/BF02173210

Shannon P, Markiel A, Ozier O, et al (2003) Cytoscape: a software environment for integrated models of biomolecular interaction networks. Genome research 13:2498

Sharma P, Mishra S, Singroha G, et al (2022) Phylogeographic Diversity Analysis of Bipolaris sorokiniana (Sacc.) Shoemaker Causing Spot Blotch Disease in Wheat and Barley. Genes 13:2206. 10.3390/genes13122206

Shi H-B, Chen N, Zhu X-M, et al (2019) F-box proteins MoFwd1, MoCdc4 and MoFbx15 regulate development and pathogenicity in the rice blast fungus Magnaporthe oryzae. Environmental Microbiology 21:3027–3045. 10.1111/1462-2920.14699

Shor E, Garcia-Rubio R, DeGregorio L, Perlin DS (2020) A Noncanonical DNA Damage Checkpoint Response in a Major Fungal Pathogen. mBio 11:10.1128/mbio.03044-20. 10.1128/mbio.03044-20

Shukla AK, Sahoo R, Kadoo N (2025) Comprehensive Genomic Analysis of Bipolaris sorokiniana Strains: Insights Into Genetic Diversity and Pathogenicity. Plant Pathology 74:2054–2073. 10.1111/ppa.70008

Simão FA, Waterhouse RM, Ioannidis P, et al (2015) BUSCO: assessing genome assembly and annotation completeness with single-copy orthologs. Bioinformatics 31:3210–3212

Simms CL, Kim KQ, Yan LL, et al (2018) Interactions between the mRNA and Rps3/uS3 at the entry tunnel of the ribosomal small subunit are important for no-go decay. PLOS Genetics 14:e1007818. 10.1371/journal.pgen.1007818

Singh Y, Nair AM, Verma PK (2021) Surviving the odds: From perception to survival of fungal phytopathogens under host-generated oxidative burst. Plant Communications 2:100142. 10.1016/j.xplc.2021.100142

Smith MD, Wertheim JO, Weaver S, et al (2015) Less Is More: An Adaptive Branch-Site Random Effects Model for Efficient Detection of Episodic Diversifying Selection. Mol Biol Evol 32:1342–1353. 10.1093/molbev/msv022

Somani D, Adhav R, Prashant R, Kadoo NY (2019) Transcriptomics analysis of propiconazole-treated Cochliobolus sativus reveals new putative azole targets in the plant pathogen. Funct Integr Genomics 19:453–465. 10.1007/s10142-019-00660-9

Son H, Seo Y-S, Min K, et al (2011) A Phenome-Based Functional Analysis of Transcription Factors in the Cereal Head Blight Fungus, Fusarium graminearum. PLOS Pathogens 7:e1002310. 10.1371/journal.ppat.1002310

Sperschneider J, Dodds PN (2022) EffectorP 3.0: Prediction of Apoplastic and Cytoplasmic Effectors in Fungi and Oomycetes. MPMI 35:146–156. 10.1094/MPMI-08-21-0201-R

Stanke M, Morgenstern B (2005) AUGUSTUS: a web server for gene prediction in eukaryotes that allows user-defined constraints. Nucleic acids research 33:W465–W467

Sun X, Li Y, Xu H, et al (2025) Terpestacin and Its Derivatives: Bioactivities and Syntheses. Chemistry & Biodiversity 22:e202401905. 10.1002/cbdv.202401905

Suyama M, Torrents D, Bork P (2006) PAL2NAL: robust conversion of protein sequence alignments into the corresponding codon alignments. Nucleic acids research 34:W609– W612

Sweany RR, Breunig M, Opoku J, et al (2022) Why Do Plant-Pathogenic Fungi Produce Mycotoxins? Potential Roles for Mycotoxins in the Plant Ecosystem. Phytopathology® 112:2044–2051. 10.1094/PHYTO-02-22-0053-SYM

Tang L, Sahasranaman A, Jakovljevic J, et al (2008) Interactions among Ytm1, Erb1, and Nop7 Required for Assembly of the Nop7-Subcomplex in Yeast Preribosomes. Molecular Biology of the Cell 19:2844–2856. 10.1091/mbc.e07-12-1281

Tang L, Zhai H, Zhang S, et al (2023) Functional Characterization of Aldehyde Dehydrogenase in Fusarium graminearum. Microorganisms 11:2875. 10.3390/microorganisms11122875

Teufel F, Almagro Armenteros JJ, Johansen AR, et al (2022) SignalP 6.0 predicts all five types of signal peptides using protein language models. Nature biotechnology 40:1023–1025

Upton GJG (1992) Fisher’s Exact Test. Journal of the Royal Statistical Society Series A (Statistics in Society) 155:395. 10.2307/2982890

Urban M, Cuzick A, Seager J, et al (2020) PHI-base: the pathogen–host interactions database. Nucleic acids research 48:D613–D620

Vanden Broeck A, Klinge S (2022) An emerging mechanism for the maturation of the Small Subunit Processome. Current Opinion in Structural Biology 73:102331. 10.1016/j.sbi.2022.102331

Viani A, Sinha P, Sharma T, Bhar LM (2017) A model for forecasting spot blotch disease in wheat. Australasian Plant Pathol 46:601–609. 10.1007/s13313-017-0514-z

Walton JD (1994) Deconstructing the Cell Wall. Plant Physiol 104:1113–1118. 10.1104/pp.104.4.1113

Wang B, Liang X, Gleason ML, et al (2020) A chromosome-scale assembly of the smallest Dothideomycete genome reveals a unique genome compaction mechanism in filamentous fungi. BMC Genomics 21:321. 10.1186/s12864-020-6732-8

Wang M, Yang X, Ruan R, et al (2018) Csn5 Is Required for the Conidiogenesis and Pathogenesis of the Alternaria alternata Tangerine Pathotype. Front Microbiol 9:. 10.3389/fmicb.2018.00508

Wheeler TJ, Clements J, Eddy SR, et al (2012) Dfam: a database of repetitive DNA based on profile hidden Markov models. Nucleic acids research 41:D70–D82

Wong TK, Ly-Trong N, Ren H, et al (2026) IQ-TREE 3: phylogenomic inference software using complex evolutionary models. Molecular Biology and Evolution 43:msag117

Wu C, Guo Z, Zhang M, et al (2022) Golgi-localized calcium/manganese transporters FgGdt1 and FgPmr1 regulate fungal development and virulence by maintaining Ca2+ and Mn2+ homeostasis in Fusarium graminearum. Environmental Microbiology 24:4623–4640. 10.1111/1462-2920.16128

Wyatt NA, Richards JK, Brueggeman RS, Friesen TL (2020) A Comparative Genomic Analysis of the Barley Pathogen Pyrenophora teres f. teres Identifies Subtelomeric Regions as Drivers of Virulence. MPMI 33:173–188. 10.1094/MPMI-05-19-0128-R

Xu D, Xue M, Shen Z, et al (2021) Phytotoxic Secondary Metabolites from Fungi. Toxins 13:261. 10.3390/toxins13040261

Xu Y, Liu Y, Wang Y, et al (2024) Whole-Genome Sequencing and Genome Annotation of Pathogenic Elsinoë batatas Causing Stem and Foliage Scab Disease in Sweet Potato. Journal of Fungi 10:882. 10.3390/jof10120882

Xu Z, Tong Q, Lv W, et al (2023) Phosphocholine cytidylyltransferase MoPct1 is crucial for vegetative growth, conidiation, and appressorium-mediated plant infection by Magnaporthe oryzae. Front Microbiol 14:. 10.3389/fmicb.2023.1136168

Yadav S, Raazi Z, Shivaraj SM, et al (2022) Whole Genome Sequencing and Comparative Genomics of Indian Isolates of Wheat Spot Blotch Pathogen Bipolaris sorokiniana Reveals Expansion of Pathogenicity Gene Clusters. Pathogens 12:1. 10.3390/pathogens12010001

Yan X, Kuang B-H, Ma S, et al (2024) NOP14-mediated ribosome biogenesis is required for mTORC2 activation and predicts rapamycin sensitivity. J Biol Chem 300:105681. 10.1016/j.jbc.2024.105681

Yang C, Tang L, Qin L, et al (2023) mRNA Turnover Protein 4 Is Vital for Fungal Pathogenicity and Response to Oxidative Stress in Sclerotinia sclerotiorum. Pathogens 12:281. 10.3390/pathogens12020281

Ye W, Liu T, Zhang W, et al (2019) Disclosure of the Molecular Mechanism of Wheat Leaf Spot Disease Caused by Bipolaris sorokiniana through Comparative Transcriptome and Metabolomics Analysis. International Journal of Molecular Sciences 20:6090. 10.3390/ijms20236090

Yu F-Y, Chiu C-M, Lee Y-Z, et al (2020) Polyketide Synthase Gene Expression in Relation to Chloromonilicin and Melanin Production in Monilinia fructicola. Phytopathology® 110:1465–1475. 10.1094/PHYTO-02-20-0059-R

Yuan Y, Mao X, Abubakar YS, et al (2023) Genome-Wide Characterization of the RNA Exosome Complex in Relation to Growth, Development, and Pathogenicity of Fusarium graminearum. Microbiology Spectrum 11:e05058–22. 10.1128/spectrum.05058-22

Yuste-Lisbona FJ, Fernández-Lozano A, Pineda B, et al (2020) ENO regulates tomato fruit size through the floral meristem development network. Proceedings of the National Academy of Sciences 117:8187–8195. 10.1073/pnas.1913688117

Zdouc MM, Blin K, Louwen NLL, et al (2025) MIBiG 4.0: advancing biosynthetic gene cluster curation through global collaboration. Nucleic Acids Res 53:D678–D690. 10.1093/nar/gkae1115

Zhang H, Zhao H, Huang Y, Zou Y (2024) Genome Mining Reveals the Biosynthesis of Sativene and Its Oxidative Conversion to seco-Sativene. Org Lett 26:338–343. 10.1021/acs.orglett.3c04005

Zhang L, Zhang B, Shao L, et al (2025) Genome Analysis of Alternaria alstroemeriae L6 Associated with Black Spot of Strawberry: Secondary Metabolite Biosynthesis and Virulence. Journal of Fungi 11:710. 10.3390/jof11100710

Zhang W, Yang Q, Yang L, et al (2023) High-Quality Nuclear Genome and Mitogenome of Bipolaris sorokiniana LK93, a Devastating Pathogen Causing Wheat Root Rot. MPMI 36:452–456. 10.1094/MPMI-09-22-0196-A

Zhang W, Zhang X, Li K, et al (2018) Introgression and gene family contraction drive the evolution of lifestyle and host shifts of hypocrealean fungi. Mycology 9:176–188. 10.1080/21501203.2018.1478333

Zhao J, Yang Y, Fan Y, et al (2020) Ribosomal Protein L40e Fused With a Ubiquitin Moiety Is Essential for the Vegetative Growth, Morphological Homeostasis, Cell Cycle Progression, and Pathogenicity of Cryptococcus neoformans. Front Microbiol 11:. 10.3389/fmicb.2020.570269

Zheng J, Ge Q, Yan Y, et al (2023) dbCAN3: automated carbohydrate-active enzyme and substrate annotation. Nucleic acids research 51:W115–W121

Zhou S-W, Zhu Y, Qin X-J, et al (2025) D-limonene inhibits the growth of Fusarium proliferatum by decreasing H3K9ac and H3K27ac modifications. BMC Genomics 27:55. 10.1186/s12864-025-12389-w

