## Supplementary material for "Integrated Pangenomic and Systems Biology Analyses Reveal the Genomic Basis of Virulence and Adaptation in *Bipolaris sorokiniana*": Figures S1-S17

**Supplementary Figures**

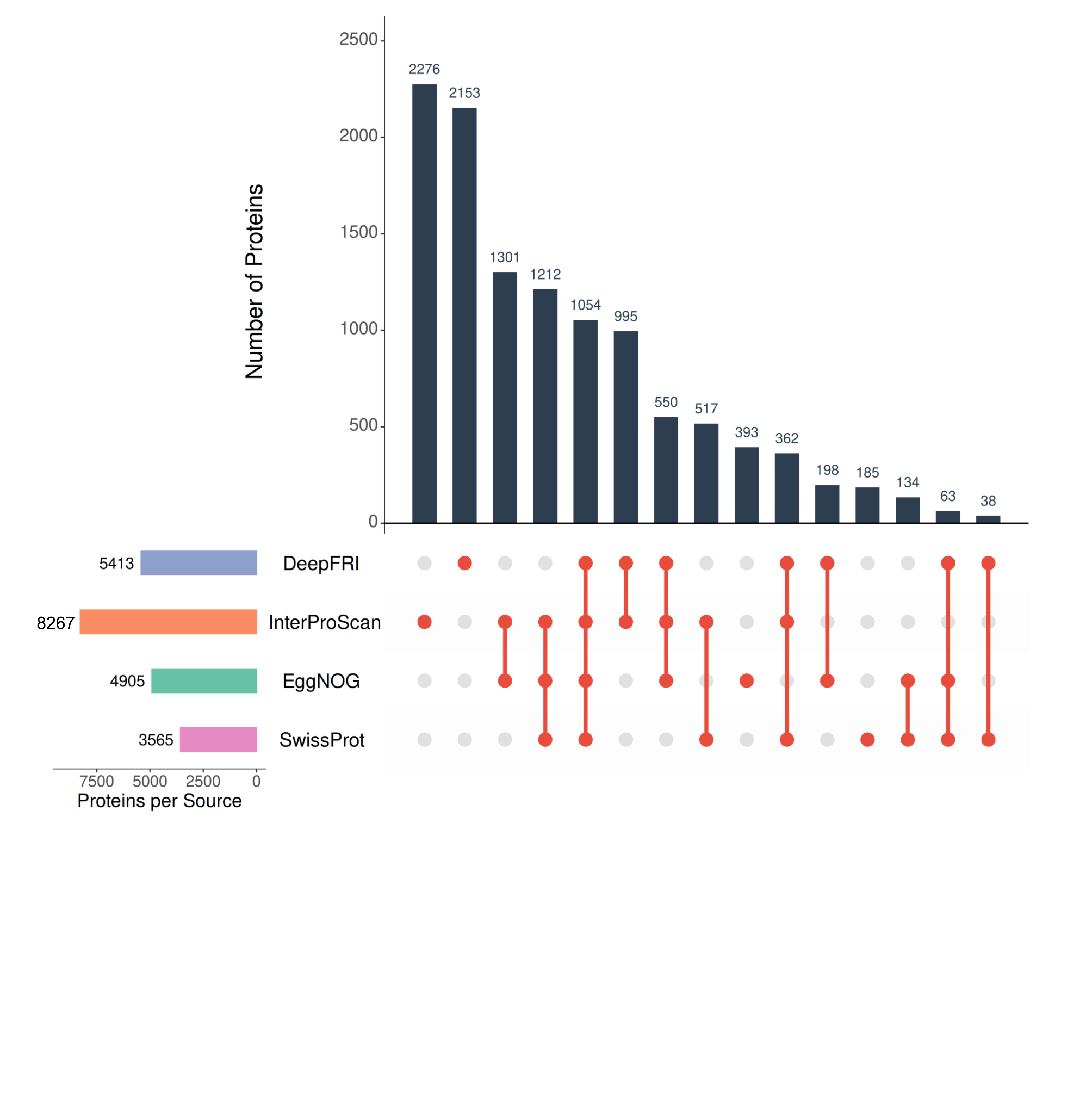

**Figure S1:** Overlap of functional annotation across multiple annotation sources. UpSet plot showing the overlap of Gene Ontology (GO) annotations for predicted orthogroups across four annotation sources: InterProScan, EggNOG, SwissProt, and DeepFRI. Each set represents proteins annotated by a given source, while the intersection bars indicate the number of proteins sharing annotations between one or more sources. The matrix layout below the bars specifies the combination of sources contributing to each intersection. The left-hand bar plot shows the total number of proteins annotated by each source.

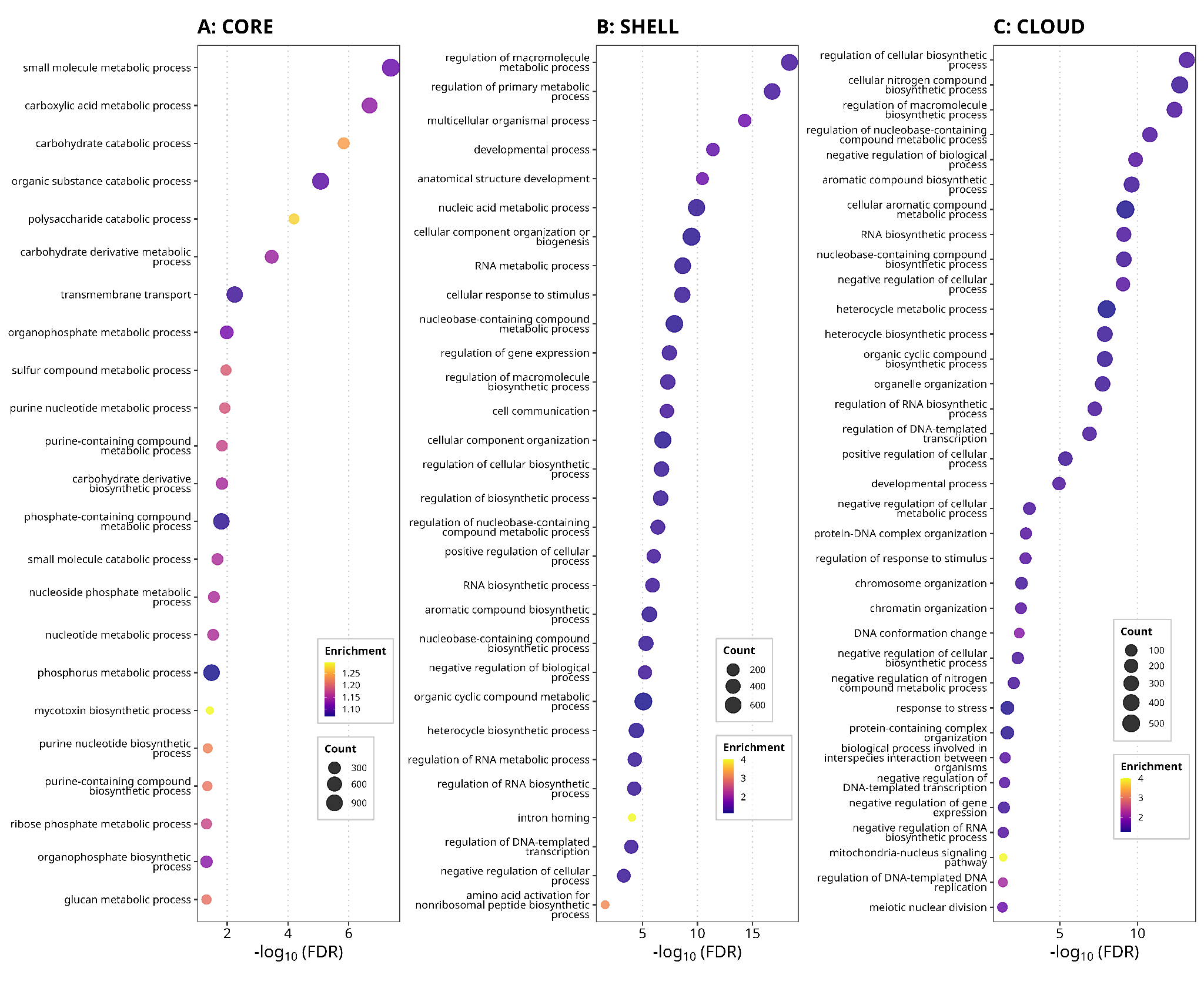

**Figure S2:** Gene Ontology (GO) enrichment analysis of pangenome orthogroup categories (Core, Shell, and Cloud) in *Bipolaris sorokiniana*.

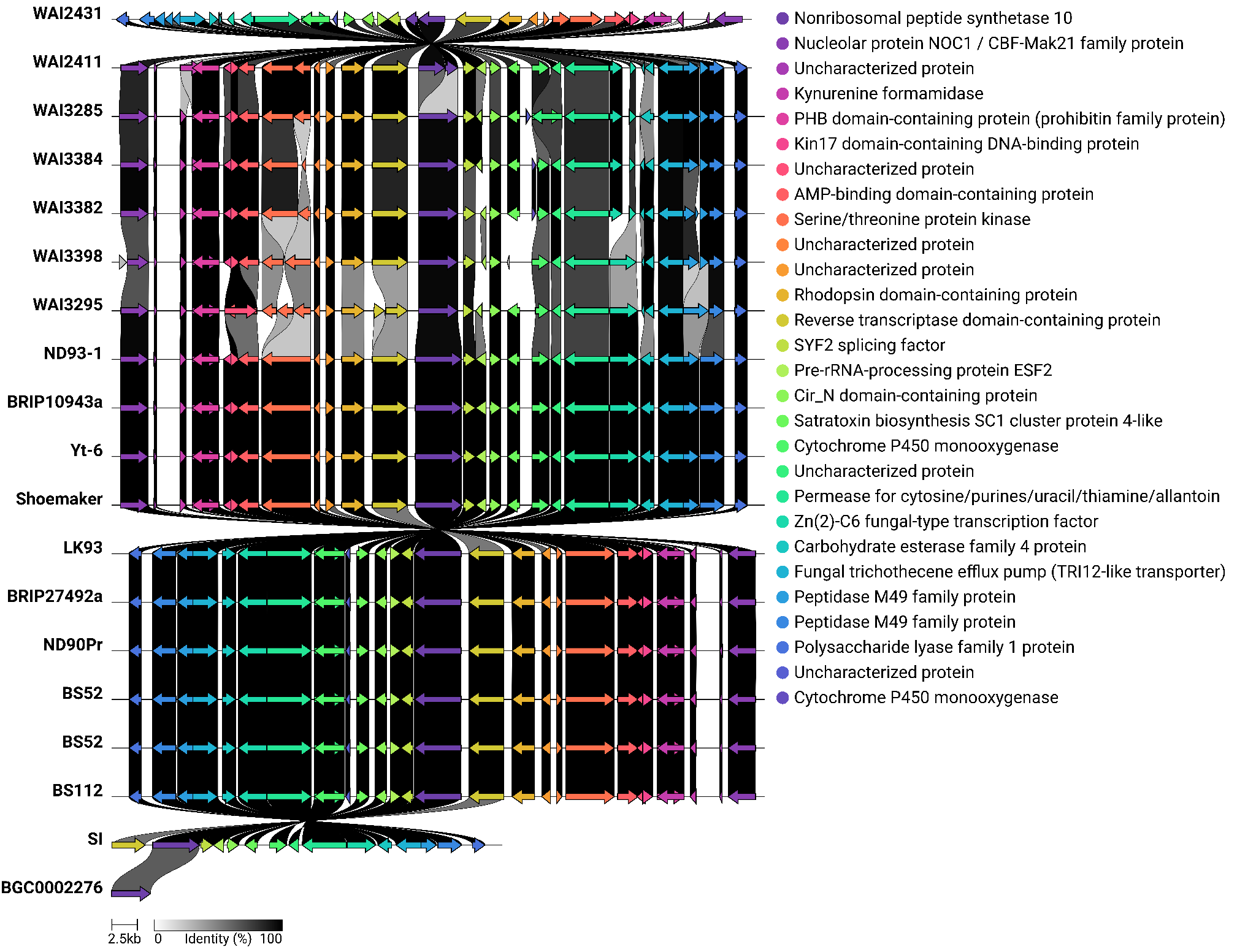

**Figure S3:** Comparative clinker visualization of the choline biosynthetic gene cluster (BGC0002276) identified across *B. sorokiniana* genomes. Conserved gene organization and synteny with the corresponding MIBiG reference cluster illustrate the high conservation of this NRPS-like biosynthetic locus, which was detected in 18 strains.

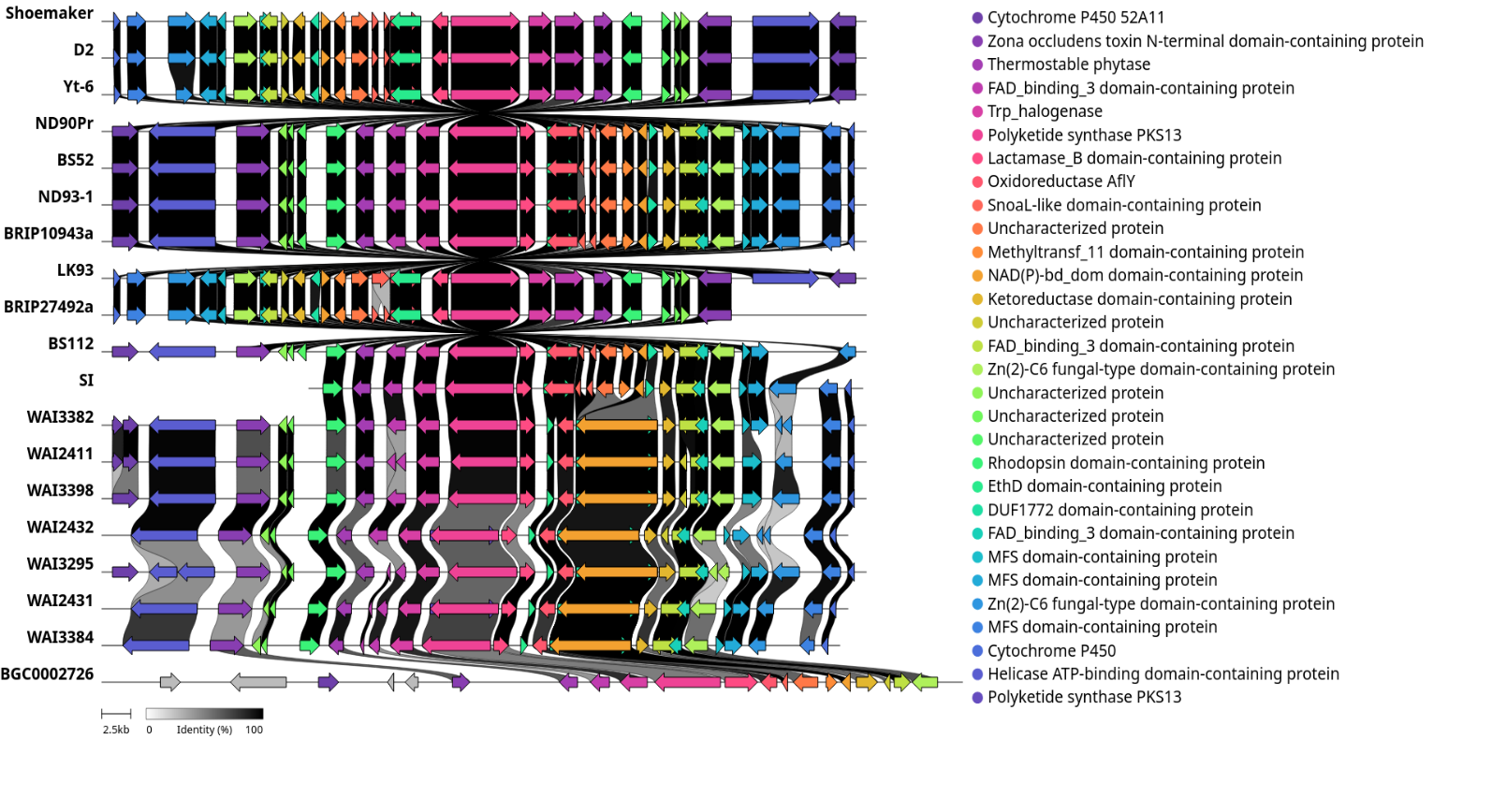

**Figure S4:** Comparative clinker visualization of the 4-chloropinselin biosynthetic gene cluster (BGC0002726) across *B. sorokiniana* genomes. The conserved arrangement of biosynthetic genes shows strong synteny with the MIBiG reference cluster, supporting the widespread conservation of this T1PKS-associated secondary metabolite pathway across 18 strains.

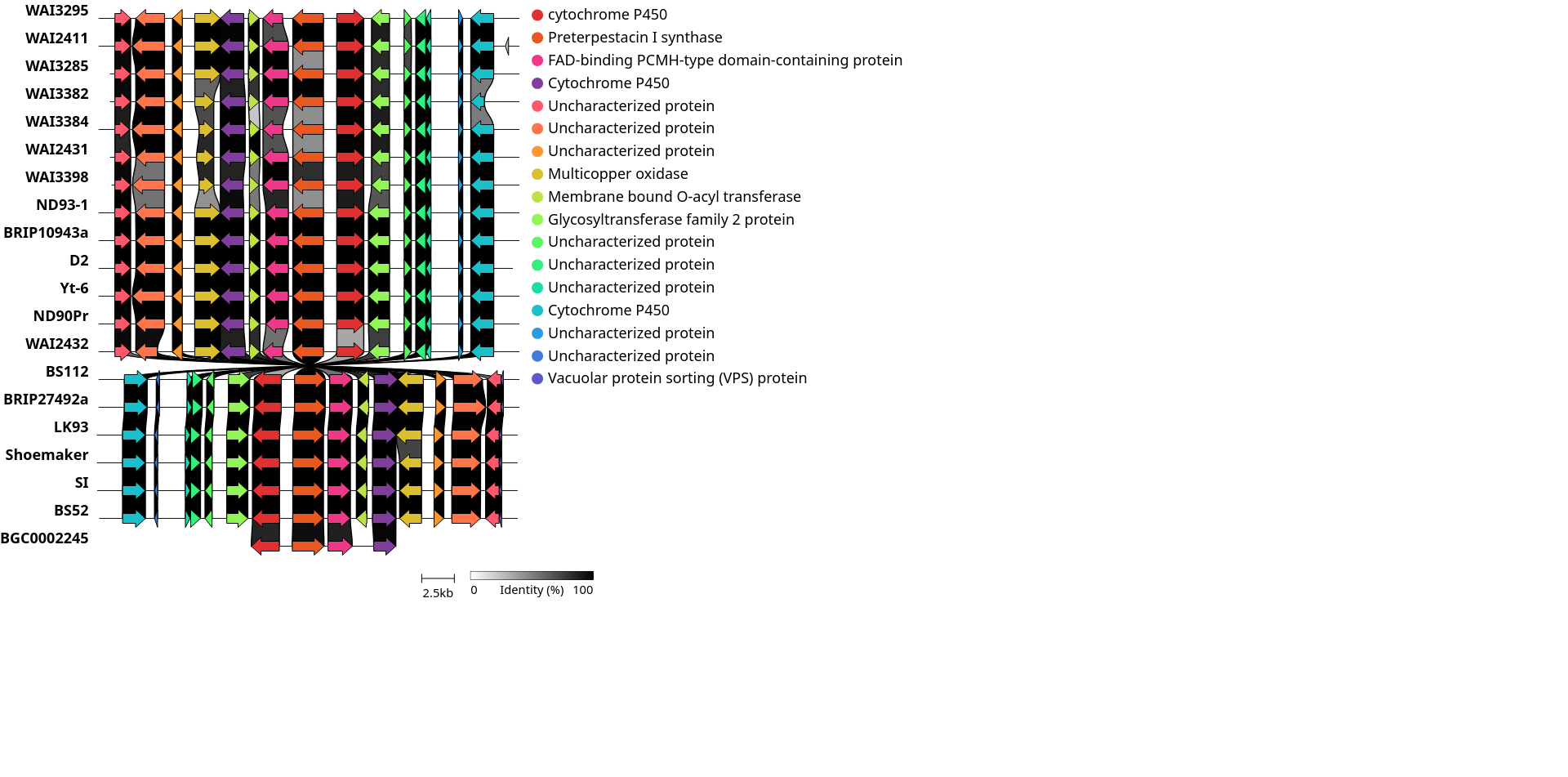

**Figure S5:** Comparative clinker visualization of the terpestacin biosynthetic gene cluster (BGC0002245) identified in *B. sorokiniana*. Conserved gene content and organization relative to the MIBiG reference cluster indicate that this terpene biosynthetic pathway is completely conserved across all 19 analyzed genomes.

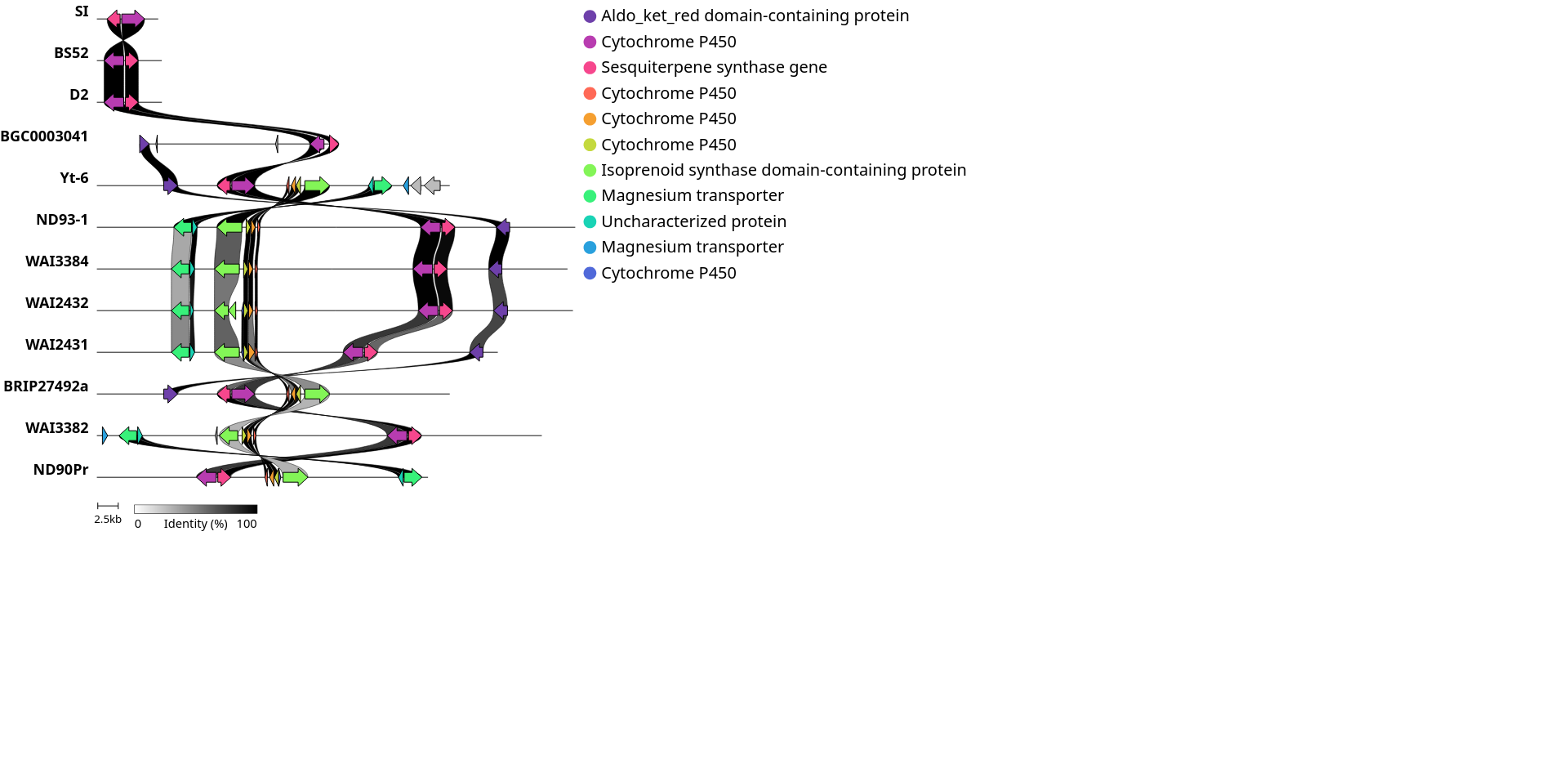

**Figure S6:** Comparative clinker visualization of the prehelminthosporol/sativene biosynthetic gene cluster (BGC0003041) across *B. sorokiniana* genomes. Conserved synteny with the reference cluster demonstrates the presence of this terpene-associated biosynthetic pathway in 11 strains, highlighting lineage-specific conservation of genes implicated in sesquiterpene biosynthesis and pathogenic adaptation.

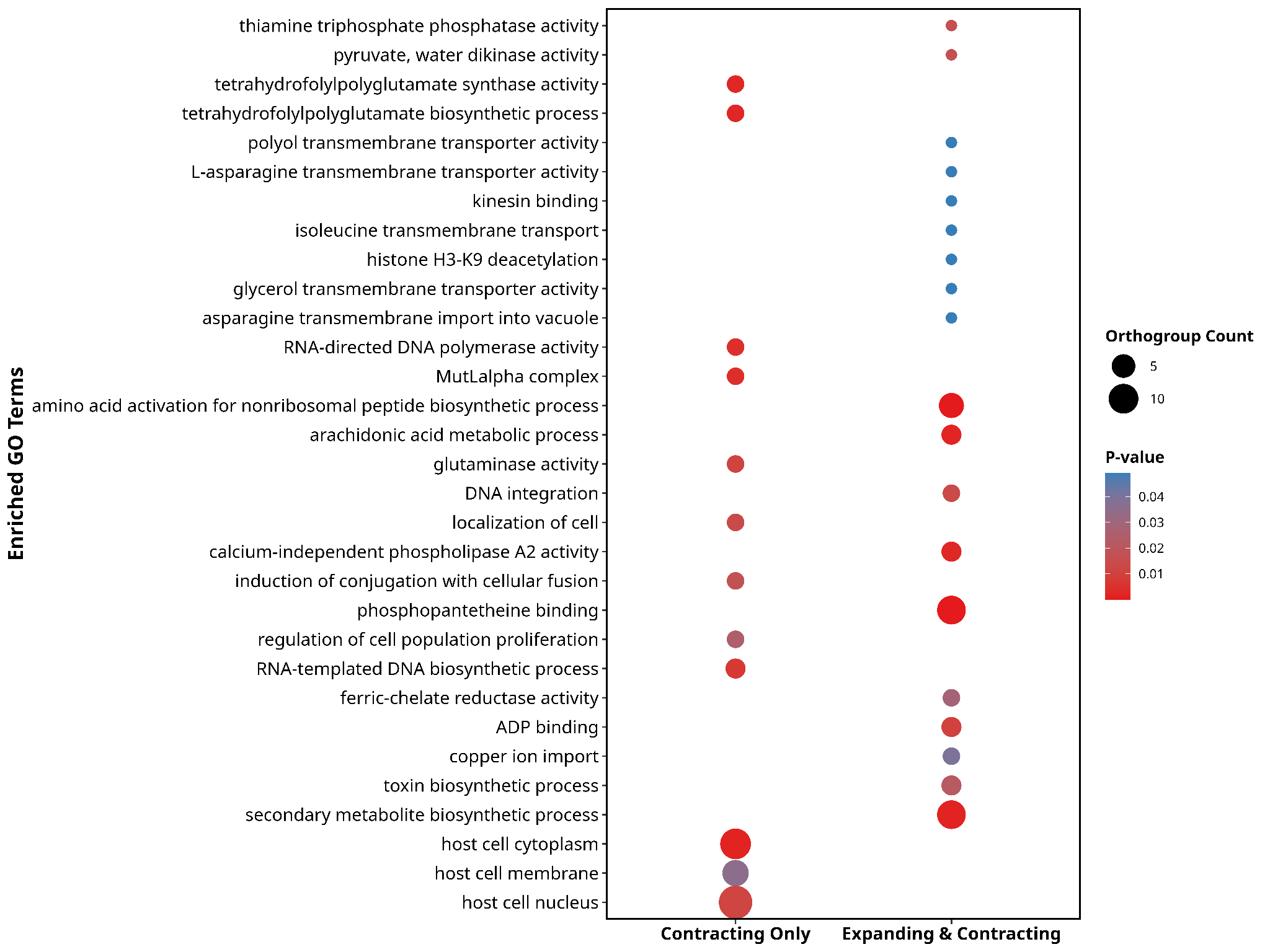

**Figure S7:** Gene Ontology (GO) enrichment analysis of significantly expanded and contracted orthogroups identified by CAFE.

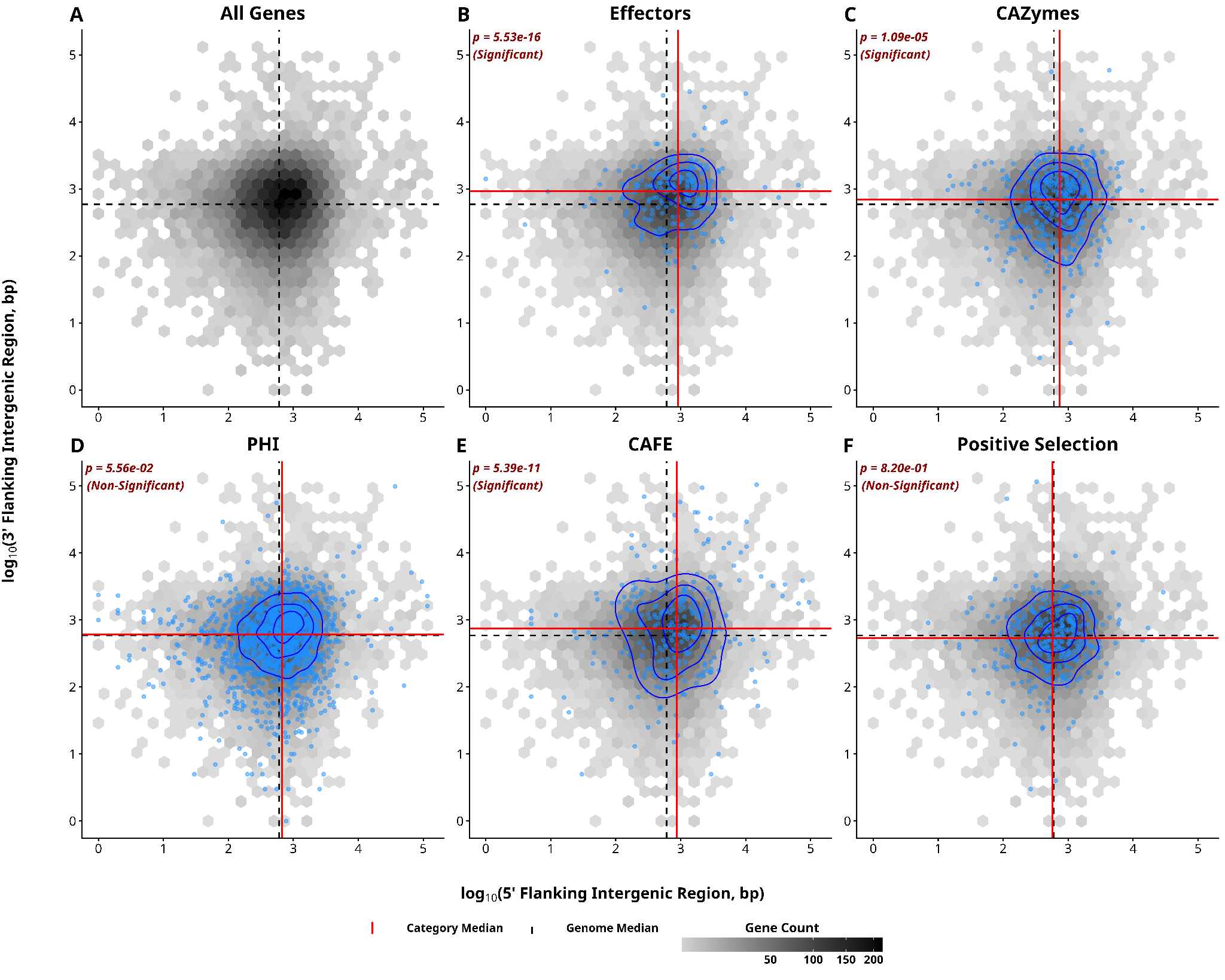

**Figure S8:** Genome compartmentalization analysis of the chromosome-level *B. sorokiniana* LK93 genome. Distribution of genes based on 5′ and 3′ flanking intergenic regions (FIRs) demonstrates a one-compartment genome architecture, lacking distinct gene-dense and gene-sparse compartments.

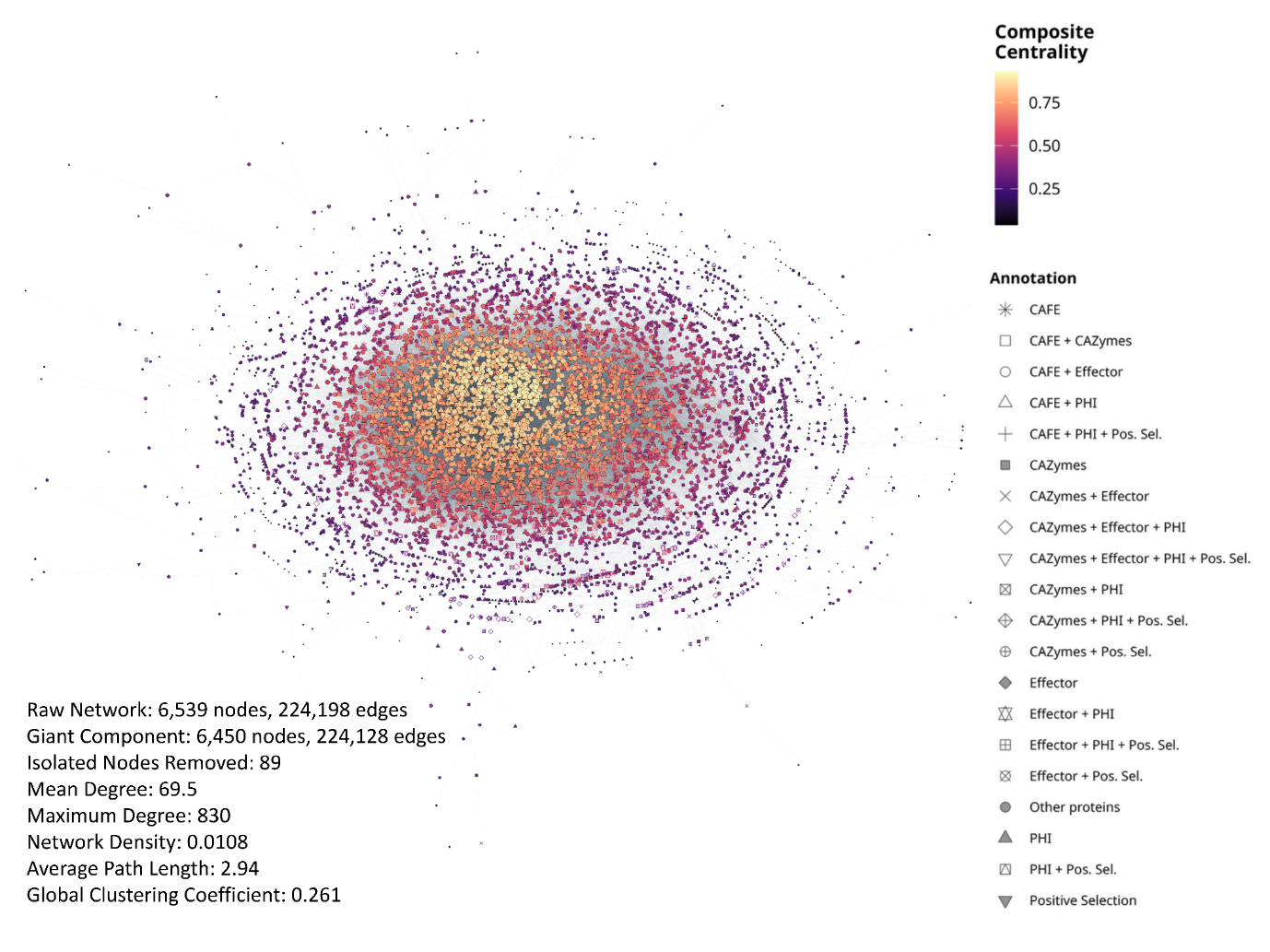

**Figure S9:** Global STRING-derived protein–protein interaction (PPI) network of the core proteome in *B. sorokiniana*. The figure represents the large-scale protein–protein interaction (PPI) network constructed from conserved core orthogroups identified across multiple *B. sorokiniana* genomes using the STRING database. The complete raw network consisted of 6,539 nodes and 224,198 edges, representing predicted and experimentally supported functional protein associations. Extraction of the giant connected component retained 6,450 nodes and 224,128 edges, indicating extensive connectivity among conserved proteins within the fungal core proteome. Nodes represent core orthogroups/proteins, while edges denote functional interactions inferred from STRING evidence channels.

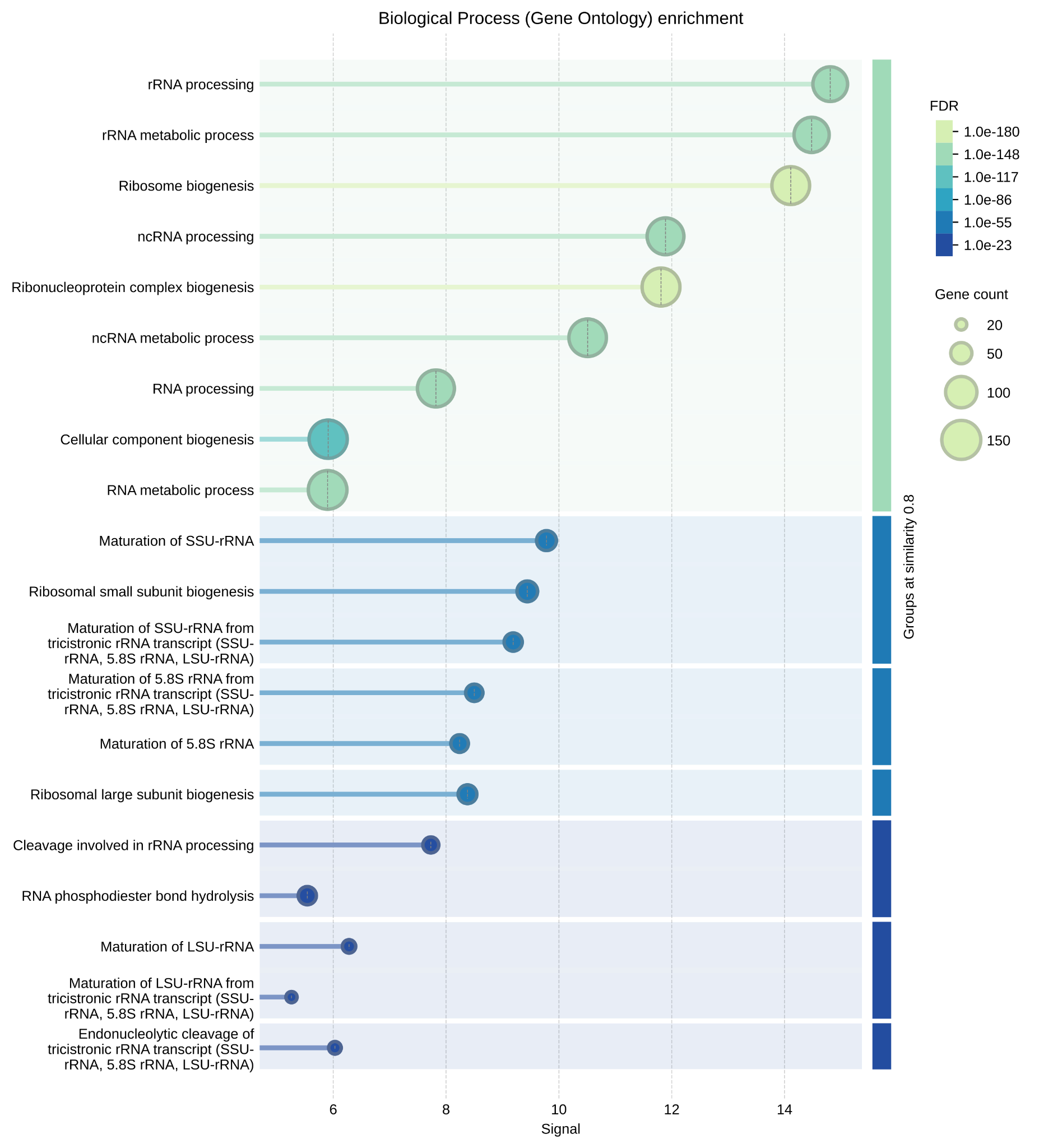

**Figure S10:** Gene Ontology (GO) enrichment analysis of Biological Processes for Community 2 (C2) identified from the STRING-derived protein–protein interaction (PPI) network of the top 10% core proteome in *B. sorokiniana*.

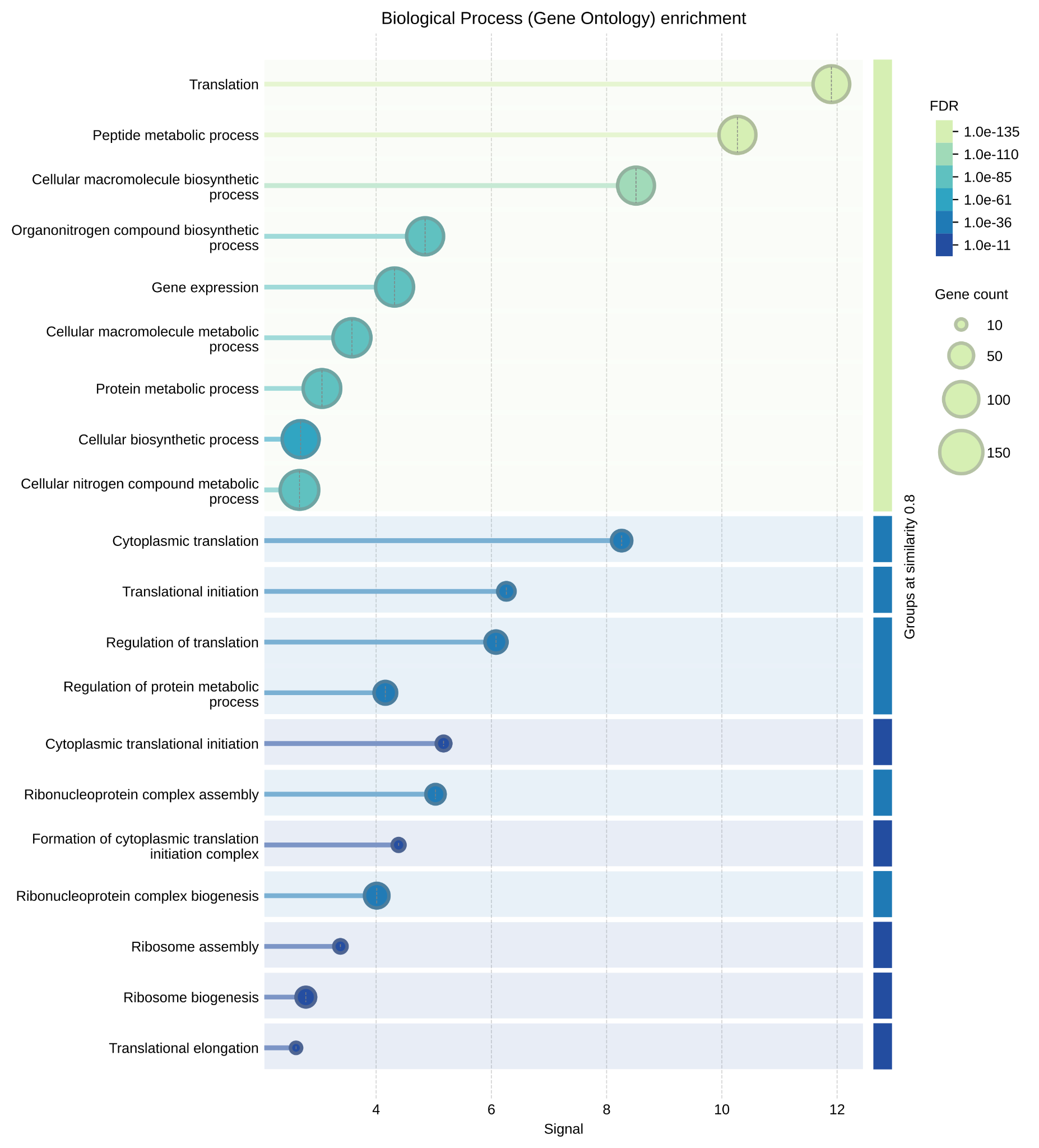

**Figure S11:** Gene Ontology (GO) enrichment analysis of Biological Processes for Community 4 (C4) identified from the STRING-derived protein–protein interaction (PPI) network of the top 10% core proteome in *B. sorokiniana*.

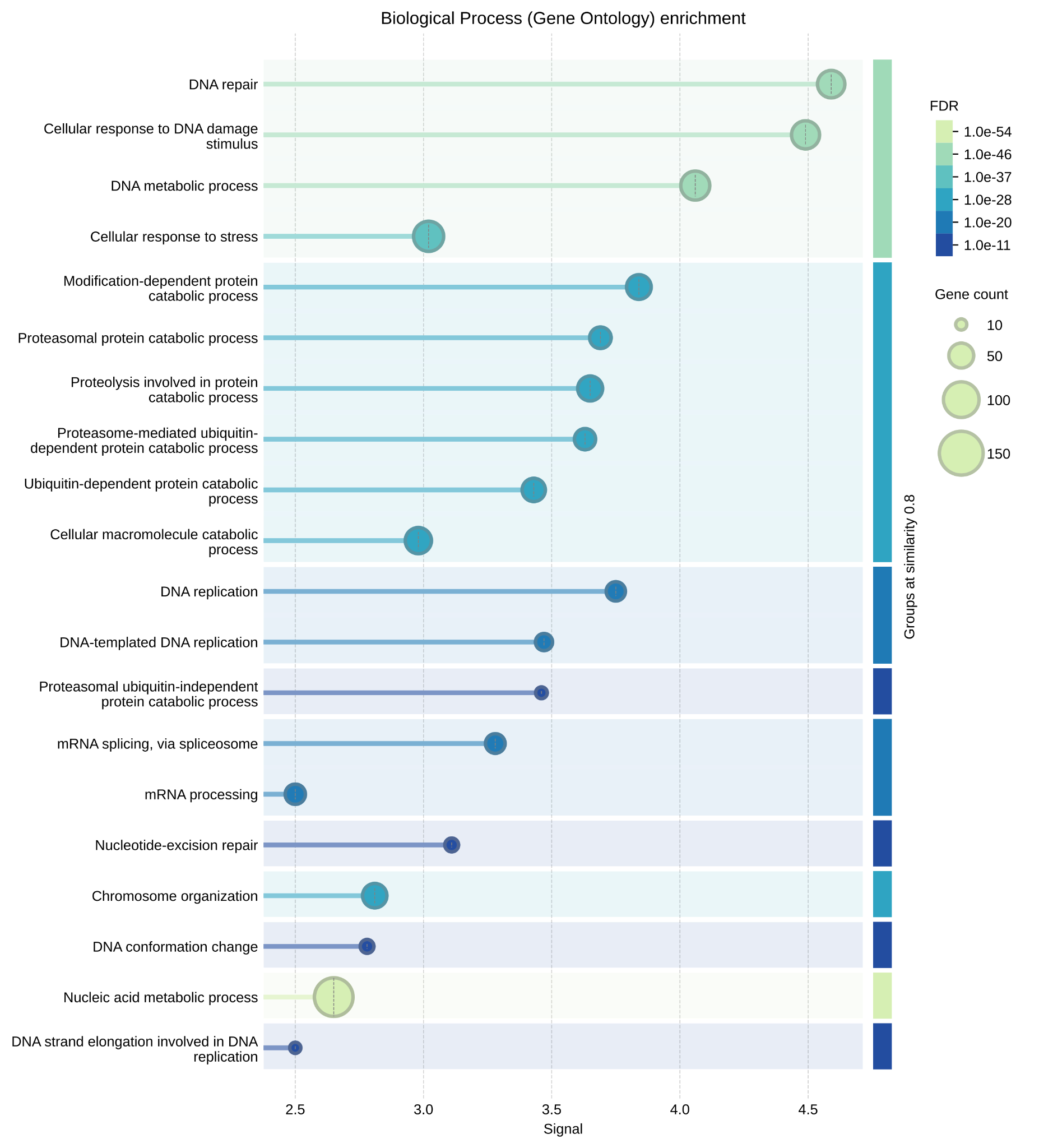

**Figure S12:** Gene Ontology (GO) enrichment analysis of Biological Processes for Community 1 (C1) identified from the STRING-derived protein–protein interaction (PPI) network of the top 10% core proteome in *B. sorokiniana*.

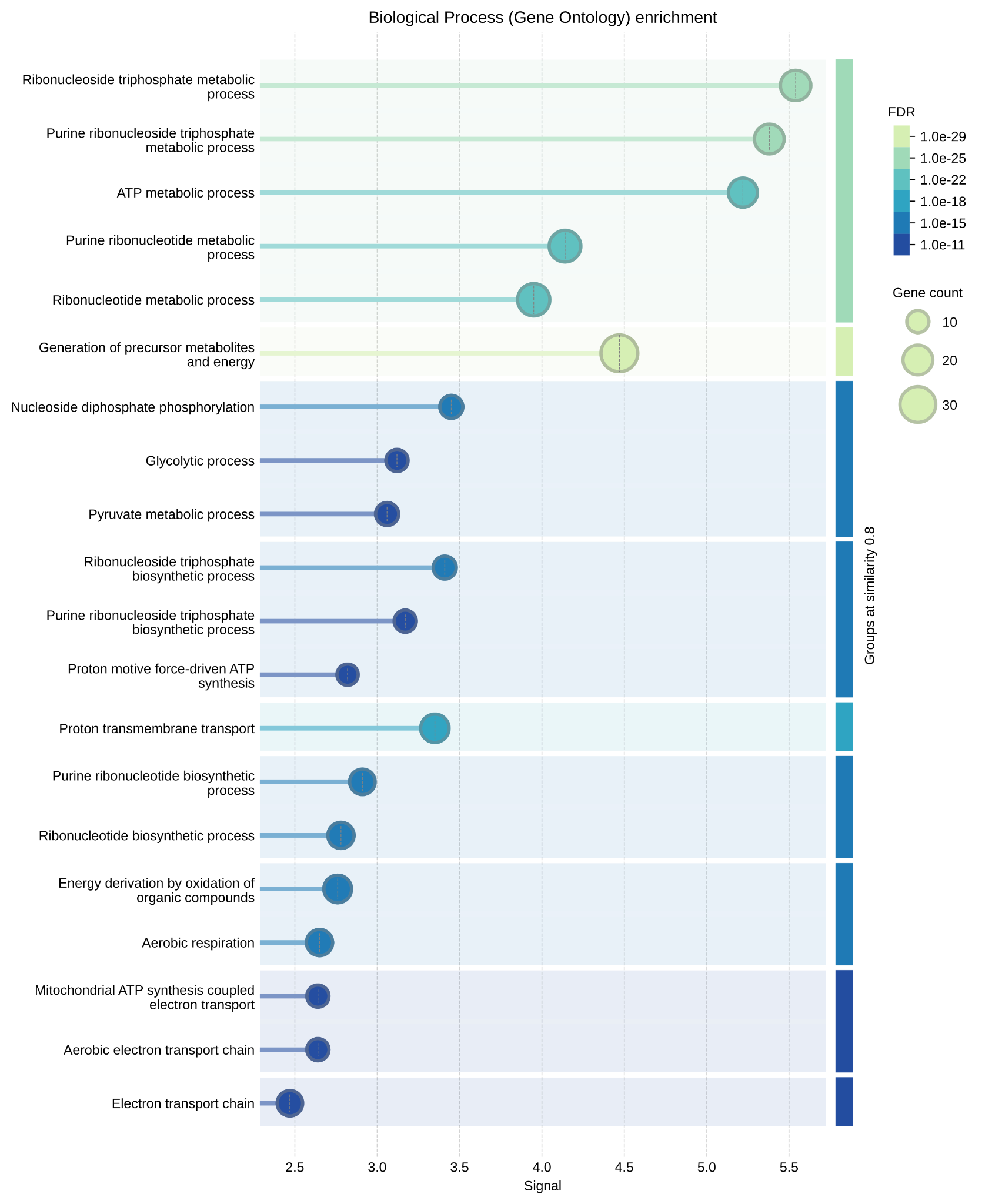

**Figure S13:** Gene Ontology (GO) enrichment analysis of Biological Processes for Community 3 (C3) identified from the STRING-derived protein–protein interaction (PPI) network of the top 10% core proteome in *B. sorokiniana*.

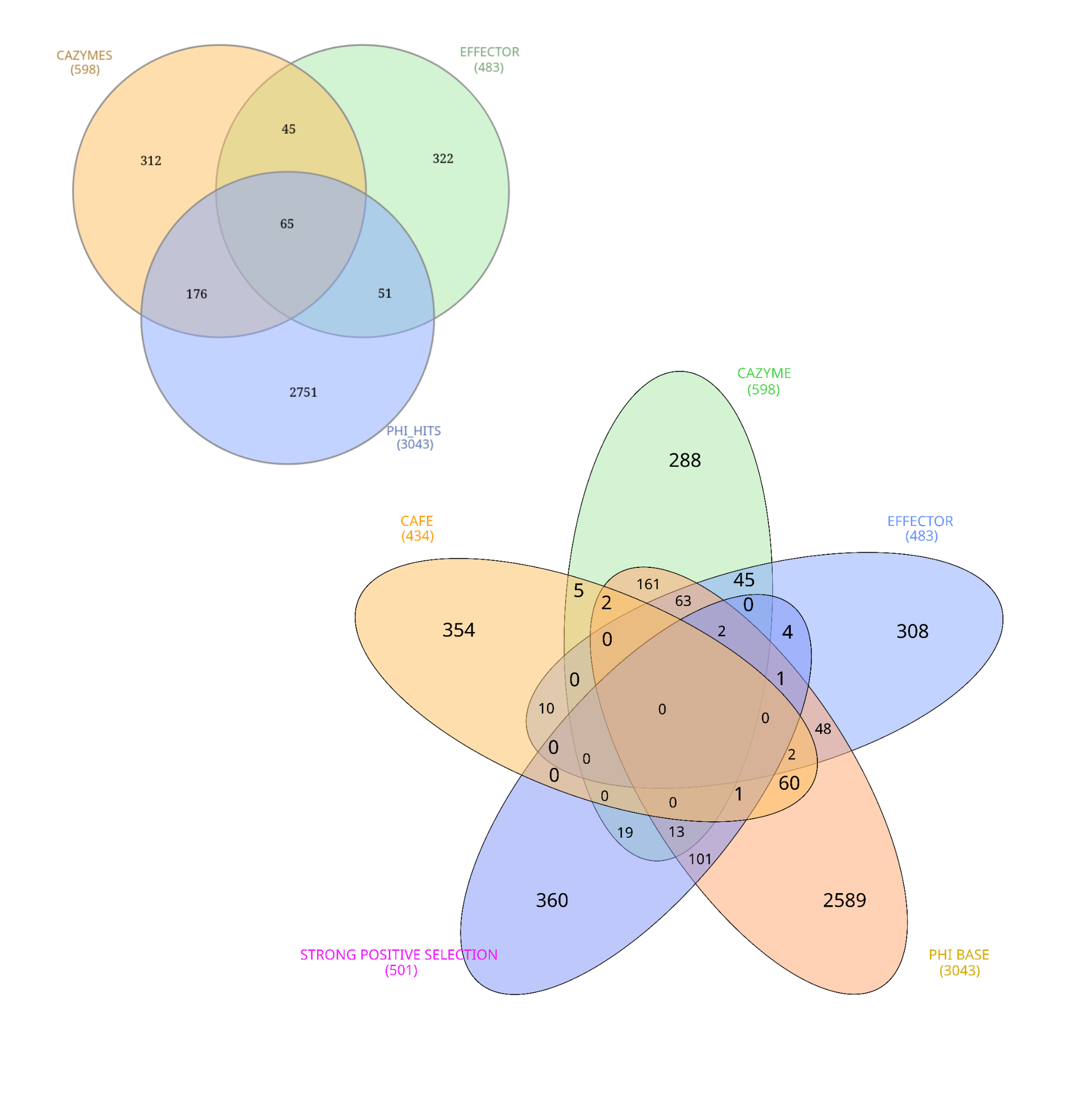

**Figure S14:** Venn diagram illustrating the overlap among orthogroups associated with carbohydrate-active enzymes (CAZymes), pathogen–host interaction (PHI-base) homologs, predicted effector proteins, genes under positive selection, and gene families identified as significantly expanded or contracted by CAFE analysis. The diagram highlights shared and unique orthogroups across these functional and evolutionary categories, providing an overview of candidate genes potentially involved in pathogenicity, host adaptation, and genome evolution in the pangenome.

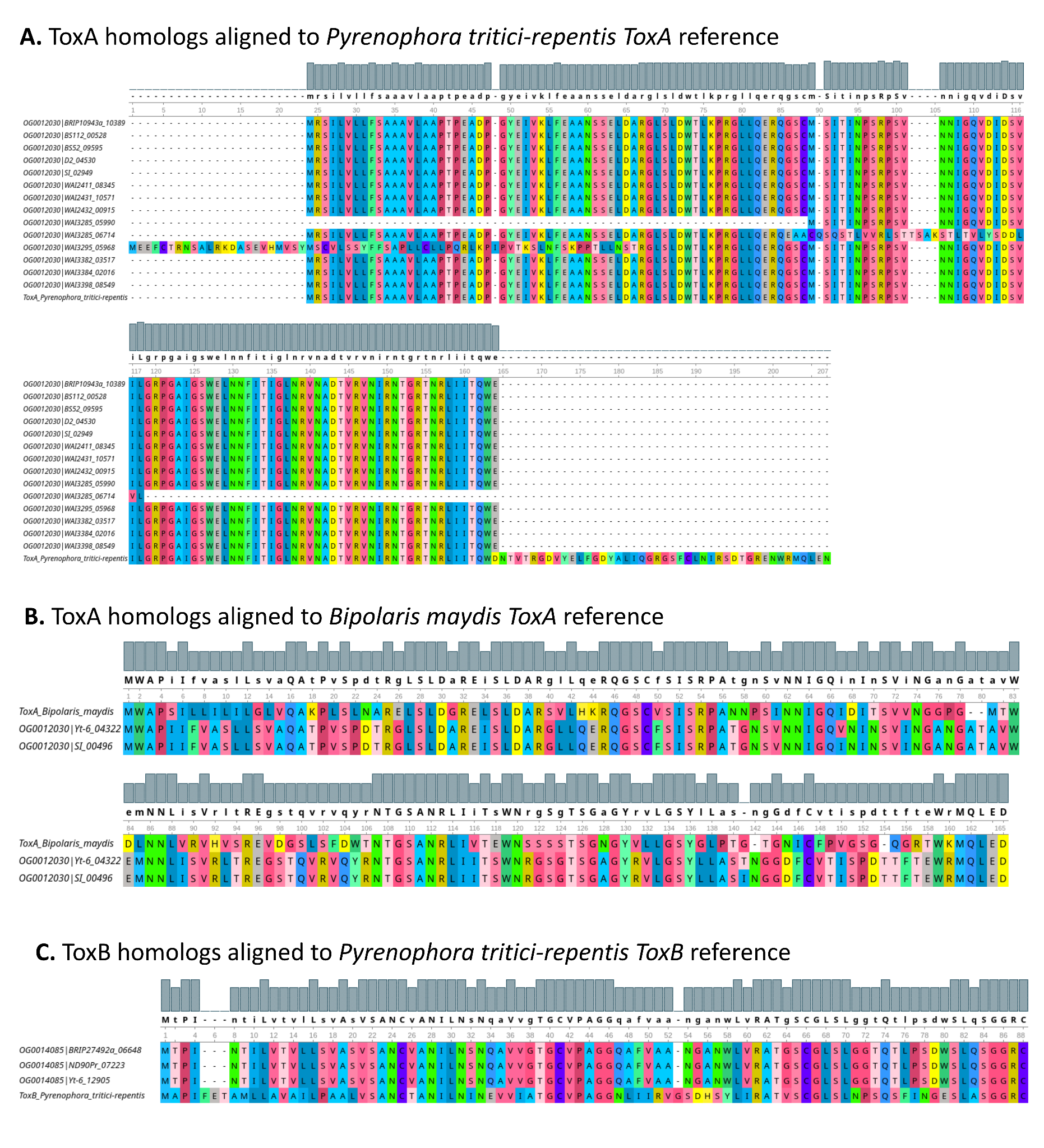

**Figure S15:** **Multiple sequence alignment of host-selective effector homologs identified in the *B. sorokiniana* pangenome.** (A) Alignment of OG0012030-encoded ToxA homologs against the ToxA reference sequence from Pyrenophora tritici-repentis, revealing highly conserved homologs with sequence identities reaching ~99%, consistent with strong conservation across multiple B. sorokiniana isolates. (B) Alignment of OG0012030 homologs against the ToxA sequence from Bipolaris maydis, showing substantially lower sequence similarity (~51%), indicative of a more divergent ToxA-like lineage. (C) Alignment of OG0014085-encoded ToxB homologs against the ToxB reference sequence from Pyrenophora tritici-repentis. ToxB homologs were detected in only three B. sorokiniana isolates and exhibited moderate sequence similarity (~52%), suggesting a restricted distribution and greater evolutionary divergence compared with Ptr-like ToxA homologs. Conserved amino acid residues are highlighted according to sequence similarity, and gaps introduced during alignment are represented by dashes.

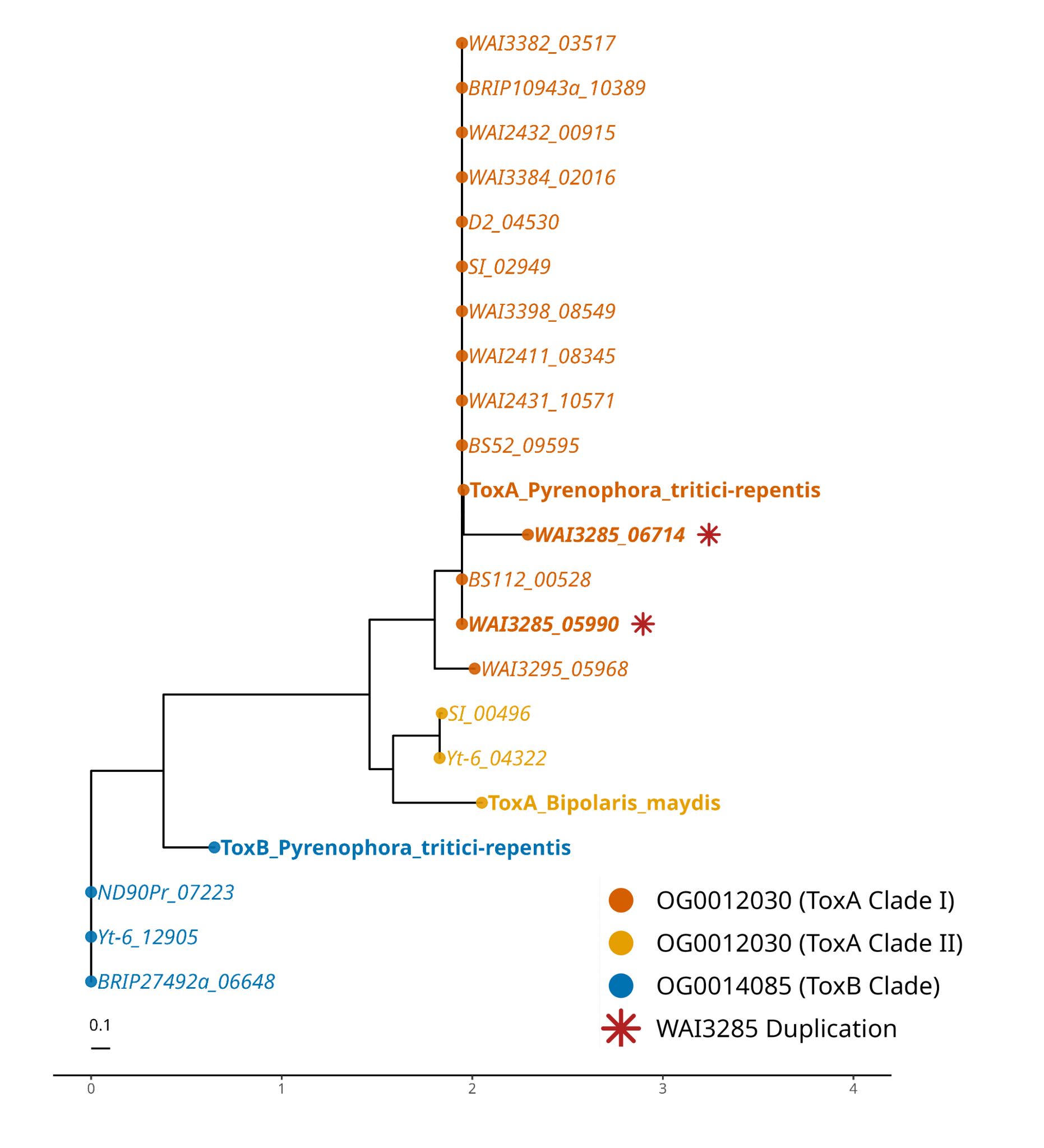

**Figure S16: Phylogenetic distribution and copy number variation of necrotrophic effectors ToxA and ToxB across pangenomic lineages of *B. sorokiniana*.** Maximum likelihood phylogenetic tree constructed using IQ-TREE (version 2.0.7) from aligned amino acid sequences of ToxA and ToxB homologs across fungal pathogen strains. Clades are color-coded based on pangenomic orthogroup annotations: OG0012030 (ToxA Clade I) is shown in vermillion, OG0012030 (ToxA Clade II) in orange/yellow, and OG0014085 (ToxB Clade) in ocean blue. Core reference strains (*Pyrenophora tritici-repentis* and *Bipolaris maydis*) are highlighted in bold. Duplicated copy of ToxA in strain WAI3285 is highlighted in bold-italic and annotated with a red asterisk (∗), indicating a structural duplication event containing two distinct paralogous copies of the *ToxA* locus (WAI3285_05990 and WAI3285_06714)

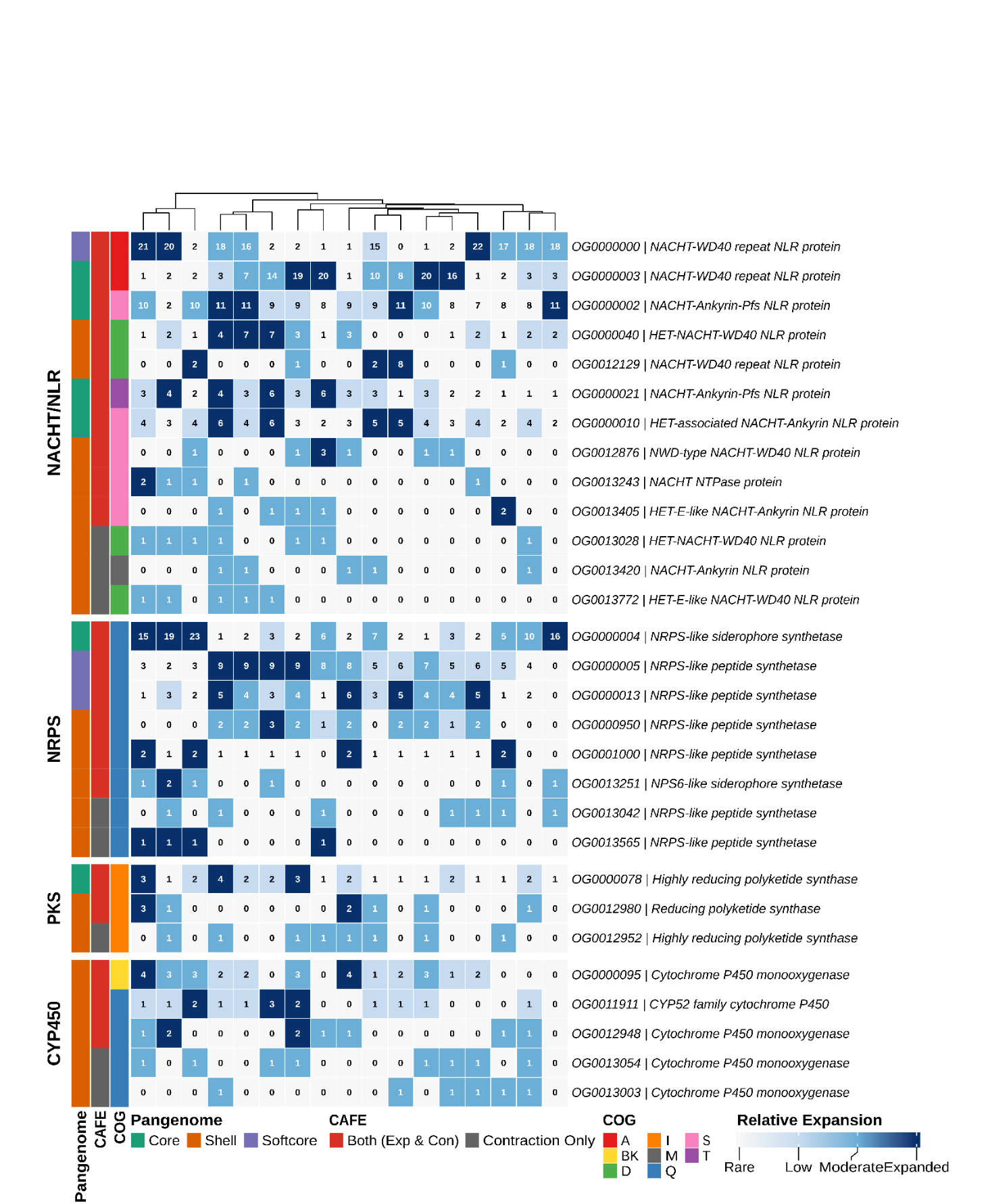

**Figure S17: Gene family expansion and contraction dynamics of pathogenicity- and defense-associated gene repertoires across 19 *B. sorokiniana* genomes.** Heatmap showing copy-number variation for 30 significantly evolving orthogroups identified by CAFE analysis. Orthogroups were grouped into four biologically relevant functional classes: NACHT/NLR-associated defense proteins (13 orthogroups), non-ribosomal peptide synthetases (NRPS; 8 orthogroups), polyketide synthases (PKS; 3 orthogroups), and cytochrome P450 monooxygenases (CYP450; 6 orthogroups). Cell values represent actual gene copy numbers in each genome, while color intensity indicates relative abundance categories derived from within-orthogroup quantile normalization (Rare, Low, Moderate, and Expanded), enabling comparison of expansion patterns among strains despite differences in absolute copy numbers. Genomes were hierarchically clustered using Ward’s method based on orthogroup copy number profiles. Left-side annotation tracks indicate pangenome classification (Core, Softcore, and Shell), CAFE evolutionary dynamics (Both Expansion and Contraction or Contraction Only), and eggNOG COG functional categories. **COG category abbreviations:** A, RNA processing and modification; D, cell cycle control, cell division and chromosome partitioning; I, lipid transport and metabolism; K, transcription; M, cell wall/membrane/envelope biogenesis; Q, secondary metabolite biosynthesis, transport and catabolism; S, function unknown; T, signal transduction mechanisms; B, Chromatin structure and dynamics.
